# The transglutaminase 2 interactome in HUVECs suggests its participation in an RNA-binding protein network

**DOI:** 10.1101/2025.08.26.672361

**Authors:** Bianka Csaholczi, Zsuzsa Csobán-Szabó, Károly Jambrovics, Ilma Rita Korponay-Szabó, László Fésüs, Róbert Király

## Abstract

Human transglutaminase 2 (TG2) is a multifunctional protein that exhibits various protein-modifying catalytic and protein-protein interaction properties. HUVECs express high levels of TG2, and identification of its interacting partners may provide mechanistic insight into its role in endothelial functions such as adhesion, migration, angiogenesis, and NO homeostasis. For this purpose, two approaches were employed. Firstly, bacterially expressed, site-specifically biotinylated recombinant TG2 was mixed with HUVEC extract to isolate interacting partners by affinity chromatography. Secondly, endogenous TG2 was silenced, and to ensure high antibody affinity, a triple Flag-tagged transgenic TG2-expressing HUVEC line was created, allowing stable binding to anti-Flag antibody-coated agarose for isolating TG2-associated protein complexes. Altogether, 170 and 356 TG2-associated proteins were identified, respectively, with 86 proteins overlapping between the two approaches.

The most enriched GO Molecular Functions of cellularly assembled TG2-interacting proteins confirm TG2’s involvement in cell-cell and cell-extracellular matrix communication, adhesion, cytoskeleton organisation, and exosome secretion. Stabilising the catalytically inactive closed TG2 conformation reduced the number of TG2 interactors, whereas stabilising its open form increased the number of associated membrane transporters.

Significant enrichment of RNA-binding proteins associated with TG2 was observed in all experiments, and a 42% overlap between the TG2 interactome and previously identified RNA-binding proteins in HUVEC was noted. Considering the recently recognised RNA-binding ability of TG2, our results suggest that TG2 participates in post-transcriptional regulations as a central hub within the network of RNA-binding proteins.

**GRAPHICAL ABSTRACT:** Cellularly assembled TG2-associated proteins were isolated and identified from endogenous TG2-silenced (eTG2-KD) and transgenic, high-affinity antibody, triple Flag-tagged TG2 (3F-TG2) expressing HUVECs under untreated conditions and after NC9 or GTPγS inhibitor treatments, which stabilise its open or closed conformation, respectively. TG2 interactome analysis suggests that it acts as a central hub in an RNA-binding protein (RBP) network.

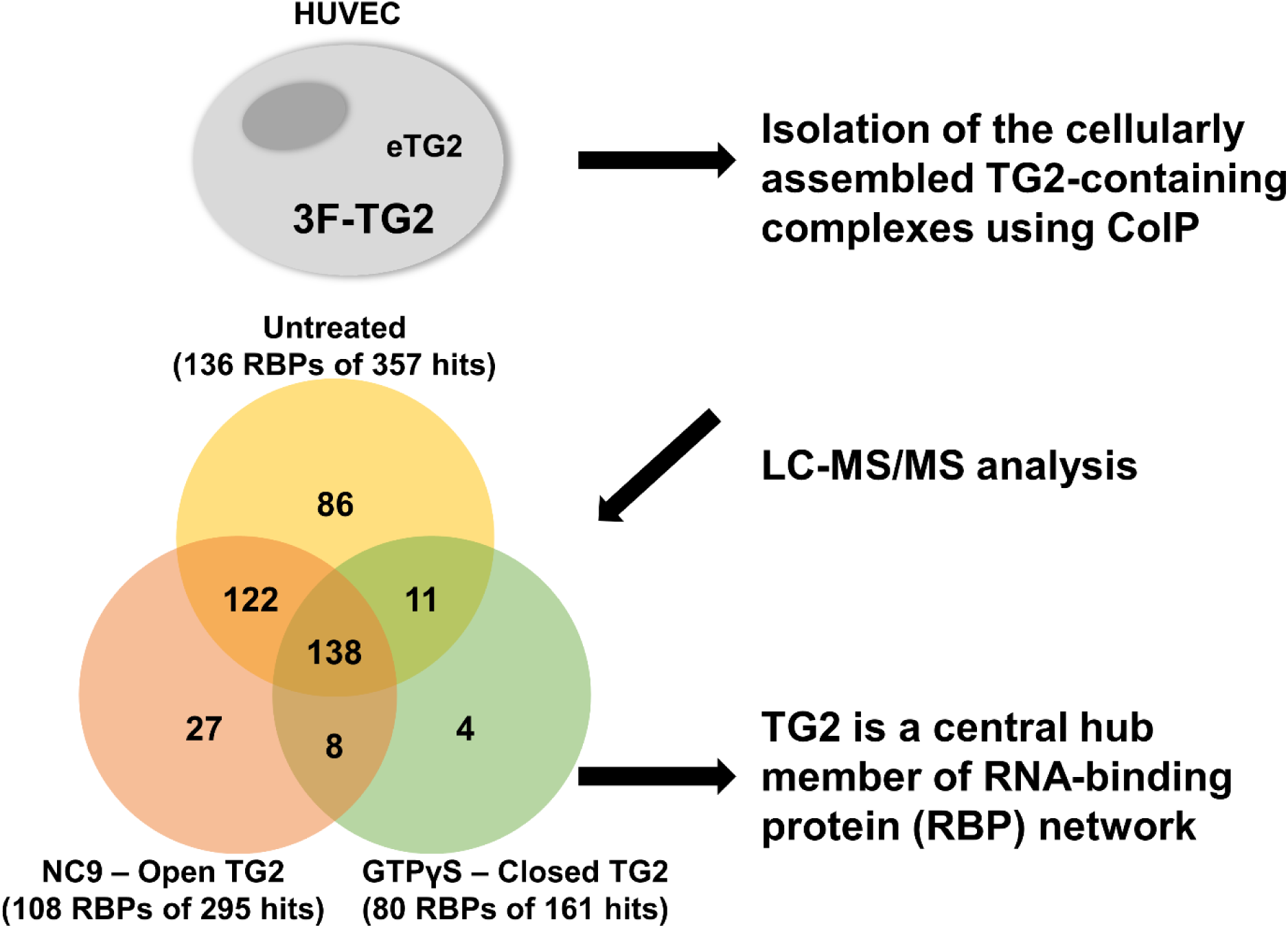

## Introduction

Mammalian transglutaminase 2 (tissue transglutaminase, TG2) is a unique multifunctional protein that manages a range of calcium-dependent posttranslational modifications, including protein crosslinking, amine incorporation, deamidation, isopeptide cleavage, and protein-disulfide isomerase activity. These functions are linked to its open conformation, while its protein kinase, GTPase, and ATPase activities are associated with its nucleotide-stabilised closed conformation (Ismaa et al., 1997). Furthermore, TG2 plays an anchoring role in the assembly of protein complexes involved in physiological processes such as focal adhesion, phagocytosis and the organisation of the extracellular matrix, as well as in the formation of a signalosome during pathological processes like acute promyelocytic leukaemia (Eckert et al., 2014; Jambrovics et al., 2023). Recently, our group published its novel biochemical feature, its RNA-binding ability, which is related to the open conformation of TG2 (Csaholczi et al., 2025).

TG2 is ubiquitously present in human tissues, particularly in the blood vessels, where endothelial cells, smooth muscle cells, and fibroblasts express TG2 (Nurminskaya & Belkin, 2012). In vascular function, endothelial cells play a crucial role in maintaining the balance between healthy and diseased states. Evidence supports the involvement of TG2 in endothelial cells’ adhesion and angiogenesis, inflammatory responses, regulation of blood pressure and vascular stiffness; however, mechanistic details have remained unclear. In rodents, age-related decrease in nitrosylation of TG2 has reciprocal correlation with vascular stiffening (Santhanam et al., 2010; Lai et al., 2017), and a transglutaminase activity inhibitor, which stabilises the closed form of TG2, can prevent the development of age-related hypertension (Pinilla et al., 2020).

Human endothelial cells express a high level of TG2, providing a model for characterising physiological functions of TG2 (Thomázy&Fésüs, 1989). On their surface, sufficiently nitrosylated TG2 can prevent the unwanted anchoring of neutrophils. Extracellular TG2 is essential for VEGF-mediated angiogenesis, which can be hindered by cell-impermeable TG2 inhibitors (Wang et al., 2013). TG2 may facilitate TGF-β1 signalling, promoting angiogenesis and endothelial-mesenchymal transition, thereby contributing to tissue remodelling during wound healing (Wang et al., 2017). Experiments silencing endogenous TG2 in human umbilical cord vein endothelial cells (HUVECs) and the consequent redistribution of extracellularly added recombinant human TG2 demonstrated its role in endothelial cell adhesion (Nadalutti et al., 2011). The absence of intracellular TG2, even in the presence of extracellular recombinant TG2, hindered cell survival and proliferation, resulting in a high ratio of apoptotic HUVECs (Nadalutti et al., 2011).

In recent years, studies employing proteomic approaches have illuminated some previously unrevealed aspects of TG2. Furini and colleagues found using a mouse unilateral ureteric obstruction (UUO) model, that exosomal transport of TG2 plays a role in the development of fibrosis leading to chronic kidney disease (Furini et al., 2018). By investigating the TG2 interactome in a mouse Alzheimer’s disease model, Wilhelmus and colleagues described TG2’s involvement in the dysregulation of synaptic transmission and cell adhesion in the brain (Wilhelmus et al., 2022). However, no comprehensive study has been conducted on human TG2-associated proteins or protein complexes in human endothelial cells under stress-free conditions.

In our study, after assembling a TG2 interactome using co-precipitation with human recombinant TG2 in HUVEC cell lysates, we developed and characterised an endogenous TG2-silenced, high antibody affinity, triple Flag-tagged transgenic TG2 expressing human endothelial cell model to reveal cellularly assembled TG2-containing protein complexes and provide insights into the physiological functions of TG2 in HUVEC. The results confirm the role of human TG2 in cell adhesion, cytoskeletal matrix modification, cell proliferation, and cell metabolism, while they also point to its possible functions in post-transcriptional regulation as a hub in the RNA-binding protein network in endothelial cells.

## Material and methods

All materials were purchased from Sigma-Aldrich-Merck (Munich, Germany) unless otherwise indicated.

### Cell cultures

The isolation, immortalisation, and culturing of HUVECs used in this study were described in detail previously (Shaw et al., 2021). In brief, immortalised HUVEC line was generated from endothelial cells isolated from the human umbilical cord vein of a healthy newborn by viral delivery of telomerase gene, and cultured in M199 medium (Biosera, Nuaille, France) containing 10% FBS (Thermo Fisher Scientific), 10% EGM2 Endothelial Growth Medium (Lonza, Basel, Switzerland), 20 mM HEPES (Biosera), 100 U/mL Penicillin, 100 μg/mL Streptomycin and 2.5 μg/mL Amphotericin B (Biosera). Immortalised cells completely retain the morphological properties of primary endothelial cells and express endothelial markers (Csaholczi et al. 2025).

### Preparation of cell extract

Cells were cultured in T75 flasks (TPP) and detached using trypsin-EDTA treatment for 5 minutes at 37°C. Then, cells were washed twice with ice-cold PBS using centrifugation (1000 g for 10 min, at 4°C). Then, pelleted cells were resuspended in lysis buffer (25 mM Tris-HCl, pH 7.4, 150 mM NaCl, 1 mM EDTA, 5% glycerol, 0.5% NP-40, 1 mM PMSF), including PIC (Protease Inhibitor Cocktail, Sigma) in a 100-fold dilution, and PhosSTOP (Phosphatase Inhibitor Cocktail Tablets, Roche) according to the manufacturer’s instructions. The cell suspensions were sonicated using five impulses, and after spin centrifugation (to remove cell debris), the clear supernatant was used as cell extract for further experiments. The protein concentration was measured using Bio-Rad Protein Assay Dye Reagent Concentrate (Bio-Rad, Hercules, CA, USA).

### Preparation of site-specifically biotinylated human recombinant TG2

To produce recombinant human TG2 biotinylated site-specifically at the N-terminus (N-BAP-rhTG2), a biotin acceptor peptide sequence (BAP, GLNDIFEAQKIEWHE) was inserted into pET-30 Ek/LIC-TG2 bacterial expression vector (Kiraly et al., 2009), exchanging the thrombin cleavage site at the N-terminus using N-BAPINSfor and N-BAPINSrev primers (Supplementary Table S14) according to Q5 Site-Directed Mutagenesis Kit Protocol (NEB, insertion mutagenesis). The vector was checked by restriction digestion and Sanger sequencing (Eurofins Genomics, Ebersberg, Germany). Then, the site-specifically biotinylated N-BAP-biotin rhTG2 protein was expressed in *E. coli Rosetta 2 (DE3)* (Merck) bacterial cells upon induction with 0.1 mM isopropyl β-d-1-thiogalactopyranoside (IPTG) and purified using His GraviTrap columns as described previously (according to the improved protocol in Kiraly et al., 2016). The Supplementary information contains the coding DNA and its translated protein sequence (S15).

### Affinity purification of TG2-associated proteins using site-specifically biotinylated recombinant TG2

First, 250 µl Pierce High Capacity Neutravidin Agarose resin (Thermo Scientific) was equilibrated with 1 times MQ water, then using 3 times R buffer (20 mM Tris, 150 mM NaCl, 1 mM EDTA, 1 mM imidazole pH 7.8-8) using a batch procedure (centrifugation with 2500 g for 2 min, 4 °C). For immobilisation, 100 µg site-specifically biotinylated N-BAP-biotin rhTG2 was added to 250 µl equilibrated resin in a total of 500 µl lysis buffer and was incubated for 2 hours at 4°C. Then, the resin was blocked with 300 µl of 333 µg/ml biotin in R buffer for 1 hour at 4 °C. The affinity purification was performed from 2.5-3 mg/ml HUVEC extract during continuous rotation overnight at 4 °C. The next morning, after three times washing with a CoIP buffer (25 mM Tris, 150 mM NaCl, 1 mM EDTA, 5 % glycerine, pH 7.4), interacting proteins were eluted using 0.1 M glycine-HCl buffer (pH 3.5). High-affinity interacting proteins were eluted from the resin, adding 6x Denaturation buffer (6xDB; 300 mM Tris-HCl, pH 6.8, 12 (m/v)% SDS, 20 (v/v)% glycerol and 0.43 M β-mercaptoethanol) and boiling the suspension at 100 °C for 10 min. Both eluted fractions were run into SDS-polyacrylamide gel (5% stacking gel, 10 % separation gel) till the top of the separation gel, and before separation, the whole Coomassie-blue-stained band from each sample was cut out and sent for LC-MS/MS analysis.

### Creation of an inducible endogenous TG2 silencing vector and a TG2 knocked-down HUVEC line

The human TG2 silencing oligonucleotide and a scramble sequence (TRCN0000272816, SCRAMBLE; Sigma-Aldrich, Supplementary Table S14) were cloned into the pLKO Tet-On puro vector, which was a gift from Dmitri Wiederschain (Addgene plasmid #21915; RRID:Addgene_21915), according to the provided protocol (Wiederschain et al., 2009). The cloning was confirmed by restriction digestion and Sanger sequencing (Eurofins Genomics).

For efficient transfer, the pseudo-virions were generated in HEK293FT cells (Thermo Fisher Scientific) using a calcium precipitation method with the pMD2.G and psPAX2 vectors (gift from Didier Trono; Addgene #12259 and #12260). The virion-containing supernatant was harvested twice every 24 hours and concentrated using Amicon Ultra-15 100 kDa concentrator tubes (approximately 20x; Merck-Sigma).

The previously immortalised HUVEC cells (by telomerase transduction; Shaw A et al. 2021 Front CDB) were infected at 70% confluence in 12-well plates (TPP) with a mixture of 5-times diluted concentrated pseudo-virions and HUVEC medium to prepare TG2-silenced (knocked-down, TG2-KD) and scramble control (SCR) cell lines. Transduced cells were selected using 4 µg/ml puromycin. Silencing was then induced with 4 µg/ml doxycycline. To ensure efficient silencing, selected cells were treated with 4 µg/ml of doxycycline for 3 weeks, with the medium changed every 48 hours.

### Preparation of the transgene TG2 expressing vector and its delivery into TG2-silenced HUVEC

Transgene human TG2, containing Val at position 224, was expressed in a triple N-terminal Flag-tagged form with an 18-mer linker sequence (MDYKDHDGDYKDHDIDYKDDDDK LGSMGTNSVDWIRYRIQT-TG2). To prepare this construct (Scheme is in Fig.3a), two additional Flag tags were cloned into the pENTR4-FLAG (w210-2) entry vector (gift from Eric Campeau & Paul Kaufman; Addgene #17423 (Campeau et al., 2009) prior to the existing Flag tag, using insertion mutagenesis in accordance with the Q5 Site-Directed Mutagenesis Kit Protocol (NEB; for primer sequences see Supplementary Table S14). Subsequently, human TG2 cDNA was subcloned into the EcoRI site from the pET-30 Ek/LIC-TG2 vector (Kiraly et al., 2009). The 3xFlag-TG2 cDNA from the constructed pENTR4-3xFlag-TG2 vector was cloned into the pCW57.1 destination vector (gift from David Root; Addgene #41393) using the Gateway LR Clonase II Enzyme mix (Invitrogen), following the manufacturer’s instructions. The all-in-one pCW57.1 destination vector contains the necessary genetic elements for a ”DOX-ON”, doxycycline-inducible transgene expression. Each cloning step was confirmed by Sanger sequencing, and the supplementary information includes the coding DNA and its translated protein sequence (S16). The generation of pseudo-virions and the infection were described in the previous session. The inducible transgene TG2-expressing and endogenous TG2-silenced cell line was selected using a higher puromycin concentration than the KD cell lines due to the increased copy number of the resistance factor (6 µg/ml). The silencing of endogenous TG2 and the production of transgenic tagged TG2 were induced with 4 µg/ml doxycycline.

### Co-immunoprecipitation of the cellularly assembled TG2-containing complexes

To identify the interactome of human TG2 in the developed 3xFlag-tagged transgenic TG2-expressing HUVEC model, an anti-Flag M2 affinity resin based on a batch protocol was employed. First, 300 µl of monoclonal anti-FLAG M2 resin suspension was equilibrated with the CoIP buffer (25 mM Tris, 150 mM NaCl, 1 mM EDTA, 5 % glycerine, pH 7.4). In each case, 3.3-3.5 mg of protein-containing cell lysate was incubated overnight with the anti-FLAG M2 resin. The nonspecific background binding to the resin was determined using mouse IgG agarose as a control. After washing three times, interacting proteins were eluted using 0.1 M glycine-HCl buffer, pH 3.5. A volume of 55 μl of 6x denaturation buffer and heat denaturation (100°C, 10 min) were also applied to remove proteins bound with high affinity. Subsequently, the eluted samples were run on a 10% SDS polyacrylamide separation gel. When the proteins migrated from the 5% stacking gel to the separation gel, electrophoresis was stopped, and after Coomassie staining, the unseparated proteins were cut out and sent for MS analysis.

To reveal the interactome of open TG2, live cells were treated with 15 µM NC9 modulators 24 hours prior to lysis. While GTPγS cannot penetrate into the cells, it was added to the cell extract at a concentration of 1 mM and maintained during the CoIP.

### MS analysis

The protein-containing gel pieces in nanopure water were submitted to LC-MS/MS analysis at the Proteomics Core Facility of the University of Debrecen.

In a previous study, a similar LC-MS/MS analysis was conducted (Csobán-Szabó et al., 2021), in which the analysis method was detailed. Briefly, the proteins in the submitted gel pieces were digested with trypsin overnight (37 °C). After digestion, the samples were dried to a volume of 10 µl. The protein samples were prepared for MS analysis using the Easy nLC1200 (Thermo Scientific) nanouplc-Orbitrap Fusion (Thermo Scientific) MS/MS system. The results were imported into the Scaffold 5.0.1 software.

In the Scaffold software, the Exclusive Unique Peptide Count display option and a protein threshold of 1.0% false discovery rate (FDR), along with a peptide threshold of 0.1% FDR, were applied, requiring a minimum of one identified peptide for each protein. The fragmentation table of each detected unique peptide was meticulously checked to verify that the detected peptide is a genuine hit. For this verification, the fragmentation table must include at least four B or Y ion series. The experiments were repeated at least 3 times, and the hits obtained from each repetition were aggregated, after which the control non-specifically bound protein hits were deducted.

### RNA isolation and Q-PCR measurement

RNA isolation and Q-PCR assay were conducted following a previously detailed procedure (Jambrovics et al., 2020) utilising TRI Reagent (cat #TR 118; Molecular Research Centre, Cincinnati, OH, USA). The total RNA concentration and quality were assessed using the Nanodrop2000 Spectrophotometer (Thermo Fisher, Waltham, MA, USA 02451). The samples were diluted to a concentration of 200 ng/µl, and RNA was reverse transcribed into cDNA with the High Capacity cDNA Reverse Transcription Kit (Applied Biosystems), in accordance with the manufacturer’s instructions. Transcript levels were evaluated by real-time Q-PCR employing TaqMan probes (TG1 #Hs_01070310_m1, TG2 #Hs_01096681_m1, TG3 #Hs_00162752_m1, TG4 #Hs_00162710_m1, TG5 #Hs_00909973_m1, TG6 #Hs_00975389_m1, TG7 #Hs_0036497_m1, F13A1 #Hs_00173388_m1, Actin #Hs_01060665_g1 Actb). Real-time monitoring was performed using a Roche LightCycler® 480 II instrument (white Roche LightCycler® 480 Multiwell Plate 384). Transcript levels were normalised to the level of actin B, and gene expression was determined by the C_T_ method.

### Western blot

Proteins were separated by 10% SDS-PAGE and then transferred to PVDF membrane (Millipore Immobilon-P membrane, 0.45 µm) using semidry blotting (Bio-Rad Trans-Blot SD Semi-Dry transfer cell device). 5 (m/v) % low-fat milk powder containing TTBS was used for blocking. Primary and secondary antibodies were diluted in 0.5 (m/v) % low-fat milk powder in TTBS, and TTBS was used for washing four times for 15 minutes. The following antibodies were used in the experiments: monoclonal transglutaminase-2 antibody (CUB7402, ThermoFisher #MA5-12736, 1/10000), actin antibody (Sigma #A2066, 1/3000), anti-FLAG M2 antibody (Sigma #F1804, 1/2000), goat anti-mouse IgG (Advansta #R-05071-500, 1/10000) and goat anti-rabbit IgG (Advansta #R-05072-500, 1/10000).

### Adhesion assay

The cell adhesion assay was conducted according to an earlier description (Shaw et al., 2021). Briefly, the cells were incubated for 24 hours in a starvation medium (1% FBS, no EGM2). On a fibronectin-coated 96-well plate, the adherence of 1000 cells was tested during a 1-hour incubation using Cell-Titer Blue reagent (Promega). The increasing fluorescence intensity values were monitored and read every 2 hours (Ex/Em: 530/590 nm; BioTek Synergy HT instrument), and their slope values (RFU/min) were determined and utilised for comparison after subtracting the slope value obtained from the wells containing only medium. The data are presented as a percentage of unmodified immortalised HUVECs.

### Labelling Annexin-V and Sorting Live/Dead Cells

The labelling and analysis were conducted following an earlier published protocol (Jambrovics et al., 2023). Briefly, cells were collected, washed with 1× PBS, and centrifuged at 700 rpm for 3 minutes at 4 °C. The cells were labelled with FITC-conjugated Annexin-V (Biolegend, San Diego, CA, USA) and propidium iodide (PI, Biolegend) for 15 minutes in the dark. The cells were analysed using a BD FACSAria™ III flow cytometer (B.D. Biosciences, San Jose, CA, USA). HUVEC cells were gated based on the size and granularity, followed by the determination of the Annexin-V and PI features. Data and results were validated by Flowing software.

### Bioinformatics analyses

The detected potential interaction partners of TG2 were analysed using the PANTHER 19.0 (pantherdb.org) and the STRING v12 (string-db.org) databases. Based on the false discovery rate values, the most enriched Gene Ontology (GO) Molecular Functions are listed and represented using GraphPad Prism version 8.0.1 (Dotmatics). Statistical analysis was performed using GraphPad Prism version 8.0.1.

## Results

### Identification of TG2 interacting partners in HUVECs extracts using recombinant protein supports its function in cell adhesion, cytoskeletal reorganisation, and signalling processes, and highlights its potential contribution to RNA binding complexes

In order to reveal the TG2 interactome in HUVECs, a biotin-neutravidin agarose based affinity chromatography was applied first. To prevent random biotinylation of the bait protein and subsequent steric effects on the interactome, a biotin acceptor peptide (BAP) was cloned into the N-terminal overhang of the recombinant TG2 coding vector. The Ni-NTA affinity-purified recombinant human TG2 protein (N-BAP-biotin rhTG2), expressed in *E. coli Rosetta 2 (DE3)* strain and biotinylated during its expression in the bacterial cells, was applied for the affinity chromatography from HUVEC extract, which was lysed by an NP-40 containing buffer to allow the formation of low-binding affinity protein complexes. Potential interacting partners were identified using LC-MS/MS analysis. Proteins, detected in control (Supplementary Table S1), were subtracted from the sample hits to close out the interactome of the agarose resin (see experimental scheme in Fig. 1).

**Figure 1.**
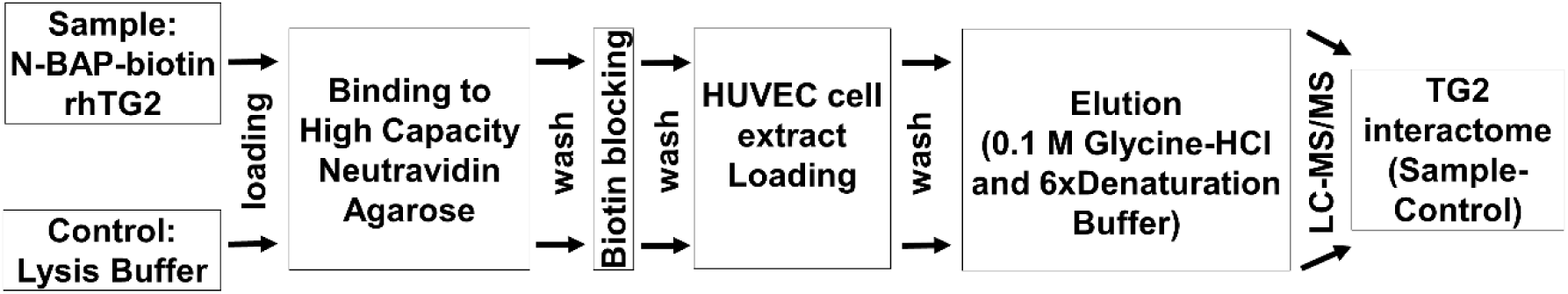
Experimental scheme to identify the interactome of bacterially produced site-specifically biotinylated recombinant human TG2 (N-BAP-biotin rhTG2) from HUVEC extract.

After control subtraction, whose list contained some already known TG2 interactors (e. g. CALM1, CALR, TLN1, TUBA/B; Supplementary table S1), 170 TG2-associated proteins were identified, including 11 already known TG2-interacting proteins (6.5 % of all hits; COPA, COPB1, CYB5R3, DNAJA1, FN1, MMP1, PRDX1, TUBB3, TUBB4A, ZYX, VCL based on the TRANSDAB database), supporting the reliability of our approach (Supplementary Table S2). To visualise the network of TG2 interacting partners, the most abundant protein hits were filtered based on the number of detected unique peptides, with a minimum of five, resulting in 26 protein hits (15.3 % of all hits; Fig. 2a). Based on STRING analysis, the protein network shows more connections than expected (30 edges instead of the expected 18), suggesting that these proteins are biologically connected. They do not show functionally enriched GO Molecular Functions, but the most enriched ‘Cellular Compartments’ are ‘Focal adhesion’, ‘Extracellular exosome’, ‘Vesicle’, ‘Side of membrane’, ‘Extracellular space’ (Supplementary Table S3), which supports the presence of TG2 in exosomal vesicles involved in the externalisation of TG2. The enriched membrane proximity of TG2-associated proteins correlates with the contribution of TG2 to signalling or transport processes. The MCAM protein contributes to the cohesion of the endothelial monolayer. Several interactors play a role in cell-cell interactions, cell-extracellular matrix links, and cytoskeletal rearrangements (FLII, CDC42BPS, DYNC1H1, PLEC, FN1, FMNL2), or consist of either membrane or vesicular proteins (CD59, NT5E, COPB1, AP2A1). Additionally, some of these TG2 interactors are transport regulatory proteins (GNB1, GNB2, GNAI2), or are involved in cell cycle control and DNA repair (TMPO, PARP1, SSRP1, XRCC5, RALA).

**Figure 2.**
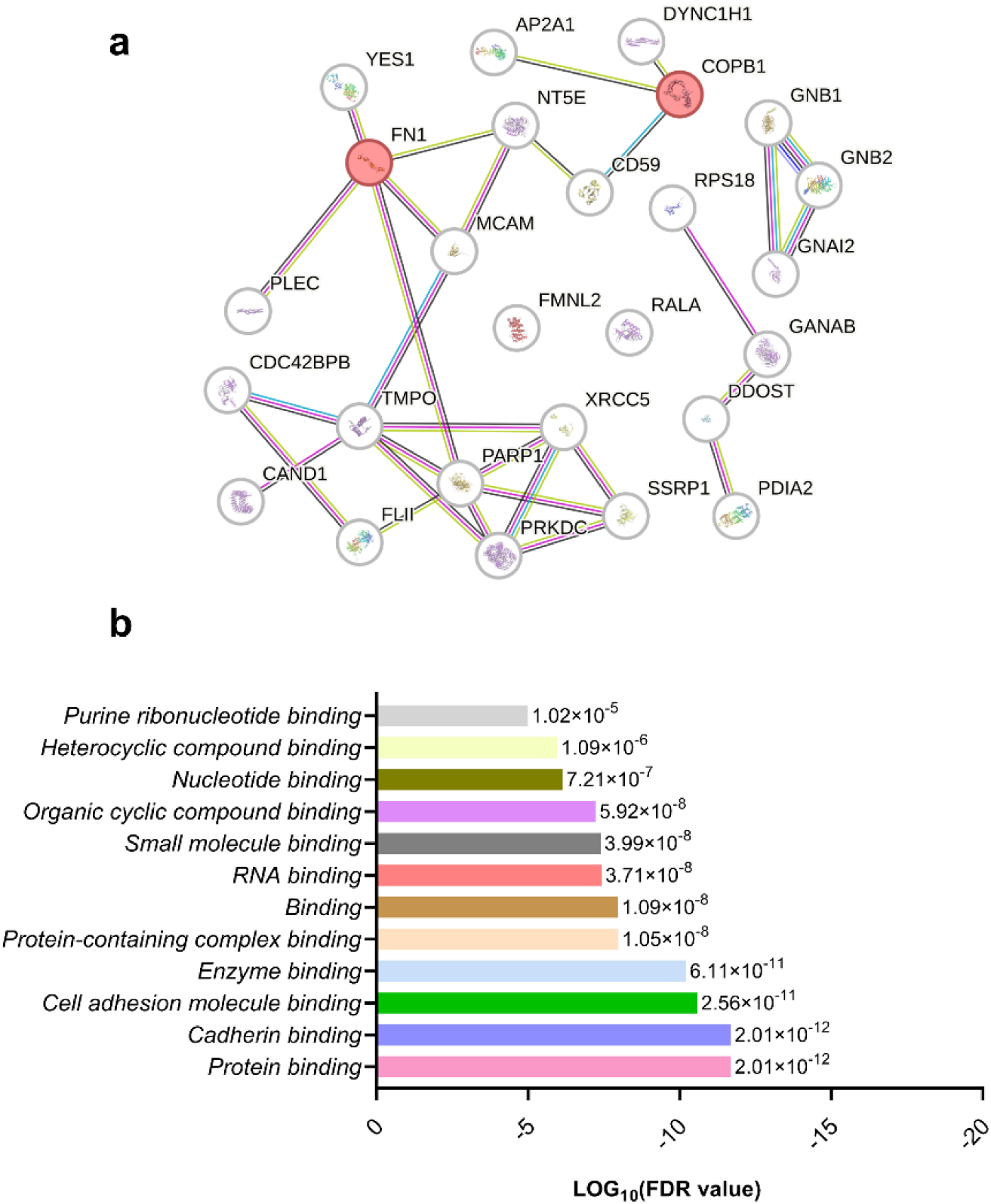
The protein interaction network of the most abundant TG2-associated proteins and GO Molecular Functions of the identified interactors of biotinylated recombinant TG2. (a) The protein interaction network of the most abundant TG2-associated partners was mapped against the Homo sapiens reference database using String v12.0, encompassing both known and predicted interactions, with the default threshold confidence level set at 0.4 (https://string-db.org/, obtained on 6 June 2025). Hits for TG2 interacting partners detected with unique peptides more than five times are deemed the most abundant. The red nodes label the already known TG2 interactors. (b) The column diagram illustrates the most enriched GO Molecular Functions of TG2-associated proteins, based on the false discovery rate (FDR), analysed by the STRING database.

The analysis of the entire list of N-BAP-biotin rhTG2-associated proteins led to similar results; the most enriched GO Molecular functions (Supplementary Table S4; Fig. 2b) include ‘Cadherin binding’ and ‘Cell adhesion molecule binding’, which are expected processes given TG2’s involvement in cell adhesion and migration. However, enrichment in ‘RNA binding’ is a novel observation in the context of TG2-associated proteins.

### Development and characterisation of Tet-inducible endogenous TG2-silenced and triple Flag-tagged transgene TG2 expressing HUVEC line

To mitigate the potential drawbacks of affinity purification with a bacterially expressed protein and to benefit from a model in which the bait TG2 protein is expressed in the target HUVECs, we decided to develop a novel model system for identifying TG2-associated proteins by separating the TG2-containing protein complexes following their assembly in living cells, utilising affinity tag to enhance binding and reduce possible steric inhibition of binding partners on antibody-affinity tag interaction. A novel HUVEC cell line was developed in which the endogenous TG2 was downregulated by an shRNA (TG2-KD), and triple Flag-tagged transgene TG2 (3F-TG2) was expressed for binding and subsequent detection of TG2-containing protein complexes formed under native conditions (Fig. 3a, scheme). The silencing short hairpin sequence specifically targeted the untranslated regions (UTR) of endogenous TG2 mRNA, and the expression of a transgene with different UTR regions was not affected by the silencing. The potential negative consequences of a lack or low level of TG2 protein in HUVEC were mitigated by inducible silencing, and both silencing and transgene expression were achieved simultaneously by doxycycline-dependent (DOX-ON) treatment. Since endothelial cells exhibit low transfection efficiency, lentiviral vectors were applied.

**Figure 3.**
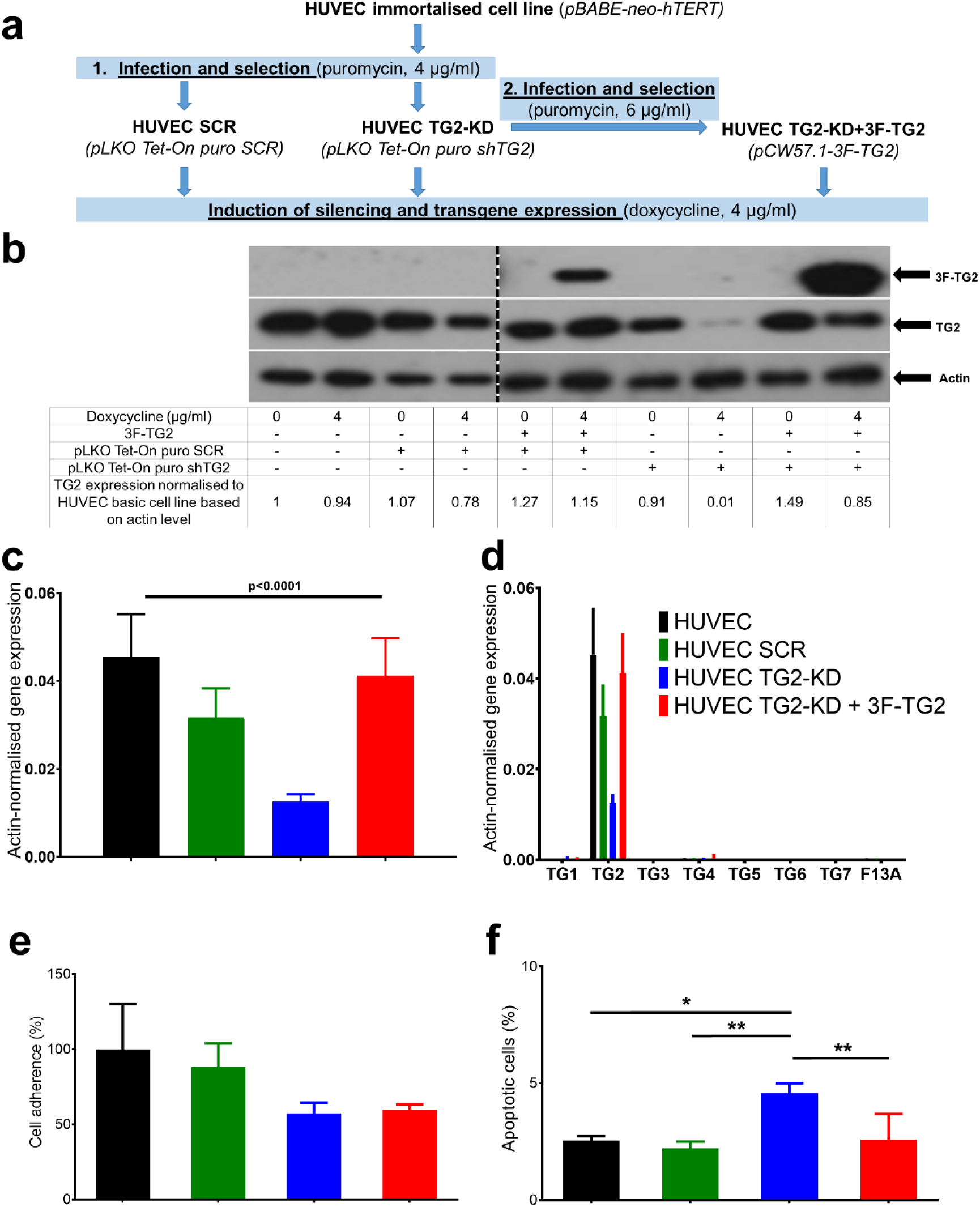
Generation and characterisation of the Tet-inducible 3xFlag-TG2 (3F-TG2) transgene overexpressing and endogenous TG2 silenced HUVEC line. (a) Scheme of the cell line development. The pLKO Tet-On puro shTG2 vector was introduced into the immortalised HUVEC cell line via lentiviral gene transfer, resulting in a HUVEC TG2-KD cell line. After confirming successful silencing, another infection was used to deliver the pCW57.1-3F-TG2 vector coding for transgenic 3F-TG2 (HUVEC TG2-KD+3F-TG2). (b) Monitoring of endogenous TG2 silencing and 3F-TG2 transgene expression by Western blot. Upon doxycycline induction, the TG2 protein levels were assessed with either anti-TG2 (CUB7402) or anti-Flag M2 antibodies. The TG2 expression level was normalised to beta-actin. Representative image (n=2). (c) TG2 mRNA level in the developed cell lines, measured by real-time qPCR. Beta-actin was used as an internal control. (d) Effect of TG2 silencing on the expression level of transglutaminase enzyme family members. The expression levels of human transglutaminases were normalised to beta-actin. (e) Effect of endogenous TG2 silencing and transgene 3F-TG2 expression on HUVEC cell adhesion. Data (mean±SD, n=3) are presented as the percentage of fluorescence change of the adhered immortalised HUVECs. (f) Effect of endogenous TG2 silencing and its subsequent compensation by 3F-TG2 transgene expression on the apoptotic cell ratio. Cells were labelled using Annexin-V and propidium iodide (PI). 10^4 cells were counted (One-way ANOVA Tukey post-hoc test; *p<0.05, **p<0.01).

First, endogenous TG2 expression was silenced by using lentiviral transduction to deliver the Tet-On puro shTG2 vector (targeting TG2 mRNA 3’UTR) into the immortalised HUVEC cell line. After three weeks of doxycycline induction, the reduced TG2 level in the puromycin selected TG2-knockdown HUVEC cell line (HUVEC TG2-KD) was confirmed at both RNA and protein levels (Fig. 3b, c). Doxycycline induction did not alter the level of TG2 protein in control cells, while the shTG2 sequence significantly reduced the amount of TG2 protein in TG2-KD HUVEC compared to the unmodified HUVEC and the scramble sequence containing HUVEC (HUVEC SCR) cell lines.

Next, the 3F-TG2 coding Tet-inducible vector was introduced into the TG2-KD HUVEC line (HUVEC TG2-KD+3F-TG2) through a second lentiviral gene transfer. Unfortunately, both Tet-inducible constructs contained puromycin resistance; however, following the second transduction, an increased level of puromycin was applied to enhance selection. After three weeks of doxycycline induction, the RT-qPCR results and Western blots demonstrated that the delivery of transgenic 3F-TG2 restored the TG2 level at both the RNA and protein levels in the developed HUVEC TG2-KD+3F-TG2 cell line (Fig. 3b, c). The expression levels of other transglutaminase family members were unaffected by silencing and transgene TG2 expression (Fig. 3d).

To functionally characterise the developed cell lines, cell adhesion properties on fibronectin surface, a TG2-dependent endothelial ability, were first compared. It is known that silencing TG2 with siRNA or inhibiting TG2 activity negatively affects cell adhesion on a fibronectin surface (Nadaluti et al. 2011) TG2-KD HUVEC demonstrated lower cell adhesion properties than the unmodified HUVEC and HUVEC SCR cells (Fig. 3e), although this difference was not statistically significant. The delivery of 3F-TG2 was unable to compensate for the observed decrease, possibly due to the relatively large hydrophobic expression tag of the protein.

It is also known that silencing TG2 increases the proportion of apoptotic HUVEC cells (Nadaluti et al. 2011). Our findings align with this earlier observation since TG2-KD HUVEC cultures have a higher ratio of apoptotic cells as compared to the control and 3F-TG2-compensated cell lines (Fig. 3f), implying that the transgene TG2 may compensate for the effects of endogenous TG2 silencing.

### Identification of TG2-associated protein complexes assembled in HUVECs using 3xFlag-tagged transgenic TG2

In order to reveal the cellularly assembled TG2 interactome, we used the extract from endogenous TG2-silenced and 3F-TG2 expressing HUVEC (the experimental scheme is shown in Fig. 4a). To prevent any steric masking effects between interacting partners and anti-TG2 antibodies, which could decrease the efficiency of co-immunoprecipitation by competing for the same binding site, we isolated TG2-containing protein complexes using anti-FLAG M2 agarose resin. This resin binds the triple Flag tag overhang on TG2 with a much higher affinity than a simple Flag tag or typical antibody-antigen interactions (Ranawakage et al., 2019). As a control, we applied mouse IgG agarose resin to identify non-specifically bound proteins from the HUVEC extract to the resin or constant chains of the mouse IgG antibody. After LC-MS/MS analysis and control (Supplementary Table S5) subtraction, which contained fewer non-specifically bound proteins compared to the neutravidin agarose, we identified 356 TG2-associated proteins (Supplementary Table S6), with 86 overlapping hits recognised by both the affinity chromatography and this approach (Supplementary Table S7). A total of 114 proteins (32% of hits) were detected more than five times by unique peptides, which are regarded as the most abundant interactors (Fig. 4b). The full list includes 19 (5.3%) already known TG2 interacting partners: TLN1, TUBA1A, H2BC4, COPG1, YWHAZ, COPB2, CALR, CD44, OSBPL3, ITGA2, ITGA6, ITGB1, MYOF, TUBA1C, UGGT1, YWHAK, APMAP, CANX, TUBB4A (TRANSDAB, Kanchan et al., 2015). Interestingly, the proportion of already known interaction partners is similar among the barely detected and the most abundant hits (app. 5 %), supporting the relevance of protein hits with low unique peptide numbers. The number of protein hits indicates that the composition of TG2-associated protein complexes may be influenced by the emergence of different structural elements during TG2 protein folding and secondary interactions through primary partners. (Brown et al., 2025).

**Figure 4.**
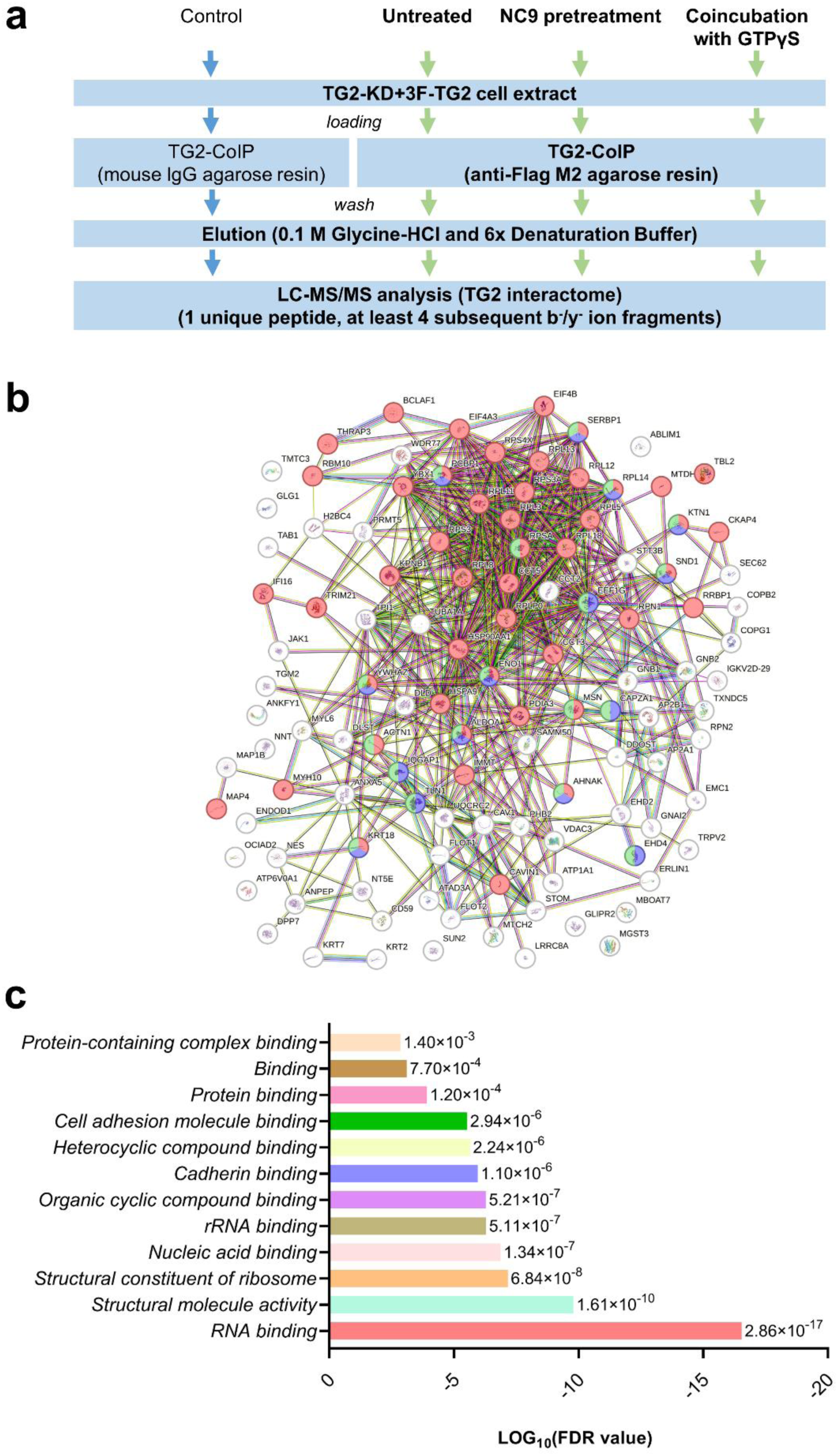
Workflow of the coimmunoprecipitation, the interaction network and the most enriched GO Molecular functions of the most abundant 3F-TG2-associated proteins, isolated by anti-Flag M2 IgG agarose resin-based coimmunoprecipitation. (a) To identify the interactome of open or closed 3F-TG2, cells were pretreated with NC9 inhibitor for 24 hours, or the cell lysate was co-incubated with GTPγS during the lysis and coimmunoprecipitation, respectively. After elution, the samples were sent for LC-MS/MS analysis (n=3). (b) The bioinformatic analysis from protein interaction network was mapped against the Homo Sapiens reference database using String V12.0 (https://string-db.org/, 6 June 2025). The map includes the known and predicted protein interactions of the protein hits, detected with unique peptide counts exceeding five in LC-MS/MS analysis. (Colour code of the nodes: red=RNA binding; purple=Cadherin binding; green=Cell adhesion molecule binding.) (c) The column diagram shows the most enriched GO Molecular functions based on the false discovery rate from the STRING database.

### Molecular functions and cellular localisations of 3F-TG2-associated protein complexes

The STRING interaction network analysis of the most abundant TG2-associated proteins (Supplementary Table S8; Fig. 4c) demonstrated the enrichment of “Structural molecule activity”, “Cadherin binding”, and “Cell adhesion molecule binding” regarding GO Molecular Functions, supporting the contribution of TG2 to cell adhesion, migration, proliferation, cadherin binding, and cytoskeleton rearrangement. These are essential processes for maintaining the physiological functions of endothelial cells, such as adhesion, cell-cell junctions, and angiogenesis. Our novel approach reveals a more comprehensive list of proteins found in TG2-associated complexes that collaborate to carry out endothelial functions. Surprisingly, the most enriched GO Molecular Function was “RNA binding” for both the complete list and the more frequently detected interactors. The interactome analysis was repeated after excluding keratins (KRTs) and ribosomal proteins (RPLs and RPSs), which are typically the most abundant proteins in the cells, and were potentially identified as nonspecific interactors, but the result remained the same (data not shown). Given that TG2 was recently identified as an RNA-binding protein (Csaholczi et al, 2025), this suggests that TG2 may play a role in assembling RNA-protein complexes, either as a scaffolding or regulatory protein, or with a special function that remains unknown. The general scaffolding functions are further supported by the significant overlap between the previously identified RNA-binding proteins (Supplementary Table S9) and TG2-associated proteins (Figure 5). The Venn diagram indicates that of the identified 356 interactors, 150 (42%) have been previously reported as RNA-binding proteins from the HUVECs (Csaholczi et al., 2025).

**Figure 5.**
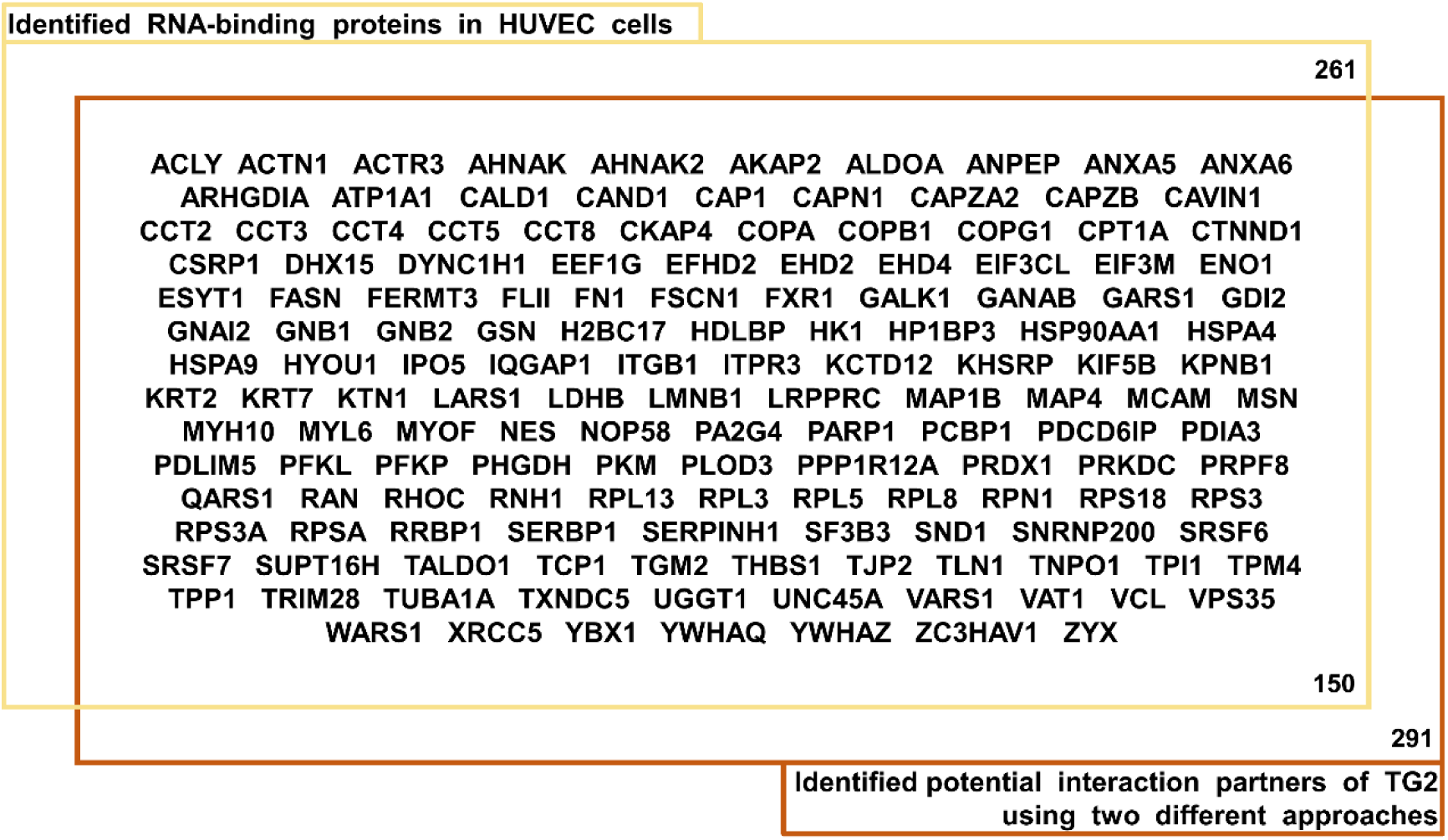
A significant overlap was found between the identified 3F-TG2-associated proteins and RNA-binding proteins isolated from HUVECs. The Venn diagram illustrates the proteins identified in this study as a 3F-TG2-associated protein and previously as RNA-binding proteins in HUVECs (Csaholczi et al., FEBSJ 2025).

The data obtained through this approach also support the involvement of TG2 in vesicular trafficking and transport of HUVEC, which is essential on both the apical and basal sides of the cell surface to maintain endothelial physiological functions. The most enriched GO Cellular Components are “Extracellular exosome”, “Vesicle”, “Extracellular space”, suggesting that TG2 is likely secreted by vesicular transport from HUVECs and might influence the composition of vesicles.

### Effect of the conformation change on the 3F-TG2-containing protein complexes

TG2 can physiologically exist in at least two distinct conformations, as determined by X-ray crystallography (Pinkas et al. 2007), suggesting that it may be capable of forming different interactions in its open or closed state. Modulator treatments were administered before or during the coimmunoprecipitation to investigate the effects of conformational changes on TG2’s interactors using anti-Flag M2 agarose resin (see the scheme in Figure 4a). The open conformation of TG2 was stabilised with TG2 inhibitor NC9 (Caron et al., 2012.; Akbar et al., 2017) pre-treatment of the cells, whereas the closed conformation was fixed by incubation with GTPγS during cell lysis and co-immunoprecipitation, as nucleotides are not cell-permeable.

In the presence of NC9 or GTPγS, 294 and 160 hits were detected after background subtraction, respectively (Supplementary Tables S10 and S11). This indicated a reduction in the interacting partners due to the modulators’ ability to stabilise distinct TG2 conformations. In the presence of GTPγS, the approximately 55% reduction in interactors (160 hits compared to 356 in the untreated case) is more pronounced than in the case of NC9 (294 hits compared to 356, representing approximately a 17% decrease), suggesting that stabilisation of the closed form can more effectively limit the number of TG2-associated proteins, likely due to the decreased surface area and less flexible structural dynamics.

Figure 6a and Supplementary Table S12 present the detailed grouping and number of interactor hits based on the conformations through which they can associate with TG2 across different experimental setups. The majority of the hits (138 out of the total 396 detected 3F-TG2-interactors, 34.8%) were found among TG2-associated proteins in all three experimental settings, suggesting that the majority of the interaction surfaces are independent of the open or closed conformations and are located on surface positions present in both the closed and open forms. This group includes several proteins involved in translation, either as ribonucleoproteins or translational factors, as well as components of the cytoskeleton and focal adhesion complex, along with those related to metabolism and signalling processes. The second largest group (122 hits) of interactors can bind to either untreated or NC9-treated TG2, likely because TG2 may be in an open conformation with a larger surface area. In this group, we identified membrane transport related terms such as “Transporter activity”, “Ion transmembrane transporter activity”, “ATPase-coupled cation transmembrane transporter activity”, “Transmembrane transporter activity”, “ATPase-coupled transmembrane transporter activity”, “Primary active transmembrane transporter activity”, “Inorganic molecular entity transmembrane transporter activity”, “Cation transmembrane transporter activity” as the most enriched GO Molecular Functions (Supplementary Table S13; example proteins: ATP13A1, ATP1A1/2A2, ATP5F1C, ATP6V0A1/2, ATP6V0D1, ATP6V1B2, ANO10, LRRC8A/C, MGT1, SFXN1/3, SLC12A4, SLC25A11, SLC2A1, TRPV2, VDAC3) and “Cadherin binding”, “RNA-binding” were less enriched. This raises the possibility that open or transglutaminase active TG2 may directly affect membrane transport processes, affecting transporter molecules.

**Figure 6.**
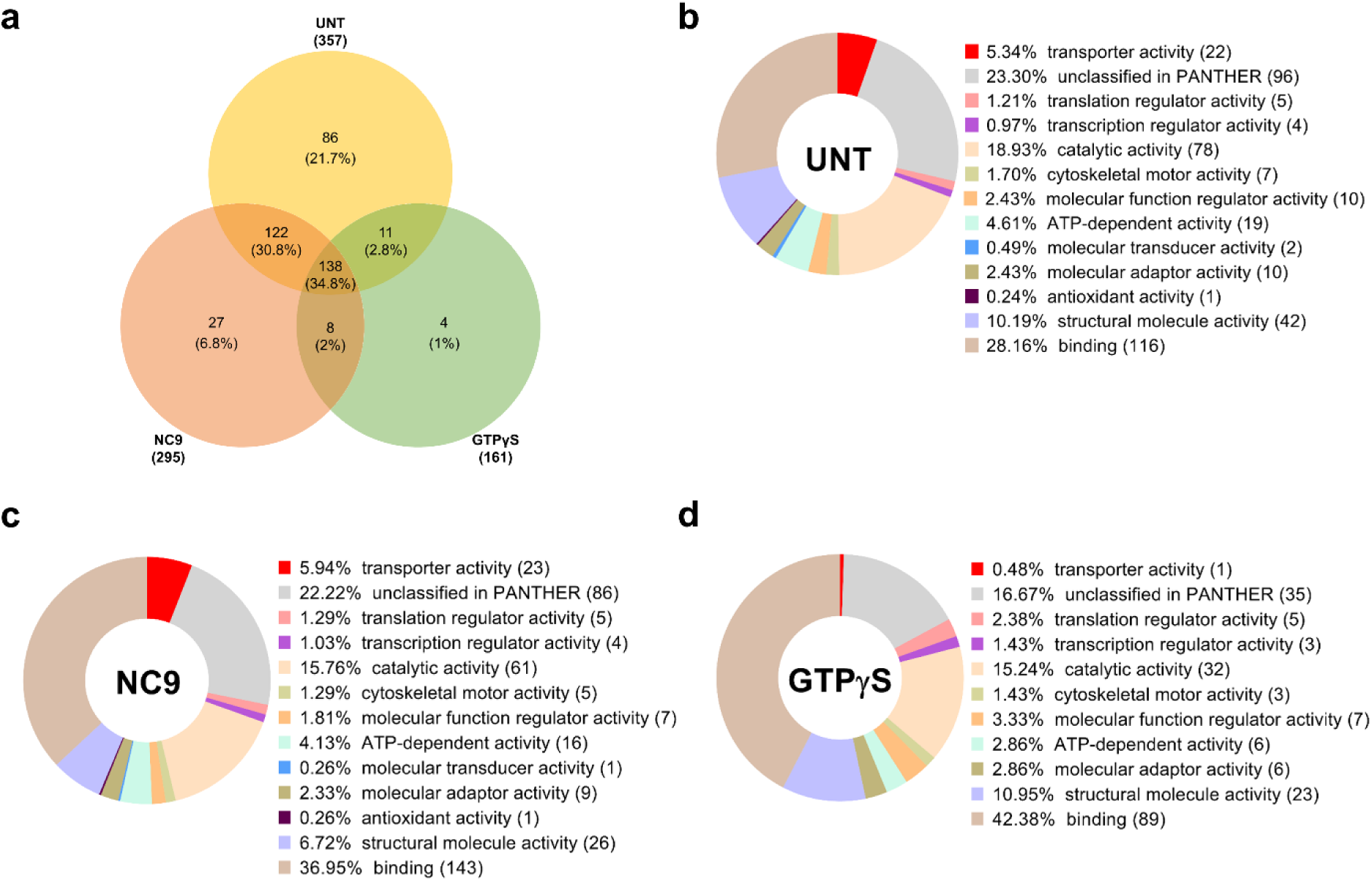
Conformation-dependent 3F-TG2-associated proteins in HUVECs and their GO-Slim Molecular functions. (a) Number and groups of 3F-TG2-associated proteins detected in a conformation-dependent manner (Venny 2.1). (b-d) The ratio of the most enriched GO-Slim Molecular functions of untreated (b), NC9-treated open (c), and GTPγS coincubated closed (d) TG2-associated proteins was assessed using the PANTHER Classification System. The number of proteins is indicated in brackets after the category (GraphPad 8.1 was used for visualisation).

To verify the observed differences in conformation-dependent TG2 interactome, we conducted data analysis using the PANTHER Classification System (Fig. 6b-d). We noted a change in the ratio of GO-Slim Molecular functions following GTPγS treatment. The stabilisation of the closed conformation reduced the ratio of interactors linked to the Transporter activity category (4 hits, 1.36%; ATP5F1A, CLIC1, PLS3, VAT1) compared to the untreated and NC9-treated samples (24 hits, 4.26% [APOA1, SFXN3, TRPV2, OSBPL3, SLC12A4, ATP5F1A, ATP1A1, ABCD3, CLIC1, SLC2A1, ATP6V1B2, ABCD1, ATP13A1, ATP5F1C, ITPR3, VDAC3, ANO10, VAT1, ATP2A2, SFXN1, SLC25A11, FLOT1, SLC25A24, ATP6V1E1] and 26 hits, 5.35% [e. g. ATP6V1B2, VDAC1, VDAC2, PLS3, SLC35B2], respectively), confirming our previous observation.

The large number of partners interacting only with untreated TG2 (93 hits) suggests that the basic protein’s flexibility and the potential for dynamic structural movements play a role in forming interactions. In this group, the most enriched molecular functions are related to RNA-binding, structural, cell-cell and cell-extracellular matrix molecule binding proteins. Interestingly, new interactions can also occur exclusively when a conformation-stabilising modulator is present (see a few interactors only in the case of the presence of NC9 and GTPγS).

## Discussion

TG2 has several kinds of catalytic capability, is localised in almost all cell compartments and has been implicated in diverse physiological functions and pathologic processes (Eckert et al., 2014). The question arises how it is possible to show such multifunctionality within a single polypeptide chain (Tangaraju et al., 2017)? One possibility is to develop a complex substrate system alongside diverse post-translational modifications based on the exact reaction mechanism (Csosz et al., 2008). Another possibility is binding various RNAs to participate in post-transcriptional regulations since TG2 has recently been recognised as an RNA-binding protein (Csaholczi et al., 2025). Alternatively or in parallel to the above mechanisms, a broad interaction network is developed that directly or indirectly regulates the functions and localisations of other proteins. In our study, we identified TG2-associated proteins in endothelial cells under physiological conditions and evaluated the potential biological functions of TG2.

To identify physiologically relevant TG2-associated proteins, two approaches were used: adding recombinant, site-specifically biotinylated TG2 to the cell extract, and as a novel method involving endogenous TG2 silencing combined with the expression of a triple Flag-tagged transgene TG2 as bait. The second approach increased the number of identified interactomes and reduced nonspecific binding of proteins. The triple Flag-tag extension enabled the possible detection of larger protein complexes, reducing steric hindrance during antibody binding. A significant advantage is the expression of tagged TG2 at levels comparable to normal conditions and the assembly of TG2-associated protein complexes within a living cellular environment. In living cells, the formation of functional protein complexes is a sequential, chaperone-assisted, and regulated process. Assembly chaperones, transient co-translational interactions, and subsequent conformational changes facilitate complex formation and the correct incorporation of subunits in sequence. Interaction with assembly chaperones can induce conformational changes necessary for binding to other proteins. GTP hydrolysis can also trigger conformational shifts that enhance affinity for specific proteins, although this process is less well understood *in situ* (Brown et al., 2025). Interaction with conformation-stabilising interactors, transglutaminase or GTPase substrates, which may influence the conformation-dependent association of proteins with TG2, could make the interactions particularly intriguing.

Previously, approximately 80 TG2-interacting proteins were listed and organised in the TRANSDAB database (Csosz et al., 2009). Most of these interactions were confirmed through co-immunoprecipitation and co-localisation experiments and reviewed in detail by Kanchan and colleagues (Kanchan et al., 2015). Another group was discovered via the co-immunoprecipitation study by Kim and colleagues (Kim et al., 2013). Here, we demonstrated significant overlap with these well-described TG2 interactors, further supporting the already known functions of TG2 in cell adhesion, migration, and cell-cell interactions. However, some well-known protein interacting partners were subtracted from our results due to their binding to the affinity resins during enrichment of TG2-associated proteins (e.g., FN1, CALM1, GSTP1, PRDX1). Over the past decade, some proteomic studies have significantly contributed to the understanding of the TG2 interactome and its biological significance, providing mechanistic insights for a more comprehensive functional understanding. Although these studies mostly used mouse models (Furini et al., 2018; Wilhelmus et al., 2022), they have greatly advanced our knowledge of TG2-associated and interacting proteins despite relying on polyclonal antibodies to bind TG2-associated protein complexes. In the mouse UUO model, exosomal proteins (e.g., ALIX, FLOT2, PRDX2, RAB1A, SDC4) have been linked to TG2, facilitating its externalisation (Furini et al., 2018). The APP23 Alzheimer’s disease mouse model, showed an increase in TG2 interactors and proteins involved in synapse assembly, synaptic transmission, and cell adhesion were enriched among these (Wilhelmus et al., 2022). In Prof. Piacentini’s laboratory, tandem affinity purification using HA and Flag-tagged overexpressed TG2 under various stress conditions was employed to uncover the TG2 interactome and potential key interactors involved in proteostasis in both mouse and human cells (Altuntas et al., 2015; D’Eletto et al., 2019.). Based on these interactome analyses, TG2-interacting chaperones (HSP70, HSP90, HSP20, HSPHSP27, HSPA6, BAG2/3, DNAJ A1/B1/C7) and autophagic receptors (p62, NBR1) were identified, indicating TG2’s role in stress response and the removal of unnecessary proteins. In cases of deficient proteasomal activity, TG2 in complex with BAG3 can collect and direct ubiquitinated proteins to the multivesicular bodies for either lysosomal degradation or exosomal externalisation, interacting with TSG101 and Alix (D’Eletto et al., 2019.). Most of these stress-related interactors were not observed in our non-stressed HUVEC system.

Extracellular exosomes are the most enriched GO Cellular Component among the identified TG2-associated proteins, suggesting that under physiological conditions, TG2 plays a role in the exosomal communication of endothelial cells. It could facilitate the regulation of extracellular vesicle (EV) formation or stabilise their structures by catalysing the covalent cross-linking of proteins within these vesicles (Shinde et al., 2020). The formation of these exosomes in endothelial cells can facilitate communication with other cells in both basal and apical directions. Exosomes are particularly released towards the apical side into the circulation, but TG2 is present in both types of secretomes from HUVEC and HCAEC cells (Wei et al., 2019). Additionally, TG2-associated proteins show significant overlap with the secreted proteins from HUVECS released via vesicular transport (ACTN1, AHNAK, CALR, CAVIN1, ENO1, FSCN1, SERPIN1, TALDO1, TCP1, TLN1, TPO1, VAT1, RACK1, VAPA, etc.) (Wei et al., 2019). This occurs on both the basal and apical sides of the endothelial cells, which communicate intensively with the extracellular matrix and potentially with distant organs. The content of secreted exosomes influences recipient cells; in healthy endothelial cells, the proteins within these exosomes may protect against the development of cardiovascular diseases (Davidson et al., 2018).

TG2 is a unique protein capable of adopting diverse conformational states. According to our results, most of the associated proteins did not exhibit significant TG2 conformation dependence, but stabilising the open form increased the number of transporters within the complexes. This open conformation could provide a surface for interaction with various membrane transporters. It has been published that TG2 can accept the Ca^2+^-selective TRPV5 as a substrate, inhibiting it through covalent modification (Boros et al., 2012). This could explain the altered mitochondrial function observed in TG2-KO mice, and may also contribute to the regulation of cellular energy production, thereby linking TG2 to human pathological conditions (Rossin et al., 2015).

The presence of numerous RNA-binding proteins in the TG2 interactome, along with their substantial overlap with the recently identified proteins which bind RNA in HUVEC (Csaholczi et al., 2025), was the most surprising finding of our study. Some of these proteins are specialised for RNA or nucleic acid binding (e. g. NOP56, RPs, RALY, SART1, SRSF1, SUPT16H), affecting their maturation, stabilisation, localisation, and function. Many of the revealed TG2-associated proteins are known for their biological roles (e. g. PKM, ENO1, ALDOA, MAP4, ATP5F1C, PDIA3, CALR, DYNC1H1), and it has recently been recognised that they can also bind RNA, showing a moonlighting property. They can influence RNA function, but conversely, the protein can also be modulated by RNA binding. TG2 might belong to these moonlighting proteins because we recently identified its RNA-binding ability (Csaholczi et al., 2025). With this capability, and considering the large number of its interacting partners, TG2 could act as an important hub in the RNA-protein regulatory network, which warrants further investigations in the future.

## Author Contributions

Conceptualisation, R.K.; methodology, B.C., I.R.K-S. and R.K.; software, B.C., Z. C-S.; formal analysis, visualisation, B.C., K. J., Z. C-S. and R.K.; investigation, B.C., Z. C-S., K.J. and R.K.; data curation, B.C., Z. C-S.; writing—original draft preparation, R.K. and B.C.; writing— review and editing, R.K., I.R.K-S., K.J. and L.F.; funding acquisition, resources, R.K. and L.F. All authors have read and agreed to the published version of the manuscript.

## Acknowledgements

We thank Dr. János Mótyán for the critical reading and suggestions for the manuscript preparation and Jennifer Nagy for technical assistance in the laboratory (Department of Biochemistry and Molecular Biology, University of Debrecen). The authors are thankful to Boglárka Kerekesné Tóth for her contribution to the generation of the N-BAP-biotin rhTG2 construct. We appreciate the helpful comments and service of Gergő Kalló and the Proteomics Core Facility (mass spectrometry analysis) in the Department of Biochemistry and Molecular Biology, University of Debrecen. The NC9 inhibitor was a kind gift from Professor Jeffrey W Keillor (Department of Chemistry and Biomolecular Sciences, University of Ottawa, Ottawa, ON, Canada). This work was supported by the following grant: National Research, Development and Innovation Office [NKFIH-K129139] and by the University of Debrecen Program for Scientific Publication. B. C. was supported by the EKÖP-24-3-II-DE-213 University Research Scholarship Program of the Ministry for Culture and Innovation from the source of the National Research, Development and Innovation Fund.

## Data Availability Statement

Data are available from the corresponding author upon reasonable request.

**Abbreviations**
BAP: biotin acceptor peptide;
CoIP: coimmunoprecipitation;
EV: extracellular vesicle;
HUVEC: human umbilical cord vein endothelial cell;
IPTG: Isopropyl β-d-1-thiogalactopyranoside;
KD: knock-down;
LC-MS/MS: liquid chromatography-tandem mass spectrometry;
RBP: RNA-binding protein;
RNA: ribonucleic acid;
TG2: transglutaminase 2;
TGF-β1: transforming growth factor beta-1;
UUO: unilateral ureteric obstruction;
UTR: untranslated region
VEGF: vascular endothelial growth factor

