## Supplementary tables for "The transglutaminase 2 interactome in HUVECs suggests its participation in an RNA-binding protein network"

**S1: Identified nonspecifically bound proteins to High Capacity Neutravidin agarose resin from HUVEC extract (controls, subtracted from sample hits, n=3)**

| Gene Name | Name of proteins |
|---|---|
| ACTB | Actin, cytoplasmic 1 |
| ACTG1 | Actin, cytoplasmic 2 |
| ACTN1 | Alpha-actinin-1 |
| ACTN4 | Alpha-actinin-4 |
| AHNAK | Neuroblast differentiation-associated protein AHNAK |
| ALB | Albumin |
| ALDOA | Fructose-bisphosphate aldolase A |
| ANPEP | Aminopeptidase N |
| ANXA1 | Annexin A1 |
| ANXA2 | Annexin A2 |
| ANXA5 | Annexin A5 |
| ANXA6 | Annexin A6 |
| ATP1A1 | Sodium/potassium-transporting ATPase subunit alpha-1 |
| ATP2A2 | Sarcoplasmic/endoplasmic reticulum calcium ATPase 2 |
| ATP5F1A | ATP synthase subunit alpha, mitochondrial |
| ATP5F1B | ATP synthase subunit beta, mitochondrial |
| ATP5PO | ATP synthase subunit O, mitochondrial |
| CAD | CAD protein |
| CALM1 | Calmodulin-1 |
| CALR | Calreticulin |
| CANX | Isoform 2 of Calnexin |
| CAP1 | Adenylyl cyclase-associated protein 1 |
| CAVIN1 | Caveolae-associated protein 1 |
| CFL1 | Cofilin-1 |
| CKAP4 | Cytoskeleton-associated protein 4 |
| CLIC1 | Chloride intracellular channel protein 1 |
| CLTC | Clathrin heavy chain 1 |
| CNN2 | Calponin-2 |
| COLGALT1 | Procollagen galactosyltransferase 1 |
| COPG1 | Coatomer subunit gamma-1 |
| CORO1B | Coronin-1B |
| CORO1C | Coronin-1C |
| CPS1 | Carbamoyl-phosphate synthase [ammonia], mitochondrial |
| CPT1A | Carnitine O-palmitoyltransferase 1, liver isoform |
| CRIP2 | Cysteine-rich protein 2 |
| CTNNA1 | Catenin alpha-1 |
| CSE1L | Exportin-2 |
| DBN1 | Drebrin |
| DDX17 | Probable ATP-dependent RNA helicase DDX17 |
| DDX3X | ATP-dependent RNA helicase DDX3X |
| DHX15 | ATP-dependent RNA helicase DHX15 |
| DHX9 | ATP-dependent RNA helicase A |
| EEF1A1 | Elongation factor 1-alpha 1 |
| EEF1D | Elongation factor 1-delta |
| EEF1E1 | Eukaryotic translation elongation factor 1 epsilon-1 |
| EEF1G | Elongation factor 1-gamma |
| EEF2 | Elongation factor 2 |
| EFHD2 | EF-hand domain-containing protein D2 |
| EHD1 | EH domain-containing protein 1 |
| EHD2 | Isoform 2 of EH domain-containing protein 2 |
| EHD4 | EH domain-containing protein 4 |
| EIF3M | Eukaryotic translation initiation factor 3 subunit M |
| EIF4A1 | Eukaryotic initiation factor 4A-I |
| ELAVL1 | Isoform 2 of ELAV-like protein 1 |
| ENG | Endoglin |
| ENO1 | Alpha-enolase |
| EWSR1 | RNA-binding protein EWS |
| FARSA | Phenylalanine--tRNA ligase alpha subunit |
| FERMT3 | Fermitin family homolog 3 |
| FLNA | Filamin-A |
| FLNB | Filamin-B |
| FLNC | Isoform 2 of Filamin-C |
| GAPDH | Glyceraldehyde-3-phosphate dehydrogenase |
| GBE1 | 1,4-alpha-glucan-branching enzyme |
| GLIPR2 | Golgi-associated plant pathogenesis-related protein 1 |
| GLRX3 | Glutaredoxin-3 |
| GSTP1 | Glutathione S-transferase P |
| H1-2 | Histone H1.2 |
| H1-3 | Histone H1.3 |
| H1-4 | Histone H1.4 |

| | |
|---|---|
| H1-5 | Histone H1.5 |
| H2AC11 | Histone H2A type 1 |
| H2AC4 | Histone H2A type 1-B/E |
| H2AC6 | Histone H2A type 1-C |
| H2AJ | Histone H2A.J |
| H2BC12 | Histone H2B type 1-K |
| H2BC18 | Histone H2B type 2-F |
| H2BC4 | Histone H2B type 1-C/E/F/G/I |
| H2BC9 | Histone H2B type 1-H |
| H3C1 | Histone H3.1 |
| H4C1 | Histone H4 |
| HIGD1A | HIG1 domain family member 1A, mitochondrial |
| HMGA1 | High mobility group protein HMG-I/HMG-Y |
| HMGA2 | High mobility group protein HMGI-C |
| HNRNPA1 | Heterogeneous nuclear ribonucleoprotein A1 |
| HNRNPA2B1 | Heterogeneous nuclear ribonucleoproteins A2/B1 |
| HNRNPA3 | Heterogeneous nuclear ribonucleoprotein A3 |
| HNRNPC | Heterogeneous nuclear ribonucleoproteins C1/C2 |
| HNRNPD | Heterogeneous nuclear ribonucleoprotein D0 |
| HNRNPDL | Heterogeneous nuclear ribonucleoprotein D-like |
| HNRNPF | Heterogeneous nuclear ribonucleoprotein F |
| HNRNPH1 | Heterogeneous nuclear ribonucleoprotein H |
| HNRNPK | Heterogeneous nuclear ribonucleoprotein K |
| HNRNPL | Heterogeneous nuclear ribonucleoprotein L |
| HNRNPM | Heterogeneous nuclear ribonucleoprotein M |
| HNRNPR | Heterogeneous nuclear ribonucleoprotein R |
| HNRNPU | Isoform 2 of Heterogeneous nuclear ribonucleoprotein U |
| HNRNPUL2 | Heterogeneous nuclear ribonucleoprotein U-like protein 2 |
| HP1BP3 | Heterochromatin protein 1-binding protein 3 |
| HRNR | Hornerin |
| HSP90AA1 | Heat shock protein HSP 90-alpha |
| HSP90AB1 | Heat shock protein HSP 90-beta |
| HSP90B1 | Endoplasmin |
| HSPA1L | Heat shock 70 kDa protein 1-like |
| HSPA5 | Endoplasmic reticulum chaperone BiP |
| HSPA8 | Heat shock cognate 71 kDa protein |
| HSPB1 | Heat shock protein beta-1 |
| HSPD1 | 60 kDa heat shock protein, mitochondrial |
| IGF2BP3 | Isoform 2 of Insulin-like growth factor 2 mRNA-binding protein 3 |
| IGKV2D-24 | Probable non-functional immunoglobulin kappa variable 2D-24 |
| IGKV2D-26 | Immunoglobulin kappa variable 2D-26 |
| ILF3 | Isoform 7 of Interleukin enhancer-binding factor 3 |
| IQGAP1 | Ras GTPase-activating-like protein IQGAP1 |
| ITGB1 | Integrin beta-1 |
| KPNB1 | Importin subunit beta-1 |
| KRT1 | Keratin, type II cytoskeletal 1 |
| KRT10 | Keratin, type II cytoskeletal 10 |
| KRT14 | Keratin, type II cytoskeletal 14 |
| KRT18 | Keratin, type II cytoskeletal 18 |
| KRT2 | Keratin, type II cytoskeletal 2 |
| KRT5 | Keratin, type II cytoskeletal 5 |
| KRT6A | Keratin, type II cytoskeletal 6a |
| KRT6B | Keratin, type II cytoskeletal 6b |
| KRT7 | Keratin, type II cytoskeletal 7 |
| KRT77 | Keratin, type II cytoskeletal 77 |
| KRT8 | Keratin, type II cytoskeletal 8 |
| KRT9 | Keratin, type II cytoskeletal 9 |
| KTN1 | Kinectin |
| LDHA | L-lactate dehydrogenase A chain |
| LETM1 | Leucine Zipper And EF-Hand Containing Transmembrane Protein 1 |
| LGALS1 | Galectin-1 |
| LIMA1 | LIM domain and actin-binding protein 1 |
| LMNA | Prelamin-A/C |
| LMNB1 | Lamin-B1 |
| LMNB2 | Lamin-B2 |
| LMO7 | Isoform 3 of LIM domain only protein 7 |
| LMO7B | LIM Domain 7 |
| MACROH2A1 | Core histone macro-H2A.1 |
| MAPK3 | Mitogen-activated protein kinase 3 |
| MATR3 | Matrin-3 |
| MSN | Moesin |
| MTDH | Protein LYRIC |
| MTHFD1 | C-1-tetrahydrofolate synthase, cytoplasmic |
| MVP | Major vault protein |

| | |
|---|---|
| MYH10 | Myosin-10 |
| MYH14 | Myosin-14 |
| MYH9 | Myosin-9 |
| MYL12B | Myosin regulatory light chain 12B |
| MYL6 | Myosin light polypeptide 6 |
| MYO1C | Unconventional myosin-Ic |
| MYO1D | Unconventional myosin-Id |
| MYO1E | Unconventional myosin-Ie |
| MYO5A | Isoform 2 of Unconventional myosin-Va |
| MYO6 | Isoform 6 of Unconventional myosin-VI |
| MYOF | Myoferlin |
| NAP1L1 | Nucleosome assembly protein 1-like 1 |
| NCL | Nucleolin |
| NES | Nestin |
| NONO | Non-POU domain-containing octamer-binding protein |
| NPM1 | Nucleophosmin |
| PA2G4 | Proliferation-associated protein 2G4 |
| PABPC1 | Polyadenylate-binding protein 1 |
| PABPN1 | Polyadenylate-binding protein 2 |
| PCBP1 | Poly(rC)-binding protein 1 |
| PCMT1 | Isoform 2 of Protein-L-isoaspartate(D-aspartate) O-methyltransferase |
| PDIA3 | Protein disulfide-isomerase A3 |
| PDIA6 | Isoform 2 of Protein disulfide-isomerase A6 |
| PDLIM7 | PDZ and LIM domain protein 7 |
| PFKP | ATP-dependent 6-phosphofructokinase, platelet type |
| PFN1 | Profilin-1 |
| PGK1 | Phosphoglycerate kinase 1 |
| PHB1 | Prohibitin 1 OS=Homo sapiens OX=9606 GN=PHB1 PE=1 SV= |
| PKM | Pyruvate kinase PKM |
| PLEC | Plectin |
| PPIA | Peptidyl-prolyl cis-trans isomerase A |
| PPP1R12A | Protein phosphatase 1 regulatory subunit 12A |
| PTBP1 | Polypyrimidine tract-binding protein 1 |
| RASIP1 | Ras-interacting protein 1 |
| RBBP4 | Histone-binding protein RBBP4 |
| RBM14 | Isoform 5 of RNA-binding protein 14 |
| RBMX | RNA-binding motif protein, X chromosome |
| RPL10A | 60S ribosomal protein L10a |
| RPL11 | 60S ribosomal protein L11 |
| RPL12 | 60S ribosomal protein L12 |
| RPL13 | 60S ribosomal protein L13 |
| RPL13A | 60S ribosomal protein L13a |
| RPL14 | 60S ribosomal protein L14 |
| RPL15 | 60S ribosomal protein L15 |
| RPL17 | 60S ribosomal protein L17 |
| RPL18 | 60S ribosomal protein L18 |
| RPL18A | 60S ribosomal protein L18a |
| RPL19 | 60S ribosomal protein L19 |
| RPL23 | 60S ribosomal protein L23 |
| RPL23A | 60S ribosomal protein L23a |
| RPL24 | 60S ribosomal protein L24 |
| RPL3 | 60S ribosomal protein L3 |
| RPL4 | 60S ribosomal protein L4 |
| RPL6 | 60S ribosomal protein L6 |
| RPL7 | 60S ribosomal protein L7 |
| RPL7A | 60S ribosomal protein L7a |
| RPL8 | 60S ribosomal protein L8 |
| RPLP0 | 60S acidic ribosomal protein P0 |
| RPN1 | Dolichyl-diphosphooligosaccharide--protein glycosyltransferase subunit 1 |
| RPN2 | Dolichyl-diphosphooligosaccharide--protein glycosyltransferase subunit 2 |
| RPS11 | 40S ribosomal protein S11 |
| RPS12 | 40S ribosomal protein S12 |
| RPS16 | 40S ribosomal protein S16 |
| RPS17 | 40S ribosomal protein S17 |
| RPS19 | 40S ribosomal protein S19 |
| RPS2 | 40S ribosomal protein S2 |
| RPS25 | 40S ribosomal protein S25 |
| RPS27A | Ubiquitin-40S ribosomal protein S27a |
| RPS3 | 40S ribosomal protein S3 |
| RPS3A | 40S ribosomal protein S3a |
| RPS4X | 40S ribosomal protein S4, X isoform |
| RPS5 | 40S ribosomal protein S5 |
| RPS6 | 40S ribosomal protein S6 |
| RPS7 | 40S ribosomal protein S7 |

| | |
|---|---|
| RPS8 | 40S ribosomal protein S8 |
| RPS9 | 40S ribosomal protein S9 |
| RPSA | 40S ribosomal protein SA |
| RRBP1 | Ribosome-binding protein 1 |
| RSL1D1 | Ribosomal L1 domain-containing protein 1 |
| S100A11 | Protein S100-A11 |
| SEC22B | Vesicle-trafficking protein SEC22b |
| SEC23A | Protein transport protein Sec23A |
| SERPINH1 | Serpin H1 |
| SFPQ | Splicing factor, proline- and glutamine-rich |
| SLC25A11 | Mitochondrial 2-oxoglutarate/malate carrier protein |
| SLC25A3 | Phosphate carrier protein, mitochondrial |
| SLC25A5 | ADP/ATP translocase 2 |
| SLC25A6 | ADP/ATP translocase 3 |
| SND1 | Staphylococcal nuclease domain-containing protein 1 |
| SNX2 | Sorting nexin-2 |
| SPCS2 | Signal peptidase complex subunit 2 |
| SPECC1L | Cytospin-A |
| SPTAN1 | Spectrin alpha chain, non-erythrocytic 1 |
| SPTBN1 | Spectrin beta chain, non-erythrocytic 1 |
| SRP14 | Signal recognition particle 14 kDa protein |
| SUPT16H | FACT complex subunit SPT16 |
| SVIL | Supervillin |
| SYNCRIP | Isoform 3 of Heterogeneous nuclear ribonucleoprotein Q |
| SYNE2 | Nesprin-2 OS=Homo sapiens |
| SYNPO | Synaptopodin |
| TAGLN2 | Transgelin-2 |
| TALDO1 | Transaldolase |
| TARDBP | TAR DNA-binding protein 43 |
| TGM2 | Protein-glutamine gamma-glutamyltransferase 2 |
| THBS1 | Thrombospondin-1 |
| THRAP3 | Thyroid hormone receptor-associated protein 3 |
| TLN1 | Talin-1 |
| TPM3 | Isoform 5 of Tropomyosin alpha-3 chain |
| TPP1 | Tripeptidyl-peptidase 1 |
| TUBA1A | Tubulin alpha-1A chain |
| TUBA1B | Tubulin alpha-1B chain |
| TUBB | Tubulin beta chain |
| TUBB4B | Tubulin beta-4B chain |
| TUBB6 | Tubulin beta-6 chain |
| TXNDC5 | Thioredoxin domain-containing protein 5 |
| UBA1 | Ubiquitin-like modifier-activating enzyme 1 |
| UBB | Polyubiquitin-B |
| UBC | Polyubiquitin-C |
| VAT1 | Synaptic vesicle membrane protein VAT-1 homolog |
| VCP | Transitional endoplasmic reticulum ATPase |
| VDAC2 | Isoform 1 of Voltage-dependent anion-selective channel protein 2 |
| VIM | Vimentin |
| XRCC6 | X-ray repair cross-complementing protein 6 |
| YBX1 | Y-box-binding protein 1 |
| ZNF185 | Zinc finger protein 185 |

**S2: Identified N-BAP-biotin rhTG2-associated proteins in HUVEC using High Capacity Neutravidin agarose resin for the affinity chromatography (n=3)**

| Gene Name | Name of TG2-associated proteins | Identified unique peptide count | Is it a known TG2 interacting partner? |
|---|---|---|---|
| PLEC1 | Plectin | 215 | |
| DYNC1H1 | Cytoplasmic dynein 1 heavy chain 1 | 24 | |
| GNAI2 | Guanine nucleotide-binding protein G(i) subunit alpha-2 | 22 | |
| GNB1 | Guanine nucleotide-binding protein G(I)/G(S)/G(T) subunit beta-1 | 11 | |
| NT5E | 5'-nucleotidase | 9 | |
| PARP1 | Poly [ADP-ribose] polymerase 1 | 9 | |
| SSRP1 | FACT complex subunit SSRP1 | 9 | |
| RPS18 | 40S ribosomal protein S18 | 8 | |
| XRCC5 | X-ray repair cross-complementing protein 5 | 8 | |
| CD59 | CD59 glycoprotein | 7 | |
| FMNL2 | Formin-like protein 2 | 7 | |
| FN1 | Fibronectin | 7 | yes |
| PRKDC | DNA-dependent protein kinase catalytic subunit | 7 | |
| COPB1 | Coatomer subunit beta | 6 | yes |
| DDOST | Dolichyl-diphosphooligosaccharide--protein glycosyltransferase 48 kDa subunit | 6 | |
| FLII | Protein flightless-1 homolog | 6 | |
| GNB2 | Guanine nucleotide-binding protein G(I)/G(S)/G(T) subunit beta-2 | 6 | |
| PDI | Protein disulfide-isomerase | 6 | |
| RALA | Ras-related protein Ral-A | 6 | |
| TMPO | Lamina-associated polypeptide 2, isoforms beta/gamma | 6 | |
| AP2A1 | AP-2 complex subunit alpha-1 | 5 | |
| CAND1 | Isoform 2 of Cullin-associated NEDD8-dissociated protein 1 | 5 | |
| CDC42BPB | Serine/threonine-protein kinase MRCK beta | 5 | |
| GANAB | Neutral alpha-glucosidase AB | 5 | |
| MCAM | Melanoma Cell Adhesion Molecule | 5 | |
| YES1 | Tyrosine-protein kinase Yes | 5 | |
| ATAD3A | Isoform 2 of ATPase family AAA domain-containing protein 3A | 4 | |
| EIF3A | Eukaryotic translation initiation factor 3 subunit A | 4 | |
| EIF3CL | Eukaryotic translation initiation factor 3 subunit C-like protein | 4 | |
| EMD | Emerin | 4 | |
| H2BC17 | Histone H2B type 1-O | 4 | |
| IFI16 | Gamma-interferon-inducible protein 16 | 4 | |
| MBOAT7 | Lysophospholipid acyltransferase 7 | 4 | |
| PHB2 | Prohibitin-2 | 4 | |
| QARS1 | Glutamine--tRNA ligase | 4 | |
| RACK1 | Receptor of activated protein C kinase 1 | 4 | |
| TTN | Titin | 4 | |
| UQCRC1 | Cytochrome B-C1 Complex Subunit 1, Mitochondrial | 4 | |
| YWHAZ | 14-3-3 protein zeta/delta | 4 | |
| ACAT1 | Acetyl-CoA acetyltransferase, mitochondrial | 3 | |
| ACLY | ATP-citrate synthase | 3 | |
| ANXA | Annexin A | 3 | |
| ARHGDIA | Rho GDP-dissociation inhibitor 1 | 3 | |
| CAPZA1 | F-actin-capping protein subunit alpha-1 | 3 | |
| COPA | Coatomer subunit alpha | 3 | yes |
| CTNND1 | Catenin delta-1 | 3 | |
| CTSB | Cathepsin B | 3 | |
| GNG12 | Guanine nucleotide-binding protein G(I)/G(S)/G(O) subunit gamma-12 | 3 | |
| GSN | Gelsolin | 3 | |
| HDLBP | Vigilin | 3 | |
| HYOU1 | Hypoxia up-regulated protein 1 | 3 | |
| PCBP2 | Poly(rC)-binding protein 2 | 3 | |
| PDCD6IP | Programmed cell death 6-interacting protein | 3 | |
| PFKL | ATP-dependent 6-phosphofructokinase, liver type | 3 | |
| RPS26 | 40S ribosomal protein S26 | 3 | |
| UQCRC2 | Cytochrome b-c1 complex subunit 2, mitochondrial | 3 | |
| VDAC1 | Voltage-dependent anion-selective channel protein 1 | 3 | |
| ATP6V1B2 | V-type proton ATPase subunit B, brain isoform | 2 | |
| CAPZB | F-actin-capping protein subunit beta | 2 | |
| CAV1 | Caveolin-1 | 2 | |
| DLD | Dihydrolipoyl dehydrogenase, mitochondrial | 2 | |
| ERLIN1 | Erlin-1 | 2 | |
| ESYT1 | Extended synaptotagmin-1 | 2 | |
| ESYT2 | Extended Synaptotagmin 2 | 2 | |
| FASN | Fatty acid synthase | 2 | |
| H1-0 | Histone H1.0 | 2 | |
| HK1 | Hexokinase-1 | 2 | |

| | | | |
|---|---|---|---|
| IGF2R | Cation-independent mannose-6-phosphate receptor | 2 | |
| IPO5 | Importin-5 | 2 | |
| KIF5B | Kinesin-1 heavy chain | 2 | |
| LARS1 | Leucine--tRNA ligase, cytoplasmic | 2 | |
| LRPPRC | Leucine-rich PPR motif-containing protein, mitochondrial | 2 | |
| MCM7 | DNA Replication Licensing Factor MCM7 | 2 | |
| MMP1 | Interstitial collagenase | 2 | yes |
| MYADM | Myeloid-associated differentiation marker | 2 | |
| MYO1B | Unconventional myosin-Ib | 2 | |
| NOP56 | Nucleolar protein 56 | 2 | |
| NOP58 | Nucleolar protein 58 | 2 | |
| PDLIM5 | PDZ and LIM domain protein 5 | 2 | |
| RAB10 | Ras-related protein Rab-10 | 2 | |
| RAN | GTP-binding nuclear protein Ran | 2 | |
| RPL5 | 60S ribosomal protein L5 | 2 | |
| SFXN1 | Sideroflexin-1 | 2 | |
| SPRED2 | Sprouty-related, EVH1 domain-containing protein 2 | 2 | |
| TNPO1 | Transportin-1 | 2 | |
| TRPV2 | Transient receptor potential cation channel subfamily V member 2 | 2 | |
| TUBB3 | Tubulin beta-3 chain | 2 | yes |
| TUBB4A | Tubulin beta-4A chain | 2 | yes |
| UBA52 | Ubiquitin-60S ribosomal protein L40 | 2 | |
| UNCA45A | Protein unc-45 homolog A | 2 | |
| VPS35 | Vacuolar protein sorting-associated protein 35 | 2 | |
| ZYX | Zyxin | 2 | yes |
| ACIN1 | Apoptotic chromatin condensation inducer in the nucleus | 1 | |
| AHNAK2 | Protein AHNAK2 | 1 | |
| AKAP2 | A-kinase anchor protein 2 | 1 | |
| AP2B1 | AP-2 complex subunit beta | 1 | |
| ARHGAP32 | Rho GTPase-activating protein 32 | 1 | |
| ATP5F1C | ATP synthase subunit gamma, mitochondrial | 1 | |
| BRCA1 | BRCA1 protein | 1 | |
| CAPN1 | Calpain 1 | 1 | |
| CAPZA2 | F-actin-capping protein subunit alpha-2 | 1 | |
| CCDC175 | Coiled-coil domain-containing protein 175 | 1 | |
| CCT3 | T-complex protein 1 subunit gamma | 1 | |
| CDC42 | Cell division control protein 42 homolog | 1 | |
| CDCP1 | CUB Domain Containing Protein 1 | 1 | |
| CLEC19A | C-Type Lectin Domain Containing 19A | 1 | |
| CNTRL | Centriolin | 1 | |
| COL5A2 | N-acylneuraminate cytidylyltransferase | 1 | |
| CYB5R3 | NADH-cytochrome b5 reductase 3 | 1 | yes |
| DLST | Dihydrolipoyllysine-residue succinyltransferase component of 2-oxoglutarate dehydrogenase complex, mitochondrial | 1 | |
| DNAJA1 | DnaJ homolog subfamily A member 1 | 1 | yes |
| DST | Dystonin | 1 | |
| EFEMP1 | EGF-containing fibulin-like extracellular matrix protein 1 | 1 | |
| FBL | rRNA 2'-O-methyltransferase fibrillarin | 1 | |
| FHL2 | Four and a half LIM domains protein 2 | 1 | |
| FKBP1A | Peptidyl-prolyl cis-trans isomerase FKBP1A | 1 | |
| G3BP1 | Ras GTPase-activating protein-binding protein 1 | 1 | |
| GALK1 | Galactokinase | 1 | |
| GARS1 | Glycine--tRNA ligase | 1 | |
| GLG1 | Isoform 3 of Golgi apparatus protein 1 | 1 | |
| H2AC14 | Histone H2A type 1-J | 1 | |
| H2AZ1 | Histone H2A.Z | 1 | |
| HEXA | Hexosaminidase Subunit Alpha | 1 | |
| HNRNPAB | Isoform 2 of Heterogeneous nuclear ribonucleoprotein A/B | 1 | |
| HSPA4 | Heat shock 70 kDa protein 4 | 1 | |
| ITGA3 | Integrin alpha-3 | 1 | |
| ITGA5 | Integrin alpha-5 | 1 | |
| KHSRP | Far upstream element-binding protein 2 | 1 | |
| LMAN1 | Protein ERGIC-53 | 1 | |
| MAP1B | Microtubule-associated protein 1B | 1 | |
| MARS1 | Methionine--tRNA ligase, cytoplasmic | 1 | |
| MGST3 | Microsomal glutathione S-transferase 3 | 1 | |
| NUP93 | Nuclear pore complex protein Nup93 | 1 | |
| OCIAD2 | OCIA domain-containing protein 2 | 1 | |
| PAPSS2 | Bifunctional 3'-phosphoadenosine 5'-phosphosulfate synthase 2 | 1 | |
| PHGDH | D-3-phosphoglycerate dehydrogenase | 1 | |
| PLAG1 | PLAG1 Zinc Finger | 1 | |
| PLOD2 | Procollagen-lysine,2-oxoglutarate 5-dioxygenase 2 | 1 | |
| PPP1CC | Serine/threonine-protein phosphatase PP1-gamma catalytic subunit | 1 | |
| PRDX1 | Peroxiredoxin-1 | 1 | yes |

| | | | |
|---|---|---|---|
| PRDX6 | Peroxiredoxin-6 | 1 | |
| PSMD2 | 26S proteasome non-ATPase regulatory subunit 2 | 1 | |
| RAB11B | Ras-related protein Rab-11B | 1 | |
| RAB7A | Ras-related protein Rab-7a | 1 | |
| RAC2 | Ras-related C3 botulinum toxin substrate 2 | 1 | |
| RAP1A | Ras-related protein Rap-1A | 1 | |
| RHOC | Rho-related GTP-binding protein RhoC | 1 | |
| SEC62 | Translocation protein SEC62 | 1 | |
| SFXN3 | Sideroflexin-3 | 1 | |
| SLTM | SAFB-like transcription modulator | 1 | |
| SMC1A | Structural maintenance of chromosomes protein 1A | 1 | |
| SRSF1 | Serine/arginine-rich splicing factor 1 | 1 | |
| SRSF3 | Serine/arginine-rich splicing factor 3 | 1 | |
| SSR4 | Translocon-associated protein subunit delta | 1 | |
| STING1 | Stimulator of interferon genes protein | 1 | |
| STOM | Stomatin | 1 | |
| SUN2 | SUN Domain-Containing Protein 2 | 1 | |
| SURF4 | Surfeit locus protein 4 | 1 | |
| TCIRG1 | V-type proton ATPase 116 kDa subunit a 3 | 1 | |
| TGFB1I1 | Transforming growth factor beta-1-induced transcript 1 protein | 1 | |
| TIMM50 | Mitochondrial import inner membrane translocase subunit TIM50 | 1 | |
| TJP2 | Tight Junction Protein 2 (Zona occludens 2) | 1 | |
| TMED10 | Transmembrane emp24 domain-containing protein 10 | 1 | |
| TMED9 | Transmembrane emp24 domain-containing protein 9 | 1 | |
| TOP2B | DNA topoisomerase 2-beta | 1 | |
| TPM1 | Tropomyosin 1 | 1 | |
| UBAP2L | Ubiquitin-associated protein 2-like | 1 | |
| UNC45A | Protein unc-45 homolog A | 1 | |
| VCL | Vinculin | 1 | yes |
| VDAC3 | Voltage-dependent anion-selective channel protein 3 | 1 | |

**S3: Enriched Cellular Compartments based on STRING database in the case of N-BAP-biotin-rhTG2 associated proteins (5≤ detected peptide count, n=3)**

| #term ID | term description | false discovery rate | matching proteins in network (labels) |
|---|---|---|---|
| GO:0005925 | Focal adhesion | 9.72e-05 | RALA,MCAM,GNB2,PLEC,FLII,RPS18,YES1,CD59 |
| GO:0031982 | Vesicle | 9.72e-05 | RALA,COPB1,NT5E,GNB2,GNAI2,PLEC,GANAB,FN1,DYNC1H1,AP2A1,CDC42BPB,GNB1,XRCC5,RPS18,DDOST,CAND1,YES1,CD59 |
| GO:0070062 | Extracellular exosome | 9.72e-05 | RALA,NT5E,GNB2,GNAI2,PLEC,GANAB,FN1,DYNC1H1,CDC42BPB,GNB1,RPS18,CAND1,YES1,CD59 |
| GO:0098552 | Side of membrane | 0.00022 | NT5E,MCAM,GNB2,GNAI2,AP2A1,GNB1,YES1,CD59 |
| GO:0005615 | Extracellular space | 0.00052 | RALA,NT5E,MCAM,GNB2,GNAI2,PLEC,GANAB,FN1,DYNC1H1,CDC42BPB,GNB1,RPS18,CAND1,YES1,CD59 |
| GO:0009898 | Cytoplasmic side of plasma membrane | 0.00067 | GNB2,GNAI2,AP2A1,GNB1,YES1 |
| GO:0030054 | Cell junction | 0.0010 | RALA,MCAM,GNB2,GNAI2,PLEC,FLII,AP2A1,CDC42BPB,GNB1,RPS18,YES1,CD59 |
| GO:0005576 | Extracellular region | 0.0015 | RALA,NT5E,MCAM,GNB2,GNAI2,PLEC,GANAB,FN1,DYNC1H1,CDC42BPB,GNB1,XRCC5,RPS18,CAND1,YES1,CD59 |
| GO:0031234 | Extrinsic component of cytoplasmic side of plasma mem | 0.0016 | GNB2,GNAI2,GNB1,YES1 |
| GO:0005834 | Heterotrimeric G-protein complex | 0.0023 | GNB2,GNAI2,GNB1 |
| GO:0070418 | DNA-dependent protein kinase complex | 0.0030 | PRKDC,XRCC5 |
| GO:0070161 | Anchoring junction | 0.0036 | RALA,MCAM,GNB2,PLEC,FLII,CDC42BPB,RPS18,YES1,CD59 |
| GO:0070419 | Nonhomologous end joining complex | 0.0093 | PRKDC,XRCC5 |
| GO:0030141 | Secretory granule | 0.0105 | COPB1,FN1,DYNC1H1,XRCC5,DDOST,CAND1,CD59 |
| GO:0016020 | Membrane | 0.0134 | RALA,COPB1,NT5E,MCAM,TMPO,GNB2,GNAI2,PRKDC,PLEC,GANAB,FN1,DYNC1H1,AP2A1,CDC42BPB,PARP1,GNB1,XRCC5,RPS18,DDOST,CAND1,YES1,CD59 |
| GO:0005829 | Cytosol | 0.0274 | COPB1,NT5E,FMNL2,GNB2,GNAI2,PRKDC,PLEC,FLII,DYNC1H1,AP2A1,CDC42BPB,GNB1,XRCC5,RPS18,CAND1,YES1 |
| GO:0005737 | Cytoplasm | 0.0336 | RALA,PDIA2,COPB1,NT5E,TMPO,FMNL2,GNB2,GNAI2,PRKDC,PLEC,FLII,GANAB,FN1,DYNC1H1,AP2A1,CDC42BPB,PARP1,GNB1,XRCC5,RPS18,DDOST,CAND1,YES1,CD59 |
| GO:0043227 | Membrane-bounded organelle | 0.0336 | RALA,PDIA2,COPB1,NT5E,MCAM,TMPO,SSRP1,GNB2,GNAI2,PRKDC,PLEC,FLII,GANAB,FN1,DYNC1H1,AP2A1,CDC42BPB,PARP1,GNB1,XRCC5,RPS18,DDOST,CAND1,YES1,CD59 |
| GO:0043229 | Intracellular organelle | 0.0336 | RALA,PDIA2,COPB1,NT5E,MCAM,TMPO,SSRP1,GNB2,GNAI2,PRKDC,PLEC,FLII,GANAB,FN1,DYNC1H1,AP2A1,CDC42BPB,PARP1,GNB1,XRCC5,RPS18,DDOST,CAND1,YES1,CD59 |
| GO:0043231 | Intracellular membrane-bounded organelle | 0.0366 | RALA,PDIA2,COPB1,NT5E,MCAM,TMPO,SSRP1,GNB2,GNAI2,PRKDC,PLEC,FLII,GANAB,FN1,DYNC1H1,AP2A1,PARP1,GNB1,XRCC5,RPS18,DDOST,CAND1,YES1,CD59 |
| GO:1902494 | Catalytic complex | 0.0430 | GNB2,GNAI2,PRKDC,DYNC1H1,GNB1,XRCC5,DDOST,CAND1 |
| GO:0005622 | Intracellular anatomical structure | 0.0475 | RALA,PDIA2,COPB1,NT5E,MCAM,TMPO,SSRP1,FMNL2,GNB2,GNAI2,PRKDC,PLEC,FLII,GANAB,FN1,DYNC1H1,AP2A1,CDC42BPB,PARP1,GNB1,XRCC5,RPS18,DDOST,CAND1,YES1,CD59 |
| GO:0005793 | Endoplasmic reticulum-Golgi intermediate compartmen | 0.0490 | COPB1,FN1,CD59 |

**S4: Enriched GO Molecular Functions based on the STRING database in the case of N-BAP-biotin-rhTG2 associated proteins (all hits, n=3)**

| #term ID | term description | false discovery rate | matching proteins in network |
|---|---|---|---|
| GO:0005515 | Protein binding | 2.01e-12 | RALA,UQCRC1,GLG1,VCL,FBL,RAC2,LMAN1,SRSF1,LRPPRC,IPO5,ACIN1,PRDX1,CAPZA1,NOP58,TOP2B,RAB10,RAB7A,TCIRG1,ACAT1,TMPO,ESYT1,PFKL,SSRP1,RHOC,STOM,FMNL2,ITGA5,CCT3,H2AZ1,MAP1B,VPS35,TMED10,FASN,GNB2,KIF5B,QARS1,DST,GNAI2,PRKDC,ITGA3,TUBB3,SMC1A,PLEC,FLII,H2AC14,TMED9,RAB11B,TNPO1,CAV1,PPP1CC,PRDX6,CTSB,FN1,SPRED2,RPS26,DYNC1H1,IGF2R,TPM1,AP2A1,PCBP2,CAPZA2,CDC42BPB,PARP1,MGST3,COPA,IFI16,RAP1A,EMD,RPL5,GNG12,SRSF3,GSN,CNTRL,COL5A2,GNB1,ATAD3A,DNAJA1,EIF3CL,NOP56,GARS1,HDLBP,XRCC5,MYO1B,EFEMP1,TGFB1I1,VDAC1,YWHAZ,CTNND1,FKBP1A,SUN2,FHL2,UBA52,CAPZB,UNC45A,ERLIN1,PDCD6IP,BRCA1,RACK1,TJP2,PHB2,CAND1,TIMM50,RAN,HEXA,YES1,TTN,H2BC17,HYOU1,PDLIM5,AP2B1,CDC42,STING1,CD59,ESYT2 |
| GO:0045296 | Cadherin binding | 2.01e-12 | VCL,PRDX1,CAPZA1,RAB10,TMPO,FMNL2,FASN,KIF5B,PLEC,RAB11B,PRDX6,RPS26,EMD,NOP56,HDLBP,MYO1B,YWHAZ,CTNND1,CAPZB,UNC45A,RACK1,TJP2,RAN,PDLIM5,ESYT2 |
| GO:0050839 | Cell adhesion molecule binding | 2.56e-11 | VCL,PRDX1,CAPZA1,RAB10,TMPO,FMNL2,ITGA5,FASN,KIF5B,DST,ITGA3,PLEC,RAB11B,PRDX6,FN1,RPS26,EMD,NOP56,HDLBP,MYO1B,YWHAZ,CTNND1,CAPZB,UNC45A,RACK1,TJP2,RAN,PDLIM5,ESYT2 |
| GO:0019899 | Enzyme binding | 6.11e-11 | RALA,UQCRC1,VCL,FBL,RAC2,SRSF1,LRPPRC,IPO5,ACIN1,NOP58,TOP2B,RAB7A,TCIRG1,ACAT1,PFKL,RHOC,STOM,FMNL2,GNB2,QARS1,PRKDC,ITGA3,TNPO1,CAV1,PPP1CC,PRDX6,FN1,SPRED2,IGF2R,AP2A1,PCBP2,CDC42BPB,PARP1,RAP1A,RPL5,SRSF3,GSN,GNB1,DNAJA1,NOP56,XRCC5,VDAC1,YWHAZ,CTNND1,UBA52,ERLIN1,BRCA1,RACK1,TJP2,YES1,TTN,PDLIM5,CDC42,STING1 |
| GO:0044877 | Protein-containing complex binding | 1.05e-08 | UQCRC1,VCL,FBL,CAPZA1,NOP58,RAB7A,SSRP1,FMNL2,ITGA5,H2AZ1,MAP1B,GNB2,KIF5B,DST,GNAI2,ITGA3,SMC1A,FLII,CAV1,PPP1CC,H1-0,CTSB,FN1,IGF2R,TPM1,CAPZA2,CDC42BPB,RAP1A,GSN,GNB1,XRCC5,MYO1B,FKBP1A,CAPZB,HNRNPAB,RACK1,TIMM50,TTN |
| GO:0005488 | Binding | 1.09e-08 | RALA,UQCRC1,GLG1,DLD,VCL,PDIA2,FBL,GALK1,RAC2,LMAN1,NT5E,SRSF1,LRPPRC,IPO5,MARS1,ACIN1,PRDX1,CAPZA1,TUBB4A,NOP58,TOP2B,RAB10,RAB7A,TCIRG1,ACAT1,TMPO,ESYT1,UQCRC2,PFKL,ATP6V1B2,SSRP1,PLOD2,RHOC,STOM,FMNL2,ITGA5,CCT3,H2AZ1,MAP1B,VPS35,HSPA4,TMED10,FASN,GNB2,KIF5B,MCM7,QARS1,DST,ARHGAP32,GNAI2,PRKDC,ITGA3,TUBB3,MMP1,SMC1A,PLEC,ZYX,FLII,PLAG1,H2AC14,TMED9,RAB11B,TNPO1,CAV1,GANAB,PPP1CC,PRDX6,H1-0,CTSB,FN1,SPRED2,RPS26,DYNC1H1,ATP5F1C,IGF2R,TPM1,AP2A1,PCBP2,CYB5R3,CAPZA2,CDC42BPB,PARP1,MGST3,COPA,IFI16,EIF3A,RAP1A,EMD,RPL5,GNG12,SRSF3,GSN,CNTRL,COL5A2,GNB1,ATAD3A,DNAJA1,SLTM,EIF3CL,NOP56,GARS1,HDLBP,XRCC5,MYO1B,G3BP1,LARS1,EFEMP1,TGFB1I1,VDAC1,YWHAZ,KHSRP,CTNND1,FKBP1A,SUN2,FHL2,UBA52,UBAP2L,RPS18,CAPZB,PAPSS2,UNC45A,ERLIN1,PDCD6IP,BRCA1,HNRNPAB,RACK1,VDAC3,CAPN1,TJP2,PHB2,CAND1,TIMM50,RAN,HEXA,YES1,ACLY,TTN,H2BC17,HYOU1,PDLIM5,AP2B1,CLEC19A,PHGDH,HK1,CDC42,STING1,CD59,ESYT2 |
| GO:0003723 | RNA binding | 3.71e-08 | FBL,SRSF1,LRPPRC,IPO5,MARS1,ACIN1,PRDX1,NOP58,SSRP1,CCT3,FASN,PRKDC,SMC1A,PLEC,ZYX,TNPO1,GANAB,PPP1CC,H1-0,RPS26,DYNC1H1,ATP5F1C,PCBP2,PARP1,IFI16,EIF3A,RPL5,SRSF3,SLTM,EIF3CL,NOP56,HDLBP,XRCC5,G3BP1,YWHAZ,KHSRP,UBAP2L,RPS18,BRCA1,HNRNPAB,RACK1,TIMM50,RAN |
| GO:0036094 | Small molecule binding | 3.99e-08 | RALA,DLD,GALK1,RAC2,LMAN1,NT5E,MARS1,TUBB4A,TOP2B,RAB10,RAB7A,ACAT1,PFKL,ATP6V1B2,PLOD2,RHOC,CCT3,HSPA4,FASN,KIF5B,MCM7,QARS1,GNAI2,PRKDC,TUBB3,SMC1A,RAB11B,CAV1,DYNC1H1,IGF2R,CYB5R3,CDC42BPB,PARP1,RAP1A,ATAD3A,DNAJA1,GARS1,XRCC5,MYO1B,G3BP1,LARS1,VDAC1,PAPSS2,ERLIN1,VDAC3,RAN,YES1,ACLY,TTN,HYOU1,PHGDH,HK1,CDC42,STING1 |
| GO:0097159 | Organic cyclic compound binding | 5.92e-08 | RALA,DLD,PDIA2,FBL,GALK1,RAC2,NT5E,SRSF1,LRPPRC,IPO5,MARS1,ACIN1,PRDX1,TUBB4A,NOP58,TOP2B,RAB10,RAB7A,ACAT1,TMPO,PFKL,ATP6V1B2,SSRP1,PLOD2,RHOC,CCT3,H2AZ1,HSPA4,FASN,KIF5B,MCM7,QARS1,GNAI2,PRKDC,TUBB3,SMC1A,PLEC,ZYX,PLAG1,H2AC14,RAB11B,TNPO1,CAV1,GANAB,PPP1CC,H1-0,RPS26,DYNC1H1,ATP5F1C,PCBP2,CYB5R3,CDC42BPB,PARP1,IFI16,EIF3A,RAP1A,RPL5,SRSF3,ATAD3A,DNAJA1,SLTM,EIF3CL,NOP56,GARS1,HDLBP,XRCC5,MYO1B,G3BP1,LARS1,VDAC1,YWHAZ,KHSRP,FKBP1A,UBAP2L,RPS18,PAPSS2,ERLIN1,BRCA1,HNRNPAB,RACK1,VDAC3,TIMM50,RAN,YES1,ACLY,TTN,H2BC17,HYOU1,PHGDH,HK1,CDC42,STING1 |
| GO:0000166 | Nucleotide binding | 7.21e-07 | RALA,DLD,GALK1,RAC2,NT5E,MARS1,TUBB4A,TOP2B,RAB10,RAB7A,ACAT1,PFKL,ATP6V1B2,RHOC,CCT3,HSPA4,KIF5B,MCM7,QARS1,GNAI2,PRKDC,TUBB3,SMC1A,RAB11B,DYNC1H1,CYB5R3,CDC42BPB,PARP1,RAP1A,ATAD3A,DNAJA1,GARS1,XRCC5,MYO1B,G3BP1,LARS1,PAPSS2,VDAC3,RAN,YES1,ACLY,TTN,HYOU1,PHGDH,HK1,CDC42,STING1 |
| GO:1901363 | Heterocyclic compound binding | 1.09e-06 | RALA,DLD,FBL,GALK1,RAC2,NT5E,SRSF1,LRPPRC,IPO5,MARS1,ACIN1,PRDX1,TUBB4A,NOP58,TOP2B,RAB10,RAB7A,ACAT1,TMPO,PFKL,ATP6V1B2,SSRP1,PLOD2,RHOC,CCT3,H2AZ1,HSPA4,FASN,KIF5B,MCM7,QARS1,GNAI2,PRKDC,TUBB3,SMC1A,PLEC,ZYX,PLAG1,H2AC14,RAB11B,TNPO1,GANAB,PPP1CC,H1-0,RPS26,DYNC1H1,ATP5F1C,PCBP2,CYB5R3,CDC42BPB,PARP1,IFI16,EIF3A,RAP1A,RPL5,SRSF3,ATAD3A,DNAJA1,SLTM,EIF3CL,NOP56,GARS1,HDLBP,XRCC5,MYO1B,G3BP1,LARS1,YWHAZ,KHSRP,FKBP1A,UBAP2L,RPS18,PAPSS2,BRCA1,HNRNPAB,RACK1,VDAC3,TIMM50,RAN,YES1,ACLY,TTN,H2BC17,HYOU1,PHGDH,HK1,CDC42,STING1 |

| | | | |
|---|---|---|---|
| GO:0032555 | Purine ribonucleotide binding | 1.02e-05 | RALA,GALK1,RAC2,MARS1,TUBB4A,TOP2B,RAB10,RAB7A,ACAT1,PFKL,ATP6V1B2,RHOC,CCT3,HSPA4,KIF5B,MCM7,QARS1,GNAI2,PRKDC,TUBB3,SMC1A,RAB11B,DYNC1H1,CDC42BPB,RAP1A,ATAD3A,DNAJA1,GARS1,XRCC5,MYO1B,G3BP1,LARS1,PAPSS2,RAN,YES1,ACLY,TTN,HYOU1,HK1,CDC42,STING1 |
| GO:0043168 | Anion binding | 1.15e-05 | RALA,DLD,GALK1,RAC2,MARS1,TUBB4A,TOP2B,RAB10,RAB7A,ACAT1,PFKL,ATP6V1B2,PLOD2,RHOC,CCT3,HSPA4,KIF5B,MCM7,QARS1,GNAI2,PRKDC,TUBB3,SMC1A,FLII,RAB11B,DYNC1H1,IGF2R,CYB5R3,CDC42BPB,RAP1A,GSN,ATAD3A,DNAJA1,GARS1,XRCC5,MYO1B,G3BP1,LARS1,PAPSS2,RAN,YES1,ACLY,TTN,HYOU1,HK1,CDC42,STING1 |
| GO:0035639 | Purine ribonucleoside triphosphate binding | 2.62e-05 | RALA,GALK1,RAC2,MARS1,TUBB4A,TOP2B,RAB10,RAB7A,PFKL,ATP6V1B2,RHOC,CCT3,HSPA4,KIF5B,MCM7,QARS1,GNAI2,PRKDC,TUBB3,SMC1A,RAB11B,DYNC1H1,CDC42BPB,RAP1A,ATAD3A,DNAJA1,GARS1,XRCC5,MYO1B,G3BP1,LARS1,PAPSS2,RAN,YES1,ACLY,TTN,HYOU1,HK1,CDC42 |
| GO:0031625 | Ubiquitin protein ligase binding | 0.00012 | RALA,UQCRC1,VCL,LRPPRC,PRDX6,PCBP2,RPL5,DNAJA1,XRCC5,YWHAZ,UBA52,ERLIN1,BRCA1,STING1 |
| GO:0097367 | Carbohydrate derivative binding | 0.00012 | RALA,GALK1,RAC2,MARS1,TUBB4A,TOP2B,RAB10,RAB7A,ACAT1,PFKL,ATP6V1B2,RHOC,CCT3,HSPA4,KIF5B,MCM7,QARS1,GNAI2,PRKDC,TUBB3,SMC1A,RAB11B,CTSB,FN1,DYNC1H1,CDC42BPB,RAP1A,ATAD3A,DNAJA1,GARS1,XRCC5,MYO1B,G3BP1,LARS1,PAPSS2,RAN,YES1,ACLY,TTN,HYOU1,HK1,CDC42,STING1 |
| GO:0005198 | Structural molecule activity | 0.00025 | VCL,COPB1,TUBB4A,H2AZ1,MAP1B,DST,NUP93,TUBB3,PLEC,H2AC14,H1-0,FN1,RPS26,TPM1,COPA,EIF3A,RPL5,COL5A2,UBA52,RPS18,TTN,H2BC17 |
| GO:0003925 | G protein activity | 0.00053 | RALA,RAB10,RAB7A,RAB11B,RAP1A,CDC42 |
| GO:0016462 | Pyrophosphatase activity | 0.00074 | RALA,RAC2,ACIN1,RAB10,RAB7A,RHOC,CCT3,GNB2,KIF5B,MCM7,GNAI2,SMC1A,RAB11B,RAP1A,GNB1,ATAD3A,GARS1,G3BP1,RAN,CDC42 |
| GO:0017111 | Nucleoside-triphosphatase activity | 0.00077 | RALA,RAC2,ACIN1,RAB10,RAB7A,RHOC,CCT3,GNB2,KIF5B,MCM7,GNAI2,SMC1A,RAB11B,RAP1A,GNB1,ATAD3A,G3BP1,RAN,CDC42 |
| GO:0003779 | Actin binding | 0.0018 | VCL,CAPZA1,FMNL2,MAP1B,DST,PLEC,FLII,TPM1,CAPZA2,EMD,GSN,MYO1B,CAPZB,TTN,PDLIM5 |
| GO:0019900 | Kinase binding | 0.0028 | RAC2,TOP2B,PFKL,RHOC,QARS1,CAV1,PPP1CC,SPRED2,AP2A1,PARP1,GSN,VDAC1,YWHAZ,CTNND1,RACK1,TJP2,TTN,PDLIM5,CDC42,STING1 |
| GO:0008092 | Cytoskeletal protein binding | 0.0031 | RALA,VCL,LRPPRC,CAPZA1,RAB10,FMNL2,MAP1B,KIF5B,DST,PLEC,FLII,RAB11B,TPM1,CAPZA2,EMD,GSN,CNTRL,MYO1B,SUN2,CAPZB,BRCA1,TTN,PDLIM5 |
| GO:0051015 | Actin filament binding | 0.0034 | VCL,CAPZA1,FMNL2,FLII,TPM1,CAPZA2,GSN,MYO1B,CAPZB,TTN |
| GO:0003924 | GTPase activity | 0.0039 | RALA,RAC2,RAB10,RAB7A,RHOC,GNB2,GNAI2,RAB11B,RAP1A,GNB1,RAN,CDC42 |
| GO:0019901 | Protein kinase binding | 0.0062 | RAC2,TOP2B,RHOC,QARS1,CAV1,PPP1CC,SPRED2,AP2A1,PARP1,VDAC1,YWHAZ,CTNND1,RACK1,TJP2,TTN,PDLIM5,CDC42,STING1 |
| GO:0019003 | GDP binding | 0.0078 | RALA,RAB10,RAB7A,RAB11B,RAP1A,RAN |
| GO:0032559 | Adenyl ribonucleotide binding | 0.0081 | GALK1,MARS1,TOP2B,ACAT1,PFKL,ATP6V1B2,CCT3,HSPA4,KIF5B,MCM7,QARS1,PRKDC,SMC1A,DYNC1H1,CDC42BPB,ATAD3A,DNAJA1,GARS1,XRCC5,MYO1B,G3BP1,LARS1,PAPSS2,YES1,ACLY,TTN,HYOU1,HK1,STING1 |
| GO:0032561 | Guanyl ribonucleotide binding | 0.0081 | RALA,RAC2,TUBB4A,RAB10,RAB7A,RHOC,GNAI2,TUBB3,RAB11B,RAP1A,RAN,CDC42,STING1 |
| GO:0048029 | Monosaccharide binding | 0.0081 | GALK1,LMAN1,PFKL,PLOD2,IGF2R,HK1 |
| GO:0140657 | ATP-dependent activity | 0.0104 | ACIN1,TOP2B,TCIRG1,ATP6V1B2,CCT3,HSPA4,KIF5B,MCM7,SMC1A,DYNC1H1,ATAD3A,XRCC5,MYO1B,G3BP1,HYOU1 |
| GO:0005525 | GTP binding | 0.0166 | RALA,RAC2,TUBB4A,RAB10,RAB7A,RHOC,GNAI2,TUBB3,RAB11B,RAP1A,RAN,CDC42 |
| GO:0005524 | ATP binding | 0.0219 | GALK1,MARS1,TOP2B,PFKL,ATP6V1B2,CCT3,HSPA4,KIF5B,MCM7,QARS1,PRKDC,SMC1A,DYNC1H1,CDC42BPB,ATAD3A,DNAJA1,GARS1,XRCC5,MYO1B,G3BP1,LARS1,PAPSS2,YES1,ACLY,TTN,HYOU1,HK1 |
| GO:0030515 | snoRNA binding | 0.0247 | NOP58,PRKDC,NOP56,XRCC5 |
| GO:0046982 | Protein heterodimerization activity | 0.0452 | TOP2B,H2AZ1,ITGA3,SMC1A,H2AC14,CAV1,TPM1,PHB2,RAN,HEXA,H2BC17 |
| GO:0004819 | glutamine-tRNA ligase activity | 0.0473 | QARS1,LARS1 |

**S5: Identified nonspecifically bound proteins to mouse IgG agarose resin from HUVEC extract (controls, subtracted from sample hits, n=3)**

| Gene Name | Name of proteins |
|---|---|
| ACTB | Actin, cytoplasmic 1 |
| ACTBL2 | Beta-actin-like protein 2 |
| ACTG1 | Actin, cytoplasmic 2 |
| ACTN4 | Alpha-actinin-4 |
| ANXA1 | Annexin A1 |
| ANXA2 | Annexin A2 |
| ARHGDIA | Rho GDP-dissociation inhibitor 1 |
| ATP5F1A | ATP synthase subunit alpha, mitochondrial |
| ATP5F1B | ATP synthase subunit beta, mitochondrial |
| BRCA2 | Breast cancer type 2 susceptibility protein |
| CALM1 | Calmodulin-1 |
| CAPRIN1 | Caprin-1 |
| CAPZA2 | F-actin-capping protein subunit alpha-2 |
| CARM1 | Histone-arginine methyltransferase CARM1 |
| CCAR2 | Cell cycle and apoptosis regulator protein 2 |
| CFL1 | Cofilin-1 |
| CLIC1 | Chloride intracellular channel protein 1 |
| CLTC | Clathrin heavy chain 1 |
| CNGA1 | cGMP-gated cation channel alpha-1 |
| CNN2 | Calponin-2 |
| COLGALT1 | Procollagen galactosyltransferase 1 |
| CORO1C | Coronin-1C |
| CPSF7 | Cleavage and polyadenylation specificity factor subunit 7 |
| CRIP2 | Cysteine-rich protein 2 |
| CTNNA1 | Catenin alpha-1 |
| CTTN | Src substrate cortactin |
| DBN1 | Drebrin |
| DDX1 | ATP-dependent RNA helicase DDX1 |
| DDX17 | Probable ATP-dependent RNA helicase DDX17 |
| DDX3X | ATP-dependent RNA helicase DDX3X |
| DDX5 | Probable ATP-dependent RNA helicase DDX5 |
| DERPC | Probable ATP-dependent RNA helicase DDX5 |
| DHX9 | ATP-dependent RNA helicase A |
| EEF1A1 | Elongation factor 1-alpha 1 |
| EEF2 | Elongation factor 2 |
| EIF4A1 | Eukaryotic initiation factor 4A-I |
| EIF4A2 | Eukaryotic initiation factor 4A-II |
| EIF4G1 | Eukaryotic translation initiation factor 4 gamma 1 |
| ELAVL1 | ELAV-like protein 1 |
| EMD | Emerin |
| EWSR1 | RNA-binding protein EWS |
| FAM120A | Constitutive coactivator of PPAR-gamma-like protein 1 |
| FARSA | Phenylalanine--tRNA ligase alpha subunit |
| FLNA | Filamin-A |
| FLNB | Filamin-B |
| FLNC | Filamin-C |
| FN1 | Fibronectin |
| FUS | RNA-binding protein FUS |
| G3BP1 | Ras GTPase-activating protein-binding protein 1 |
| GAPDH | Glyceraldehyde-3-phosphate dehydrogenase |
| GSTP1 | Glutathione S-transferase P |
| H1-2 | Histone H1.2 |
| H1-3 | Histone H1.3 |
| H1-4 | Histone H1.4 |
| H1-5 | Histone H1.5 |
| H2AC11 | Histone H2A type 1 |
| H2AC20 | Histone H2A type 2-C |
| H2AC4 | Histone H2A type 1-B/E |
| H2AJ | Histone H2A.J |
| H2BC12 | Histone H2B type 1-K |
| H2BC17 | Histone H2B type 1-O |
| H2BC18 | Histone H2B type 1-O |
| H2BC9 | Histone H2B type 1-H |
| H3C1 | Histone H3.1 |
| H4C1 | Histone H4 |
| HMGA1 | High mobility group protein HMG-I/HMG-Y |
| HNRNPA1 | Heterogeneous nuclear ribonucleoprotein A1 |
| HNRNPA2B1 | Heterogeneous nuclear ribonucleoproteins A2/B1 |
| HNRNPA3 | Heterogeneous nuclear ribonucleoprotein A3 |
| HNRNPAB | Isoform 2 of Heterogeneous nuclear ribonucleoprotein A/B |

| | |
|---|---|
| HNRNPC | Heterogeneous nuclear ribonucleoproteins C1/C2 |
| HNRNPD | Heterogeneous nuclear ribonucleoprotein D0 |
| HNRNPDL | Heterogeneous nuclear ribonucleoprotein D-like |
| HNRNPF | Heterogeneous nuclear ribonucleoprotein F |
| HNRNPH1 | Heterogeneous nuclear ribonucleoprotein H |
| HNRNPH3 | Heterogeneous nuclear ribonucleoprotein H3 |
| HNRNPK | Heterogeneous nuclear ribonucleoprotein K |
| HNRNPL | Heterogeneous nuclear ribonucleoprotein L |
| HNRNPM | Heterogeneous nuclear ribonucleoprotein M |
| HNRNPR | Heterogeneous nuclear ribonucleoprotein R |
| HNRNPU | Heterogeneous nuclear ribonucleoprotein U |
| HNRNPUL1 | Heterogeneous nuclear ribonucleoprotein U-like protein 1 |
| HNRNPUL2 | Heterogeneous nuclear ribonucleoprotein U-like protein 2 |
| HSD17B4 | Peroxisomal multifunctional enzyme type 2 |
| HSP90AB1 | Heat shock protein HSP 90-beta |
| HSP90B1 | Endoplasmin |
| HSPA5 | Endoplasmic reticulum chaperone BiP |
| HSPA8 | Heat shock cognate 71 kDa protein |
| HSPB1 | Heat shock protein beta-1 |
| HSPD1 | 60 kDa heat shock protein, mitochondrial |
| HSPG2 | Basement membrane-specific heparan sulfate proteoglycan core protein |
| IGF2BP2 | Insulin-like growth factor 2 mRNA-binding protein 2 |
| IGF2BP3 | Insulin-like growth factor 2 mRNA-binding protein 3 |
| IGKV2D-26 | Immunoglobulin kappa variable 2D-26 |
| ILF3 | Interleukin enhancer-binding factor 3 |
| KHDRBS1 | KH domain-containing, RNA-binding, signal transduction-associated protein 1 |
| KHSRP | Far upstream element-binding protein 2 |
| KRT1 | Keratin, type II cytoskeletal 1 |
| KRT10 | Keratin, type II cytoskeletal 10 |
| KRT9 | Keratin, type II cytoskeletal 9 |
| LDHA | L-lactate dehydrogenase A |
| LETM1 | Mitochondrial proton/calcium exchanger protein |
| LGALS1 | Galectin-1 |
| LMNA | Prelamin-A/C |
| LMO7 | Isoform 3 of LIM domain only protein 7 |
| MACROH2A1 | Core histone macro-H2A.1 |
| MATR3 | Matrin-3 |
| MRPL28 | 39S ribosomal protein L28, mitochondrial |
| MVP | Major vault protein |
| MYH9 | Myosin-9 |
| MYL12B | Myosin regulatory light chain 12B |
| MYO1C | Unconventional myosin-Ic |
| MYO1D | Unconventional myosin-Id |
| MYO6 | Isoform 6 of Unconventional myosin-VI |
| NCL | Nucleolin |
| NONO | Non-POU domain-containing octamer-binding protein |
| NPM1 | Nucleophosmin |
| NUDT21 | Cleavage and polyadenylation specificity factor subunit 5 |
| PABPC1 | Polyadenylate-binding protein 1 |
| PABPC4 | Isoform 2 of Polyadenylate-binding protein 4 |
| PABPN1 | Polyadenylate-binding protein 2 |
| PDIA6 | Protein disulfide-isomerase A6 |
| PDLIM7 | PDZ and LIM domain protein 7 |
| PFN1 | Profilin-1 |
| PGK1 | Phosphoglycerate kinase 1 |
| PHB1 | Prohibitin 1 |
| PLEC | Plectin |
| PLS3 | Plastin-3 |
| PPIA | Peptidyl-prolyl cis-trans isomerase A |
| PRDX1 | Peroxiredoxin-1 |
| PRDX6 | Peroxiredoxin-6 |
| PSPC1 | Paraspeckle component 1 |
| PTBP1 | Polypyrimidine tract-binding protein 1 |
| RASIP1 | Ras-interacting protein 1 |
| RBM14 | RNA-binding protein 14 |
| RBMS1 | RNA-binding motif, single-stranded-interacting protein 1 |
| RBMS2 | RNA-binding motif, single-stranded-interacting protein 2 |
| RBMX | RNA-binding motif protein, X chromosome |
| RPL15 | 60S ribosomal protein L15 |
| RPL18A | 60S ribosomal protein L18a |
| RPL19 | 60S ribosomal protein L19 |
| RPL24 | 60S ribosomal protein L24 |
| RPL4 | 60S ribosomal protein L4 |
| RPL6 | 60S ribosomal protein L6 |

| | |
|---|---|
| RPL7 | 60S ribosomal protein L7a |
| RPL7A | 60S ribosomal protein L7a |
| RPS11 | 40S ribosomal protein S11 |
| RPS12 | 40S ribosomal protein S12 |
| RPS18 | 40S ribosomal protein S18 |
| RPS20 | 40S ribosomal protein S20 |
| RPS27A | Ubiquitin-40S ribosomal protein S27a |
| RPS5 | 40S ribosomal protein S5 |
| RPS6 | 40S ribosomal protein S6 |
| RPS8 | 40S ribosomal protein S8 |
| RPS9 | 40S ribosomal protein S9 |
| RTCB | RNA-splicing ligase RtcB homolog |
| SAFB | Scaffold attachment factor B1 |
| SEC23A | Protein transport protein Sec23A |
| SFPQ | Splicing factor, proline- and glutamine-rich |
| SH3GLB2 | Endophilin-B2 |
| SLC25A3 | Isoform B of Phosphate carrier protein, mitochondrial |
| SLC25A5 | ADP/ATP translocase 2 |
| SLC25A6 | ADP/ATP translocase 3 |
| SNX9 | Sorting nexin-9 |
| SPCS2 | Signal peptidase complex subunit 2 |
| SPECC1L | Cytospin-A |
| SPTAN1 | Spectrin alpha chain, non-erythrocytic 1 |
| SPTBN1 | Spectrin beta chain, non-erythrocytic 1 |
| SRSF10 | Serine/arginine-rich splicing factor 10 |
| STK31 | Serine/threonine-protein kinase 31 |
| SYNCRIP | Isoform 3 of Heterogeneous nuclear ribonucleoprotein Q |
| TAF15 | TATA-binding protein-associated factor 2N |
| TARDBP | Isoform 2 of TAR DNA-binding protein 43 |
| TFG | Protein TFG |
| TKT | Transketolase |
| TPM1 | Tropomyosin alpha-1 chain |
| TUBA1B | Tubulin alpha-1B |
| TUBB | Tubulin beta chain |
| TUBB4B | Tubulin beta-4B chain |
| TUBB6 | Tubulin beta-6 chain |
| UBA1 | Ubiquitin-like modifier-activating enzyme 1 |
| UBAP2L | Ubiquitin-associated protein 2-like |
| UBB | Polyubiquitin-B |
| VCP | Transitional endoplasmic reticulum ATPase |
| VIM | Vimentin |
| XRCC6 | X-ray repair cross-complementing protein 6 |
| XRN2 | 5'-3' exoribonuclease 2 |
| ZNF185 | Zinc finger protein 185 |
| ZYX | Zyxin |

**S6: Identified 3F-TG2-associated proteins in HUVEC using anti-Flag M2 agarose resin for the CoIP**

| Gene Name | Name of potential TG2 interacting partners | Identified unique peptide count (n=3) | Known TG2 interacting partner protein |
|---|---|---|---|
| TGM2 | Protein-glutamine gamma-glutamyltransferase 2 | 170 | |
| CKAP4 | Cytoskeleton-associated protein 4 | 91 | |
| THRAP3 | Thyroid hormone receptor-associated protein 3 | 45 | |
| ACTN1 | Alpha-actinin-1 | 39 | |
| RPN1 | Dolichyl-diphosphooligosaccharide--protein glycosyltransferase subunit 1 | 36 | |
| EEF1G | Elongation factor 1-gamma | 29 | |
| MAP1B | Microtubule-associated protein 1B | 28 | |
| MSN | Moesin | 24 | |
| NT5E | 5'-nucleotidase | 22 | |
| ANXA5 | Annexin A5 | 21 | |
| TLN1 | Talin-1 | 20 | yes |
| TUBA1A | Tubulin alpha-1A chain | 19 | yes |
| PHB2 | Prohibitin-2 | 18 | |
| ANPEP | Aminopeptidase N | 17 | |
| ERLIN1 | Erlin-1 | 17 | |
| TRPV2 | Transient receptor potential cation channel subfamily V member 2 | 17 | |
| CAVIN1 | Caveolae-associated protein 1 | 16 | |
| GNAI2 | Guanine nucleotide-binding protein G(i) subunit alpha-2 | 16 | |
| RPS3A | 40S ribosomal protein S3a | 16 | |
| PDIA3 | Protein disulfide-isomerase A3 | 15 | |
| PRMT5 | Protein arginine N-methyltransferase 5 | 15 | |
| IGKV2D-29 | Immunoglobulin kappa variable 2D-29 | 14 | |
| RRBP1 | Ribosome-binding protein 1 | 14 | |
| ATP1A1 | Sodium/potassium-transporting ATPase subunit alpha-1 | 13 | |
| DPP7 | Dipeptidyl peptidase 2 | 13 | |
| FLOT2 | Flotillin-2 | 13 | |
| RBM10 | RNA-binding protein 10 | 13 | |
| RPL3 | 60S ribosomal protein L3 | 13 | |
| CD59 | CD59 glycoprotein | 12 | |
| H2BC4 | Histone H2B type 1-C/E/F/G/I | 12 | yes |
| MYL6 | Myosin light polypeptide 6 | 12 | |
| RPL18 | 60S ribosomal protein L18 | 12 | |
| RPL8 | 60S ribosomal protein L8 | 12 | |
| RPN2 | Dolichyl-diphosphooligosaccharide--protein glycosyltransferase subunit 2 | 12 | |
| DDOST | Dolichyl-diphosphooligosaccharide--protein glycosyltransferase 48 kDa subunit | 11 | |
| JAK1 | Tyrosine-protein kinase JAK1 | 11 | |
| MBOAT7 | Lysophospholipid acyltransferase 7 | 11 | |
| RPL13 | 60S ribosomal protein L13 | 11 | |
| RPSA | 40S ribosomal protein Sa | 11 | |
| AHNAK | Neuroblast differentiation-associated protein AHNAK | 10 | |
| AP2A1 | AP-2 complex subunit alpha-1 | 10 | |
| HSP90AA1 | Heat shock protein HSP 90-alpha | 10 | |
| KRT18 | Keratin, type I cytoskeletal 18 | 10 | |
| KRT7 | Keratin, type I cytoskeletal 7 | 10 | |
| MTDH | Protein LYRIC | 10 | |
| RPS3 | 40S ribosomal protein S3 | 10 | |
| VDAC3 | Voltage-dependent anion-selective channel protein 3 | 10 | |
| YBX1 | Y-box-binding protein 1 | 10 | |
| ALDOA | Fructose-bisphosphate aldolase A | 9 | |
| CAV1 | Caveolin-1 | 9 | |
| CCT3 | T-complex protein 1 subunit gamma | 9 | |
| COPG1 | Coatomer subunit gamma-1 | 9 | yes |
| FLOT1 | Flotillin-1 | 9 | |
| GNB1 | Guanine nucleotide-binding protein G(I)/G(S)/G(T) subunit beta-1 | 9 | |
| MAP4 | Microtubule-associated protein 4 | 9 | |
| NNT | NAD(P) transhydrogenase, mitochondrial | 9 | |
| RPS4X | 40S ribosomal protein S4x | 9 | |
| TBL2 | Transducin beta-like protein 2 | 9 | |
| TRIM21 | Tripartite Motif Containing 21 | 9 | |
| WDR77 | Methylosome protein 50 | 9 | |
| ATAD3A | Isoform 2 of ATPase family AAA domain-containing protein 3A | 8 | |

| | | | |
|---|---|---|---|
| BCLAF1 | Bcl-2-associated transcription factor 1 | 8 | |
| EIF4B | Eukaryotic translation initiation factor 4B | 8 | |
| ENO1 | Alpha-enolase | 8 | |
| IFI16 | Gamma-interferon-inducible protein 16 | 8 | |
| LRRC8A | Volume-regulated anion channel subunit LRRC8A | 8 | |
| OCIAD2 | OCIA domain-containing protein 2 | 8 | |
| STT3B | Dolichyl-diphosphooligosaccharide--protein glycosyltransferase subunit STT3B | 8 | |
| TXNDC5 | Thioredoxin domain-containing protein 5 | 8 | |
| UQCRC2 | Cytochrome b-c1 complex subunit 2, mitochondrial | 8 | |
| AP2B1 | AP-2 complex subunit beta | 7 | |
| ATP6V0A1 | V-type proton ATPase 116 kDa subunit a 1 | 7 | |
| DLD | Dihydrolipoyl dehydrogenase, mitochondrial | 7 | |
| IGKV2D-24 | Probable non-functional immunoglobulin kappa variable 2D-24 | 7 | |
| IQGAP1 | Ras GTPase-activating-like protein IQGAP1 | 7 | |
| KRT2 | Keratin, type I cytoskeletal 2 | 7 | |
| KTN1 | Kinectin | 7 | |
| RPL5 | 60S ribosomal protein L5 | 7 | |
| RPLP0 | 60S acidic ribosomal protein P0 | 7 | |
| SEC62 | Translocation protein SEC62 | 7 | |
| CCT2 | T-complex protein 1 subunit beta | 6 | |
| IMMT | Inner Membrane Mitochondrial Protein | 6 | |
| KPNB1 | Importin subunit beta-1 | 6 | |
| MTCH2 | Mitochondrial carrier homolog 2 | 6 | |
| RPL11 | 60S ribosomal protein L11 | 6 | |
| RPL12 | 60S ribosomal protein L12 | 6 | |
| RPL14 | 60S ribosomal protein L14 | 6 | |
| SND1 | Staphylococcal nuclease domain-containing protein 1 | 6 | |
| SUN2 | SUN Domain-Containing Protein 2 | 6 | |
| TAB1 | TGF-beta-activated kinase 1 and MAP3K7-binding protein 1 | 6 | |
| TMTC3 | Protein O-mannosyl-transferase TMTC3 | 6 | |
| YWHAZ | 14-3-3 protein zeta/delta | 6 | yes |
| ABLIM1 | Actin Binding LIM Protein 1 | 5 | |
| ANKFY1 | Ankyrin Repeat And FYVE Domain Containing 1 | 5 | |
| CAPZA1 | F-actin-capping protein subunit alpha-1 | 5 | |
| CCT5 | T-complex protein 1 subunit epsilon | 5 | |
| COPB2 | Coatomer subunit beta' | 5 | yes |
| DLST | Dihydrolipoyllysine-residue succinyltransferase component of 2-oxoglutarate dehydrogenase complex, mitochondrial | 5 | |
| EHD2 | EH Domain-Containing Protein 2 | 5 | |
| EHD4 | EH domain-containing protein 4 | 5 | |
| EIF4A3 | Eukaryotic initiation factor 4A-III | 5 | |
| EMC1 | ER membrane protein complex subunit 1 | 5 | |
| ENDOD1 | Endonuclease domain-containing 1 protein | 5 | |
| GLG1 | Isoform 3 of Golgi apparatus protein 1 | 5 | |
| GLIPR2 | Golgi-associated plant pathogenesis-related protein 1 | 5 | |
| GNB2 | Guanine nucleotide-binding protein G(I)/G(S)/G(T) subunit beta-2 | 5 | |
| HSPA9 | Stress-70 protein, mitochondrial | 5 | |
| MGST3 | Microsomal glutathione S-transferase 3 | 5 | |
| MYH10 | Myosin-10 | 5 | |
| NES | Nestin | 5 | |
| PCBP1 | Poly(rC)-binding protein 1 | 5 | |
| SAMM50 | Sorting and assembly machinery component 50 homolog | 5 | |
| SERBP1 | Plasminogen activator inhibitor 1 RNA-binding protein | 5 | |
| STOM | Stomatin | 5 | |
| TPI1 | Triosephosphate isomerase | 5 | |
| ACAP2 | ArfGAP With Coiled-Coil, Ankyrin Repeat And PH Domains 2 | 4 | |
| ACTR1A | Alpha-centractin | 4 | |
| ADRM1 | Proteasomal ubiquitin receptor | 4 | |
| ANXA6 | Annexin A6 | 4 | |
| ATP6V1B2 | V-type proton ATPase subunit B, brain isoform | 4 | |
| CALR | Calreticulin | 4 | yes |
| CPT1A | Carnitine O-palmitoyltransferase 1, liver isoform | 4 | |
| DYNC1H1 | Cytoplasmic dynein 1 heavy chain 1 | 4 | |
| ESYT1 | Extended synaptotagmin-1 | 4 | |
| HCLS1 | Hematopoietic lineage cell-specific protein | 4 | |
| IGF2BP1 | Insulin-like growth factor 2 mRNA-binding protein 1 | 4 | |
| ILF2 | Interleukin enhancer-binding factor 2 | 4 | |
| LDHB | L-lactate dehydrogenase B chain | 4 | |
| LRRC8C | Volume-regulated anion channel subunit LRRC8C | 4 | |
| MAGT1 | Magnesium transporter protein 1 | 4 | |
| NOP2 | Probable 28S rRNA (cytosine(4447)-C(5))-methyltransferase | 4 | |

| | | | |
|---|---|---|---|
| RPL17 | 60S ribosomal protein L17 | 4 | |
| RPS19 | 40S ribosomal protein S19 | 4 | |
| RPS7 | 40S ribosomal protein S7 | 4 | |
| SURF4 | Surfeit locus protein 4 | 4 | |
| TMED10 | Transmembrane emp24 domain-containing protein 10 | 4 | |
| VAT1 | Synaptic vesicle membrane protein VAT-1 homolog | 4 | |
| WARS1 | Tryptophan--tRNA ligase, cytoplasmic | 4 | |
| YES1 | Tyrosine-protein kinase Yes | 4 | |
| ABCD3 | ATP-binding cassette sub-family D member 3 | 3 | |
| ATP13A1 | Endoplasmic reticulum transmembrane helix translocase | 3 | |
| ATP2A2 | Sarcoplasmic/endoplasmic reticulum calcium ATPase 2 | 3 | |
| CAP1 | Adenylyl cyclase-associated protein 1 | 3 | |
| CAPZB | F-actin-capping protein subunit beta | 3 | |
| CCT8 | T-Complex Protein 1 Subunit Theta | 3 | |
| CD44 | CD44 antigen | 3 | yes |
| DDX23 | Probable ATP-dependent RNA helicase DDX23 | 3 | |
| DHCR7 | 7-dehydrocholesterol reductase | 3 | |
| DHX15 | ATP-dependent RNA helicase DHX15 | 3 | |
| DNAJC11 | DnaJ homolog subfamily C member 11 | 3 | |
| EIF3A | Eukaryotic translation initiation factor 3 subunit A | 3 | |
| EIF3B | Isoform 2 of Eukaryotic translation initiation factor 3 subunit B | 3 | |
| EIF3I | Eukaryotic translation initiation factor 3 subunit I | 3 | |
| EMC2 | ER membrane protein complex subunit 2 | 3 | |
| FLII | Protein flightless-1 homolog | 3 | |
| FSCN1 | Fascin | 3 | |
| GOLGA3 | Golgin A3 | 3 | |
| H2AC6 | Histone H2A type 1-C | 3 | |
| HDLBP | Vigilin | 3 | |
| HP1BP3 | Heterochromatin protein 1-binding protein 3 | 3 | |
| ITPR3 | Inositol 1,4,5-Trisphosphate Receptor Type 3 | 3 | |
| KIF5B | Kinesin-1 heavy chain | 3 | |
| KRT16 | Keratin, type I cytoskeletal 16 | 3 | |
| MYADM | Myeloid-associated differentiation marker | 3 | |
| MYH14 | Myosin-14 | 3 | |
| NAPA | Alpha-soluble NSF attachment protein | 3 | |
| NOP56 | Nucleolar protein 56 | 3 | |
| NOP58 | Nucleolar protein 58 | 3 | |
| OSBPL3 | Oxysterol-binding protein-related protein 3 | 3 | yes |
| PA2G4 | Proliferation-associated protein 2G4 | 3 | |
| PARP1 | Poly [ADP-ribose] polymerase 1 | 3 | |
| PHOX2B | Paired mesoderm homeobox protein 2B | 3 | |
| PPM1B | Protein phosphatase 1B | 3 | |
| PPP1R12A | Protein phosphatase 1 regulatory subunit 12A | 3 | |
| PRPF31 | Pre-MRNA Processing Factor 31 | 3 | |
| PRPSAP1 | Phosphoribosyl pyrophosphate synthase-associated protein 1 | 3 | |
| RAB5C | Ras-related protein Rab-5C | 3 | |
| RPL23 | 60S ribosomal protein L23 | 3 | |
| RPS16 | 40S ribosomal protein S16 | 3 | |
| RPS26 | 40S ribosomal protein S26 | 3 | |
| SERPINH1 | Serpin H1 | 3 | |
| SLC25A11 | Mitochondrial 2-oxoglutarate/malate carrier protein | 3 | |
| SLC25A24 | Calcium-binding mitochondrial carrier protein SCaMC-1 | 3 | |
| SNRNP200 | Small Nuclear Ribonucleoprotein U5 Subunit 200 | 3 | |
| SRP14 | Signal recognition particle 14 kDa protein | 3 | |
| STING1 | Stimulator of interferon genes protein | 3 | |
| STT3A | Dolichyl-diphosphooligosaccharide--protein glycosyltransferase subunit STT3A | 3 | |
| TALDO1 | Transaldolase | 3 | |
| TCIRG1 | V-type proton ATPase 116 kDa subunit a 3 | 3 | |
| TCP1 | T-Complex 1 | 3 | |
| TICAM2 | Isoform 2 of TIR domain-containing adapter molecule 2 | 3 | |
| TMEM43 | Transmembrane Protein 43 | 3 | |
| TMPO | Lamina-associated polypeptide 2, isoforms beta/gamma | 3 | |
| XRCC5 | X-ray repair cross-complementing protein 5 | 3 | |
| ABCD1 | ATP-binding cassette sub-family D member 1 | 2 | |
| ACIN1 | Apoptotic chromatin condensation inducer in the nucleus | 2 | |
| ALB | Albumin | 2 | |
| ALYREF | THO complex subunit 4 | 2 | |
| AP2A2 | AP-2 complex subunit alpha-2 | 2 | |
| ARCN1 | Coatomer subunit delta | 2 | |
| ATP6V0A2 | ATPase H+ Transporting V0 Subunit A2 | 2 | |
| C3 | Complement C3 | 2 | |
| CALD1 | Caldesmon | 2 | |

| | | | |
|---|---|---|---|
| CCT4 | T-complex protein 1 subunit delta | 2 | |
| CD55 | Complement decay-accelerating factor | 2 | |
| CDC42 | Cell division control protein 42 homolog | 2 | |
| COL5A2 | N-acylneuraminate cytidylyltransferase | 2 | |
| COPS3 | COP9 signalosome complex subunit 3 | 2 | |
| DCTN1 | Dynactin Subunit 1 | 2 | |
| EEF1D | Isoform 2 of Elongation factor 1-delta | 2 | |
| EFHD2 | EF-hand domain-containing protein D2 | 2 | |
| EIF2S1 | Eukaryotic translation initiation factor 2 subunit 1 | 2 | |
| EIF3M | Eukaryotic translation initiation factor 3 subunit M | 2 | |
| ERLIN2 | Erlin-2 | 2 | |
| ERMP1 | Endoplasmic reticulum metallopeptidase 1 | 2 | |
| FXR1 | FMR1 Autosomal Protein-Like Protein 1 | 2 | |
| GDI2 | Rab GDP dissociation inhibitor beta | 2 | |
| H1-0 | Histone H1.0 | 2 | |
| ITGA2 | Integrin alpha-2 | 2 | yes |
| ITGA6 | Integrin alpha-6 | 2 | yes |
| ITGB1 | Integrin beta-1 | 2 | yes |
| KCTD12 | BTB/POZ domain-containing protein KCTD12 | 2 | |
| KCTD5 | BTB/POZ domain-containing protein KCTD5 | 2 | |
| KIF11 | Kinesin Family Member 11 | 2 | |
| KIF5C | Kinesin heavy chain isoform 5C | 2 | |
| LBR | Delta(14)-sterol reductase LBR | 2 | |
| LIMA1 | LIM domain and actin-binding protein 1 | 2 | |
| LMAN1 | Protein ERGIC-53 | 2 | |
| LMNB1 | Lamin-B1 | 2 | |
| LMNB2 | Lamin-B2 | 2 | |
| LRPPRC | Leucine-rich PPR motif-containing protein, mitochondrial | 2 | |
| MLEC | Malectin | 2 | |
| MYOF | Myoferlin | 2 | yes |
| NAP1L1 | Nucleosome assembly protein 1-like 1 | 2 | |
| PCBP2 | Poly(rC)-binding protein 2 | 2 | |
| PFKP | ATP-dependent 6-phosphofructokinase, platelet type | 2 | |
| PHGDH | D-3-phosphoglycerate dehydrogenase | 2 | |
| PHLDB1 | Pleckstrin homology-like domain family B member 1 | 2 | |
| PICALM | Phosphatidylinositol-binding clathrin assembly protein | 2 | |
| PKM | Pyruvate kinase PKM | 2 | |
| PLOD2 | Procollagen-lysine,2-oxoglutarate 5-dioxygenase 2 | 2 | |
| PLOD3 | Multifunctional procollagen lysine hydroxylase and glycosyltransferase LH3 | 2 | |
| PPP1CB | Serine/threonine-protein phosphatase PP1-beta catalytic subunit | 2 | |
| PPP1R18 | Phostensin | 2 | |
| PRPF8 | Pre-MRNA Processing Factor 8 | 2 | |
| PRPS1 | Ribose-phosphate pyrophosphokinase 1 | 2 | |
| PTDSS1 | Phosphatidylserine synthase 1 | 2 | |
| RAB11B | Ras-related protein Rab-11B | 2 | |
| RAB14 | Ras-related protein Rab-14 | 2 | |
| RAB7A | Ras-related protein Rab-7a | 2 | |
| RAC2 | Ras-related C3 botulinum toxin substrate 2 | 2 | |
| RER1 | Protein RER1 | 2 | |
| RHOC | Rho-related GTP-binding protein RhoC | 2 | |
| RNH1 | Ribonuclease inhibitor | 2 | |
| ROCK1 | Rho Associated Coiled-Coil Containing Protein Kinase 1 | 2 | |
| RPL21 | 60S ribosomal protein L21 | 2 | |
| RPL23A | 60S ribosomal protein L23a | 2 | |
| RPL29 | 60S ribosomal protein L29 | 2 | |
| RPL34 | 60S ribosomal protein L34 | 2 | |
| RPL35 | 60S ribosomal protein L35 | 2 | |
| RPL36A | 60S ribosomal protein L36a | 2 | |
| RPL9 | 60S ribosomal protein L9 | 2 | |
| RPS14 | 40S ribosomal protein S14 | 2 | |
| RPS2 | 40S ribosomal protein S2 | 2 | |
| RPS25 | 40S ribosomal protein S25 | 2 | |
| SART1 | U4/U6.U5 tri-snRNP-associated protein 1 | 2 | |
| SEC11A | Signal peptidase complex catalytic subunit SEC11A | 2 | |
| SERPINE1 | Serpin E1 | 2 | |
| SF3B1 | Splicing factor 3B subunit 1 | 2 | |
| SFXN1 | Sideroflexin-1 | 2 | |
| SLC12A4 | Solute carrier family 12 member 4 | 2 | |
| SRRT | Serrate RNA effector molecule homolog | 2 | |
| SRSF1 | Serine/arginine-rich splicing factor 1 | 2 | |
| SRSF3 | Serine/arginine-rich splicing factor 3 | 2 | |
| SRSF7 | Serine/arginine-rich splicing factor 7 | 2 | |

| | | | |
|---|---|---|---|
| SSRP1 | FACT complex subunit SSRP1 | 2 | |
| STK38 | Serine/Threonine Kinase 38 | 2 | |
| SVIL | Supervillin | 2 | |
| TAF4 | Transcription initiation factor TFIID subunit 4 | 2 | |
| THBS1 | Thrombospondin-1 | 2 | |
| TPM4 | Tropomyosin alpha-4 chain | 2 | |
| TPP1 | Tripeptidyl-peptidase 1 | 2 | |
| TRIM28 | Transcription intermediary factor 1-beta | 2 | |
| TUBA1C | Tubulin alpha-1C chain | 2 | yes |
| UGGT1 | UDP-glucose:glycoprotein glucosyltransferase 1 | 2 | yes |
| UQCRC1 | Cytochrome B-C1 Complex Subunit 1, Mitochondrial | 2 | |
| YBX3 | Y-box-binding protein 3 | 2 | |
| YWHAQ | 14-3-3 protein theta | 2 | yes |
| ZC3HAV1 | Zinc finger CCCH-type antiviral protein 1 | 2 | |
| ZMPSTE24 | CAAX prenyl protease 1 homolog | 2 | |
| ACTR3 | Actin-related protein 3 | 1 | |
| AKAP2 | A-kinase anchor protein 2 | 1 | |
| ANO10 | Anoctamin-10 | 1 | |
| APMAP | Adipocyte plasma membrane-associated protein | 1 | yes |
| APOA1 | Apolipoprotein A-I | 1 | |
| ATP5F1C | ATP synthase subunit gamma, mitochondrial | 1 | |
| ATP6AP1 | ATPase H+ Transporting Accessory Protein 1 | 1 | |
| ATP6V0D1 | ATPase H+ Transporting V0 Subunit D1 | 1 | |
| ATP6V1E1 | V-type proton ATPase subunit E 1 | 1 | |
| BST1 | Bone Marrow Stromal Cell Antigen 1 | 1 | |
| CALU | Calumenin | 1 | |
| CANX | Calnexin | 1 | yes |
| CCAR1 | Cell Division Cycle And Apoptosis Regulator 1 | 1 | |
| CDC25C | M-phase inducer phosphatase 3 | 1 | |
| CDC42BPB | Serine/threonine-protein kinase MRCK beta | 1 | |
| CDC5L | Cell division cycle 5-like protein | 1 | |
| CHTOP | Chromatin target of PRMT1 protein | 1 | |
| CMAS | N-acylneuraminate cytidylyltransferase | 1 | |
| CTNND1 | Catenin delta-1 | 1 | |
| CTSB | Cathepsin B | 1 | |
| CSRP1 | Cysteine and glycine-rich protein 1 | 1 | |
| DNHD1 | Dynein heavy chain domain-containing protein 1 | 1 | |
| EFTUD2 | 116 kDa U5 small nuclear ribonucleoprotein component | 1 | |
| EIF3L | Eukaryotic Translation Initiation Factor 3 Subunit L | 1 | |
| FERMT3 | Fermitin family homolog 3 | 1 | |
| FMNL2 | Formin-like protein 2 | 1 | |
| GANAB | Neutral alpha-glucosidase AB | 1 | |
| GNG12 | Guanine nucleotide-binding protein G(I)/G(S)/G(O) subunit gamma-12 | 1 | |
| HSPA1L | Heat shock 70 kDa protein 1-like | 1 | |
| IARS1 | Isoleucine--tRNA ligase, cytoplasmic | 1 | |
| KRT6A | Keratin, type I cytoskeletal 6A | 1 | |
| MCAM | Melanoma Cell Adhesion Molecule | 1 | |
| MCM3 | DNA replication licensing factor MCM3 | 1 | |
| PIGR | Polymeric immunoglobulin receptor | 1 | |
| PIK3C2A | Polymeric immunoglobulin receptor | 1 | |
| PLAT | ATP-dependent 6-phosphofructokinase, platelet type | 1 | |
| PPP1CA | Serine/threonine-protein phosphatase PP1-alpha catalytic subunit | 1 | |
| QARS1 | Glutamine--tRNA ligase | 1 | |
| RAB10 | Ras-related protein Rab-10 | 1 | |
| RACK1 | Receptor of activated protein C kinase 1 | 1 | |
| RALA | Ras-related protein Ral-A | 1 | |
| RALY | RALY Heterogeneous Nuclear Ribonucleoprotein | 1 | |
| RAN | GTP-binding nuclear protein Ran | 1 | |
| RB1CC1 | RB1-inducible coiled-coil protein 1 | 1 | |
| RPL10 | 60S ribosomal protein L10 | 1 | |
| RPL10A | 60S ribosomal protein L10a | 1 | |
| RPL13A | 60S ribosomal protein L13a | 1 | |
| RPL26 | 60S ribosomal protein L26 | 1 | |
| RPL27A | 60S ribosomal protein L27a | 1 | |
| RPL31 | 60S ribosomal protein L31 | 1 | |
| RSL1D1 | Ribosomal L1 Domain Containing 1 | 1 | |
| S100A11 | Protein S100-A11 | 1 | |
| SEC22B | Vesicle-trafficking protein SEC22b | 1 | |
| SF3B3 | Splicing factor 3B subunit 3 | 1 | |
| SFXN3 | Sideroflexin-3 | 1 | |
| SLC2A1 | Solute carrier family 2, facilitated glucose transporter member 1 | 1 | |

| | | | |
|---|---|---|---|
| SMCHD1 | Structural maintenance of chromosomes flexible hinge domain-containing protein 1 | 1 | |
| SRSF6 | Serine/arginine-rich splicing factor 6 | 1 | |
| SUPT16H | FACT complex subunit SPT16 | 1 | |
| SYNE1 | Nesprin-1 | 1 | |
| TMED9 | Transmembrane emp24 domain-containing protein 9 | 1 | |
| TMOD3 | Tropomodulin-3 | 1 | |
| TRA2B | Transformer 2 Beta Homolog | 1 | |
| TUBB4A | Tubulin beta-4A chain | 1 | yes |
| UBA52 | Ubiquitin-60S ribosomal protein L40 | 1 | |
| VAPA | VAMP Associated Protein A | 1 | |
| VARS1 | Valine--tRNA ligase | 1 | |

| | |
|---|---|
| **S7: TG2-associated proteins detected by both approaches (N-BAP-biotin rhTG2 and 3F-TG2)** | |
| ACIN1 | Apoptotic chromatin condensation inducer in the nucleus |
| AKAP2 | A-kinase anchor protein 2 |
| AP2A1 | AP-2 complex subunit alpha-1 |
| AP2B1 | AP-2 complex subunit beta |
| ATAD3A | Isoform 2 of ATPase family AAA domain-containing protein 3A |
| ATP5F1C | ATP synthase subunit gamma, mitochondrial |
| ATP6V1B2 | V-type proton ATPase subunit B, brain isoform |
| CAPZA1 | F-actin-capping protein subunit alpha-1 |
| CAPZB | F-actin-capping protein subunit beta |
| CAV1 | Caveolin-1 |
| CCT3 | T-complex protein 1 subunit gamma |
| CD59 | CD59 glycoprotein |
| CDC42 | Cell division control protein 42 homolog |
| CDC42BPB | Serine/threonine-protein kinase MRCK beta |
| COL5A2 | N-acylneuraminate cytidylyltransferase |
| CTNND1 | Catenin delta-1 |
| CTSB | Cathepsin B |
| DDOST | Dolichyl-diphosphooligosaccharide--protein glycosyltransferase 48 kDa subunit |
| DLD | Dihydrolipoyl dehydrogenase, mitochondrial |
| DLST | Dihydrolipoyllysine-residue succinyltransferase component of 2-oxoglutarate dehydrogenase complex, mitochondrial |
| DYNC1H1 | Cytoplasmic dynein 1 heavy chain 1 |
| EIF3A | Eukaryotic translation initiation factor 3 subunit A |
| ERLIN1 | Erlin-1 |
| ESYT1 | Extended synaptotagmin-1 |
| FLII | Protein flightless-1 homolog |
| FMNL2 | Formin-like protein 2 |
| GANAB | Neutral alpha-glucosidase AB |
| GLG1 | Isoform 3 of Golgi apparatus protein 1 |
| GNAI2 | Guanine nucleotide-binding protein G(i) subunit alpha-2 |
| GNB1 | Guanine nucleotide-binding protein G(I)/G(S)/G(T) subunit beta-1 |
| GNB2 | Guanine nucleotide-binding protein G(I)/G(S)/G(T) subunit beta-2 |
| GNG12 | Guanine nucleotide-binding protein G(I)/G(S)/G(O) subunit gamma-12 |
| H1-0 | Histone H1.0 |
| HDLBP | Vigilin |
| IFI16 | Gamma-interferon-inducible protein 16 |
| KIF5B | Kinesin-1 heavy chain |
| LMAN1 | Protein ERGIC-53 |
| LRPPRC | Leucine-rich PPR motif-containing protein, mitochondrial |
| MAP1B | Microtubule-associated protein 1B |
| MBOAT7 | Lysophospholipid acyltransferase 7 |
| MCAM | Melanoma Cell Adhesion Molecule |
| MGST3 | Microsomal glutathione S-transferase 3 |
| MYADM | Myeloid-associated differentiation marker |
| NOP56 | Nucleolar protein 56 |
| NOP58 | Nucleolar protein 58 |
| NT5E | 5'-nucleotidase |
| OCIAD2 | OCIA domain-containing protein 2 |
| PARP1 | Poly [ADP-ribose] polymerase 1 |
| PCBP2 | Poly(rC)-binding protein 2 |
| PHB2 | Prohibitin-2 |
| PHGDH | D-3-phosphoglycerate dehydrogenase |
| PLOD2 | Procollagen-lysine,2-oxoglutarate 5-dioxygenase 2 |
| QARS1 | Glutamine--tRNA ligase |
| RAB10 | Ras-related protein Rab-10 |
| RAB11B | Ras-related protein Rab-11B |
| RAB7A | Ras-related protein Rab-7a |
| RAC2 | Ras-related C3 botulinum toxin substrate 2 |
| RACK1 | Receptor of activated protein C kinase 1 |
| RALA | Ras-related protein Ral-A |
| RAN | GTP-binding nuclear protein Ran |
| RHOC | Rho-related GTP-binding protein RhoC |
| RPL5 | 60S ribosomal protein L5 |
| RPS26 | 40S ribosomal protein S26 |
| SEC62 | Translocation protein SEC62 |
| SFXN1 | Sideroflexin-1 |
| SFXN3 | Sideroflexin-3 |
| SRSF1 | Serine/arginine-rich splicing factor 1 |
| SRSF3 | Serine/arginine-rich splicing factor 3 |
| SSRP1 | FACT complex subunit SSRP1 |
| STING1 | Stimulator of interferon genes protein |
| STOM | Stomatin |
| SUN2 | SUN Domain-Containing Protein 2 |

| | |
|---|---|
| SURF4 | Surfeit locus protein 4 |
| TCIRG1 | V-type proton ATPase 116 kDa subunit a 3 |
| TMED10 | Transmembrane emp24 domain-containing protein 10 |
| TMED9 | Transmembrane emp24 domain-containing protein 9 |
| TMPO | Lamina-associated polypeptide 2, isoforms beta/gamma |
| TRPV2 | Transient receptor potential cation channel subfamily V member 2 |
| TUBB4A | Tubulin beta-4A chain |
| UBA52 | Ubiquitin-60S ribosomal protein L40 |
| UQCRC1 | Cytochrome B-C1 Complex Subunit 1, Mitochondrial |
| UQCRC2 | Cytochrome b-c1 complex subunit 2, mitochondrial |
| VDAC3 | Voltage-dependent anion-selective channel protein 3 |
| XRCC5 | X-ray repair cross-complementing protein 5 |
| YES1 | Tyrosine-protein kinase Yes |
| YWHAZ | 14-3-3 protein zeta/delta |

**S8: Enriched GO Molecular Function based on STRING database using anti-Flag M2 agarose resin in case of untreated sample (5≤ detected peptide count)**

| #term ID | term description | false discovery rate | matching proteins in network |
|---|---|---|---|
| GO:0003723 | RNA binding | 2.86e-17 | TRIM21,RPL8,RPS3,CCT5,KPNB1,CCT3,RPN1,HSPA9,PDIA3,PCBP1,TBL2,RPL13,RBM10,HSP90AA1,MTDH,RPL3,RPS3A,THRAP3,SND1,CAVIN1,MAP4,MSN,MYH10,RPL12,IFI16,RPL5,SERBP1,YBX1,RPS4X,RRBP1,AHNAK,CKAP4,KRT18,ACTN1,KTN1,YWHAZ,RPL14,IMMT,EIF4B,RPSA,BCLAF1,RPL18,RPLP0,ENO1,ALDOA,RPL11,EIF4A3 |
| GO:0005198 | Structural molecule activity | 1.61e-10 | RPL8,RPS3,MAP1B,RPL13,KRT2,TLN1,H2BC4,COPG1,KRT7,COPB2,RPL3,RPS3A,MAP4,MSN,RPL12,RPL5,RPS4X,AHNAK,KRT18,ACTN1,RPL14,RPSA,TUBA1A,MYL6,RPL18,RPLP0,RPL11 |
| GO:0003735 | Structural constituent of ribosome | 6.84e-08 | RPL8,RPS3,RPL13,RPL3,RPS3A,RPL12,RPL5,RPS4X,RPL14,RPSA,RPL18,RPLP0,RPL11 |
| GO:0003676 | Nucleic acid binding | 1.34e-07 | EHD4,WDR77,TRIM21,RPL8,EHD2,ENDOD1,RPS3,CCT5,KPNB1,CCT3,RPN1,HSPA9,PDIA3,PCBP1,TBL2,RPL13,PRMT5,H2BC4,RBM10,EEF1G,HSP90AA1,MTDH,RPL3,RPS3A,THRAP3,SND1,CAVIN1,MAP4,MSN,MYH10,RPL12,IFI16,RPL5,SERBP1,YBX1,RPS4X,RRBP1,AHNAK,CKAP4,KRT18,ACTN1,KTN1,YWHAZ,RPL14,IMMT,EIF4B,RPSA,BCLAF1,RPL18,RPLP0,ENO1,ALDOA,RPL11,EIF4A3 |
| GO:0019843 | rRNA binding | 5.11e-07 | RPL8,RPS3,RPL3,CAVIN1,RPL12,RPL5,RPS4X,RPLP0,RPL11 |
| GO:0097159 | Organic cyclic compound binding | 5.21e-07 | DLD,EHD4,WDR77,TRIM21,NT5E,RPL8,EHD2,NNT,ENDOD1,RPS3,CCT5,KPNB1,CCT3,RPN1,HSPA9,CCT2,PDIA3,PCBP1,TBL2,RPL13,GNAI2,PRMT5,H2BC4,RBM10,EEF1G,HSP90AA1,MTDH,CAV1,RPL3,RPS3A,THRAP3,SND1,CAVIN1,MAP4,MSN,MYH10,RPL12,TGM2,IFI16,RPL5,SERBP1,YBX1,RPS4X,RRBP1,AHNAK,CKAP4,ATAD3A,KRT18,ACTN1,KTN1,YWHAZ,RPL14,IMMT,EIF4B,RPSA,ERLIN1,VDAC3,BCLAF1,TUBA1A,ATP1A1,RPL18,RPLP0,ENO1,ALDOA,RPL11,EIF4A3,JAK1 |
| GO:0045296 | Cadherin binding | 1.10e-06 | EHD4,CAPZA1,IQGAP1,PCBP1,TLN1,EEF1G,SND1,SERBP1,AHNAK,KRT18,KTN1,YWHAZ,RPL14,ENO1,ALDOA |
| GO:1901363 | Heterocyclic compound binding | 2.24e-06 | DLD,EHD4,WDR77,TRIM21,NT5E,RPL8,EHD2,NNT,ENDOD1,RPS3,CCT5,KPNB1,CCT3,RPN1,HSPA9,CCT2,PDIA3,PCBP1,TBL2,RPL13,GNAI2,PRMT5,H2BC4,RBM10,EEF1G,HSP90AA1,MTDH,RPL3,RPS3A,THRAP3,SND1,CAVIN1,MAP4,MSN,MYH10,RPL12,TGM2,IFI16,RPL5,SERBP1,YBX1,RPS4X,RRBP1,AHNAK,CKAP4,ATAD3A,KRT18,ACTN1,KTN1,YWHAZ,RPL14,IMMT,EIF4B,RPSA,VDAC3,BCLAF1,TUBA1A,ATP1A1,RPL18,RPLP0,ENO1,ALDOA,RPL11,EIF4A3,JAK1 |
| GO:0050839 | Cell adhesion molecule binding | 2.94e-06 | EHD4,CAPZA1,IQGAP1,PCBP1,TLN1,EEF1G,SND1,MSN,SERBP1,AHNAK,KRT18,ACTN1,KTN1,YWHAZ,RPL14,RPSA,ENO1,ALDOA |
| GO:0005515 | Protein binding | 0.00012 | GLG1,TAB1,EHD4,TPI1,TRIM21,LRRC8A,CAPZA1,EHD2,ATP6V0A1,IQGAP1,ABLIM1,RPS3,CCT5,STOM,KPNB1,CCT3,MAP1B,HSPA9,CCT2,PDIA3,GNB2,PCBP1,TBL2,KRT2,GNAI2,TLN1,PRMT5,H2BC4,RBM10,EEF1G,HSP90AA1,MTDH,CAV1,THRAP3,SND1,CAVIN1,AP2A1,MAP4,MSN,MYH10,MGST3,IFI16,NES,RPL5,SERBP1,YBX1,FLOT1,GLIPR2,AHNAK,GNB1,ATAD3A,KRT18,ACTN1,KTN1,YWHAZ,RPL14,SUN2,RPSA,ERLIN1,TUBA1A,PHB2,ATP1A1,ANKFY1,AP2B1,ENO1,ALDOA,RPL11,CD59,JAK1 |
| GO:0005488 | Binding | 0.00077 | GLG1,DLD,TAB1,EHD4,TPI1,WDR77,RPN2,TRIM21,NT5E,LRRC8A,RPL8,CAPZA1,EHD2,ATP6V0A1,NNT,IQGAP1,UQCRC2,ABLIM1,ENDOD1,RPS3,CCT5,STOM,KPNB1,CCT3,STT3B,RPN1,ANXA5,MAP1B,HSPA9,CCT2,ANPEP,PDIA3,GNB2,PCBP1,TBL2,RPL13,KRT2,GNAI2,TLN1,PRMT5,H2BC4,RBM10,EEF1G,HSP90AA1,MTDH,CAV1,RPL3,RPS3A,THRAP3,SND1,CAVIN1,AP2A1,MAP4,MSN,MYH10,RPL12,TGM2,MGST3,IFI16,NES,RPL5,SERBP1,YBX1,RPS4X,FLOT1,RRBP1,GLIPR2,AHNAK,CKAP4,GNB1,ATAD3A,KRT18,ACTN1,KTN1,YWHAZ,RPL14,SUN2,IMMT,EIF4B,RPSA,ERLIN1,VDAC3,BCLAF1,TUBA1A,PHB2,ATP1A1,MYL6,RPL18,RPLP0,ANKFY1,AP2B1,ENO1,ALDOA,RPL11,EIF4A3,CD59,JAK1 |
| GO:0044877 | Protein-containing complex binding | 0.0014 | TAB1,RPN2,CAPZA1,IQGAP1,ABLIM1,RPS3,MAP1B,GNB2,GNAI2,TLN1,PRMT5,HSP90AA1,CAV1,SND1,MYH10,NES,SERBP1,GNB1,ACTN1,EIF4B,RPSA,EIF4A3 |
| GO:0003729 | mRNA binding | 0.0015 | RPS3,CCT5,PCBP1,HSP90AA1,RPS3A,MYH10,RPL5,SERBP1,YBX1,BCLAF1,EIF4A3 |
| GO:0044183 | Protein folding chaperone | 0.0073 | CCT5,CCT3,HSPA9,CCT2,HSP90AA1 |
| GO:0048027 | mRNA 5-UTR binding | 0.0083 | CCT5,RPS3A,MYH10,RPL5 |
| GO:0043021 | Ribonucleoprotein complex binding | 0.0116 | RPN2,PRMT5,SND1,SERBP1,EIF4B,RPSA,EIF4A3 |
| GO:0140662 | ATP-dependent protein folding chaperone | 0.0138 | CCT5,CCT3,HSPA9,CCT2 |
| GO:0008097 | 5S rRNA binding | 0.0139 | RPL3,RPL5,RPL11 |
| GO:0031625 | Ubiquitin protein ligase binding | 0.0190 | TPI1,HSPA9,CCT2,HSP90AA1,RPL5,YWHAZ,ERLIN1,RPL11,JAK1 |
| GO:0051020 | GTPase binding | 0.0204 | IQGAP1,KPNB1,GNB2,HSP90AA1,CAV1,YBX1,GNB1,ANKFY1,ENO1 |
| GO:0019899 | Enzyme binding | 0.0298 | TAB1,TPI1,TRIM21,ATP6V0A1,IQGAP1,RPS3,STOM,KPNB1,HSPA9,CCT2,GNB2,TBL2,HSP90AA1,CAV1,AP2A1,MSN,RPL5,YBX1,FLOT1,GNB1,YWHAZ,ERLIN1,ANKFY1,ENO1,RPL11,JAK1 |

| S9: Identified RNA-binding proteins in the immortalised HUVEC cells using OOPS and LC-MS/MS analysis | |
|---|---|
| **Identified Proteins (413)** | **Alternate ID** |
| Myosin-9 OS=Homo sapiens OX=9606 GN=MYH9 PE=1 SV=4 | MYH9 |
| Plectin OS=Homo sapiens OX=9606 GN=PLEC PE=1 SV=3 | PLEC |
| Vimentin OS=Homo sapiens OX=9606 GN=VIM PE=1 SV=4 | VIM |
| Filamin-A OS=Homo sapiens OX=9606 GN=FLNA PE=1 SV=4 | FLNA |
| Filamin-B OS=Homo sapiens OX=9606 GN=FLNB PE=1 SV=2 | FLNB |
| Annexin A2 OS=Homo sapiens OX=9606 GN=ANXA2 PE=1 SV=2 | ANXA2 |
| Neuroblast differentiation-associated protein AHNAK OS=Homo sapiens OX=9606 GN=AHNAK PE=1 SV=2 | AHNAK |
| Prelamin-A/C OS=Homo sapiens OX=9606 GN=LMNA PE=1 SV=1 | LMNA |
| Talin-1 OS=Homo sapiens OX=9606 GN=TLN1 PE=1 SV=3 | TLN1 |
| Actin, cytoplasmic 2 OS=Homo sapiens OX=9606 GN=ACTG1 PE=1 SV=1 | ACTG1 |
| Alpha-actinin-1 OS=Homo sapiens OX=9606 GN=ACTN1 PE=1 SV=2 | ACTN1 |
| Tubulin beta chain OS=Homo sapiens OX=9606 GN=TUBB PE=1 SV=2 | TUBB |
| Alpha-enolase OS=Homo sapiens OX=9606 GN=ENO1 PE=1 SV=2 | ENO1 |
| Cytoplasmic dynein 1 heavy chain 1 OS=Homo sapiens OX=9606 GN=DYNC1H1 PE=1 SV=5 | DYNC1H1 |
| Pyruvate kinase PKM OS=Homo sapiens OX=9606 GN=PKM PE=1 SV=4 | PKM |
| Endoplasmic reticulum chaperone BiP OS=Homo sapiens OX=9606 GN=HSPA5 PE=1 SV=2 | HSPA5 |
| Filamin-C OS=Homo sapiens OX=9606 GN=FLNC PE=1 SV=3 | FLNC |
| Elongation factor 2 OS=Homo sapiens OX=9606 GN=EEF2 PE=1 SV=4 | EEF2 |
| Clathrin heavy chain 1 OS=Homo sapiens OX=9606 GN=CLTC PE=1 SV=5 | CLTC |
| Isoform 2 of Tubulin alpha-1A chain OS=Homo sapiens OX=9606 GN=TUBA1A | TUBA1A |
| Heat shock protein HSP 90-beta OS=Homo sapiens OX=9606 GN=HSP90AB1 PE=1 SV=4 | HSP90AB1 |
| Heat shock cognate 71 kDa protein OS=Homo sapiens OX=9606 GN=HSPA8 PE=1 SV=1 | HSPA8 |
| Glyceraldehyde-3-phosphate dehydrogenase OS=Homo sapiens OX=9606 GN=GAPDH PE=1 SV=3 | GAPDH |
| Annexin A1 OS=Homo sapiens OX=9606 GN=ANXA1 PE=1 SV=2 | ANXA1 |
| **Protein-glutamine gamma-glutamyltransferase 2 OS=Homo sapiens OX=9606 GN=TGM2 PE=1 SV=2** | **TGM2** |
| Transketolase OS=Homo sapiens OX=9606 GN=TKT PE=1 SV=3 | TKT |
| Heterogeneous nuclear ribonucleoprotein U OS=Homo sapiens OX=9606 GN=HNRNPU PE=1 SV=6 | HNRNPU |
| Transitional endoplasmic reticulum ATPase OS=Homo sapiens OX=9606 GN=VCP PE=1 SV=4 | VCP |
| Isoform 2 of Ubiquitin-like modifier-activating enzyme 1 OS=Homo sapiens OX=9606 GN=UBA1 | UBA1 |
| Spectrin beta chain, non-erythrocytic 1 OS=Homo sapiens OX=9606 GN=SPTBN1 PE=1 SV=2 | SPTBN1 |
| Alpha-actinin-4 OS=Homo sapiens OX=9606 GN=ACTN4 PE=1 SV=2 | ACTN4 |
| Protein disulfide-isomerase A3 OS=Homo sapiens OX=9606 GN=PDIA3 PE=1 SV=4 | PDIA3 |
| Elongation factor 1-alpha 1 OS=Homo sapiens OX=9606 GN=EEF1A1 PE=1 SV=1 | EEF1A1 |
| Fibronectin OS=Homo sapiens OX=9606 GN=FN1 PE=1 SV=5 | FN1 |
| Heterogeneous nuclear ribonucleoproteins A2/B1 OS=Homo sapiens OX=9606 GN=HNRNPA2B1 PE=1 SV=2 | HNRNPA2B1 |
| Myoferlin OS=Homo sapiens OX=9606 GN=MYOF PE=1 SV=1 | MYOF |
| Polyubiquitin-B OS=Homo sapiens OX=9606 GN=UBB PE=1 SV=1 | UBB |
| Basement membrane-specific heparan sulfate proteoglycan core protein OS=Homo sapiens OX=9606 GN=HSPG2 PE=1 SV=4 | HSPG2 |
| Nucleolin OS=Homo sapiens OX=9606 GN=NCL PE=1 SV=3 | NCL |
| Fructose-bisphosphate aldolase A OS=Homo sapiens OX=9606 GN=ALDOA PE=1 SV=2 | ALDOA |
| Moesin OS=Homo sapiens OX=9606 GN=MSN PE=1 SV=3 | MSN |
| Spectrin alpha chain, non-erythrocytic 1 OS=Homo sapiens OX=9606 GN=SPTAN1 PE=1 SV=3 | SPTAN1 |
| Cofilin-1 OS=Homo sapiens OX=9606 GN=CFL1 PE=1 SV=3 | CFL1 |
| Unconventional myosin-Ic OS=Homo sapiens OX=9606 GN=MYO1C PE=1 SV=4 | MYO1C |
| Vinculin OS=Homo sapiens OX=9606 GN=VCL PE=1 SV=4 | VCL |
| Peroxiredoxin-1 OS=Homo sapiens OX=9606 GN=PRDX1 PE=1 SV=1 | PRDX1 |
| Heterogeneous nuclear ribonucleoprotein R OS=Homo sapiens OX=9606 GN=HNRNPR PE=1 SV=1 | HNRNPR |
| Major vault protein OS=Homo sapiens OX=9606 GN=MVP PE=1 SV=4 | MVP |
| Annexin A6 OS=Homo sapiens OX=9606 GN=ANXA6 PE=1 SV=3 | ANXA6 |
| Staphylococcal nuclease domain-containing protein 1 OS=Homo sapiens OX=9606 GN=SND1 PE=1 SV=1 | SND1 |

| | |
|---|---|
| Thrombospondin-1 OS=Homo sapiens OX=9606 GN=THBS1 PE=1 SV=2 | THBS1 |
| Cytoskeleton-associated protein 4 OS=Homo sapiens OX=9606 GN=CKAP4 PE=1 SV=2 | CKAP4 |
| Heterogeneous nuclear ribonucleoprotein H OS=Homo sapiens OX=9606 GN=HNRNPH1 PE=1 SV=4 | HNRNPH1 |
| Microtubule-associated protein 4 OS=Homo sapiens OX=9606 GN=MAP4 PE=1 SV=3 | MAP4 |
| DNA-dependent protein kinase catalytic subunit OS=Homo sapiens OX=9606 GN=PRKDC PE=1 SV=3 | PRKDC |
| ATP-dependent 6-phosphofructokinase, platelet type OS=Homo sapiens OX=9606 GN=PFKP PE=1 SV=2 | PFKP |
| Thioredoxin domain-containing protein 5 OS=Homo sapiens OX=9606 GN=TXNDC5 PE=1 SV=2 | TXNDC5 |
| Probable ATP-dependent RNA helicase DDX17 OS=Homo sapiens OX=9606 GN=DDX17 PE=1 SV=2 | DDX17 |
| Chloride intracellular channel protein 1 OS=Homo sapiens OX=9606 GN=CLIC1 PE=1 SV=4 | CLIC1 |
| Stress-70 protein, mitochondrial OS=Homo sapiens OX=9606 GN=HSPA9 PE=1 SV=2 | HSPA9 |
| Annexin A5 OS=Homo sapiens OX=9606 GN=ANXA5 PE=1 SV=2 | ANXA5 |
| Lamin-B1 OS=Homo sapiens OX=9606 GN=LMNB1 PE=1 SV=2 | LMNB1 |
| Endoplasmin OS=Homo sapiens OX=9606 GN=HSP90B1 PE=1 SV=1 | HSP90B1 |
| WD repeat-containing protein 1 OS=Homo sapiens OX=9606 GN=WDR1 PE=1 SV=4 | WDR1 |
| Heterogeneous nuclear ribonucleoprotein K OS=Homo sapiens OX=9606 GN=HNRNPK PE=1 SV=1 | HNRNPK |
| Far upstream element-binding protein 2 OS=Homo sapiens OX=9606 GN=KHSRP PE=1 SV=4 | KHSRP |
| Peptidyl-prolyl cis-trans isomerase A OS=Homo sapiens OX=9606 GN=PPIA PE=1 SV=2 | PPIA |
| Protein disulfide-isomerase A6 OS=Homo sapiens OX=9606 GN=PDIA6 PE=1 SV=1 | PDIA6 |
| ATP-dependent RNA helicase A OS=Homo sapiens OX=9606 GN=DHX9 PE=1 SV=4 | DHX9 |
| ATP synthase subunit alpha, mitochondrial OS=Homo sapiens OX=9606 GN=ATP5F1A PE=1 SV=1 | ATP5F1A |
| 60S ribosomal protein L4 OS=Homo sapiens OX=9606 GN=RPL4 PE=1 SV=5 | RPL4 |
| T-complex protein 1 subunit beta OS=Homo sapiens OX=9606 GN=CCT2 PE=1 SV=4 | CCT2 |
| Src substrate cortactin OS=Homo sapiens OX=9606 GN=CTTN PE=1 SV=2 | CTTN |
| X-ray repair cross-complementing protein 6 OS=Homo sapiens OX=9606 GN=XRCC6 PE=1 SV=2 | XRCC6 |
| T-complex protein 1 subunit delta OS=Homo sapiens OX=9606 GN=CCT4 PE=1 SV=4 | CCT4 |
| Kinesin-1 heavy chain OS=Homo sapiens OX=9606 GN=KIF5B PE=1 SV=1 | KIF5B |
| Ras GTPase-activating-like protein IQGAP1 OS=Homo sapiens OX=9606 GN=IQGAP1 PE=1 SV=1 | IQGAP1 |
| Guanine nucleotide-binding protein G(i) subunit alpha-2 OS=Homo sapiens OX=9606 GN=GNAI2 PE=1 SV=3 | GNAI2 |
| BTB/POZ domain-containing protein KCTD12 OS=Homo sapiens OX=9606 GN=KCTD12 PE=1 SV=1 | KCTD12 |
| Vigilin OS=Homo sapiens OX=9606 GN=HDLBP PE=1 SV=2 | HDLBP |
| Calpain-2 catalytic subunit OS=Homo sapiens OX=9606 GN=CAPN2 PE=1 SV=6 | CAPN2 |
| Tubulin beta-6 chain OS=Homo sapiens OX=9606 GN=TUBB6 PE=1 SV=1 | TUBB6 |
| Polypyrimidine tract-binding protein 1 OS=Homo sapiens OX=9606 GN=PTBP1 PE=1 SV=1 | PTBP1 |
| Glucose-6-phosphate 1-dehydrogenase OS=Homo sapiens OX=9606 GN=G6PD PE=1 SV=4 | G6PD |
| L-lactate dehydrogenase A chain OS=Homo sapiens OX=9606 GN=LDHA PE=1 SV=2 | LDHA |
| Coronin-1B OS=Homo sapiens OX=9606 GN=CORO1B PE=1 SV=1 | CORO1B |
| Neutral alpha-glucosidase AB OS=Homo sapiens OX=9606 GN=GANAB PE=1 SV=3 | GANAB |
| Glutathione S-transferase P OS=Homo sapiens OX=9606 GN=GSTP1 PE=1 SV=2 | GSTP1 |
| T-complex protein 1 subunit theta OS=Homo sapiens OX=9606 GN=CCT8 PE=1 SV=4 | CCT8 |
| 40S ribosomal protein SA OS=Homo sapiens OX=9606 GN=RPSA PE=1 SV=4 | RPSA |
| Bifunctional glutamate/proline--tRNA ligase OS=Homo sapiens OX=9606 GN=EPRS1 PE=1 SV=5 | EPRS1 |
| Dolichyl-diphosphooligosaccharide--protein glycosyltransferase subunit 1 OS=Homo sapiens OX=9606 GN=RPN1 PE=1 SV=1 | RPN1 |
| ELAV-like protein 1 OS=Homo sapiens OX=9606 GN=ELAVL1 PE=1 SV=2 | ELAVL1 |

| | |
|---|---|
| Keratin, type II cytoskeletal 1 OS=Homo sapiens OX=9606 GN=KRT1 PE=1 SV=6 | KRT1 |
| Cell surface glycoprotein MUC18 OS=Homo sapiens OX=9606 GN=MCAM PE=1 SV=2 | MCAM |
| Phosphoglycerate kinase 1 OS=Homo sapiens OX=9606 GN=PGK1 PE=1 SV=3 | PGK1 |
| Fatty acid synthase OS=Homo sapiens OX=9606 GN=FASN PE=1 SV=3 | FASN |
| Nestin OS=Homo sapiens OX=9606 GN=NES PE=1 SV=2 | NES |
| Myosin light polypeptide 6 OS=Homo sapiens OX=9606 GN=MYL6 PE=1 SV=2 | MYL6 |
| Synaptic vesicle membrane protein VAT-1 homolog OS=Homo sapiens OX=9606 GN=VAT1 PE=1 SV=2 | VAT1 |
| Keratin, type II cytoskeletal 7 OS=Homo sapiens OX=9606 GN=KRT7 PE=1 SV=5 | KRT7 |
| Heterogeneous nuclear ribonucleoprotein F OS=Homo sapiens OX=9606 GN=HNRNPF PE=1 SV=3 | HNRNPF |
| Multifunctional protein ADE2 OS=Homo sapiens OX=9606 GN=PAICS PE=1 SV=3 | PAICS |
| Ras-interacting protein 1 OS=Homo sapiens OX=9606 GN=RASIP1 PE=1 SV=1 | RASIP1 |
| D-3-phosphoglycerate dehydrogenase OS=Homo sapiens OX=9606 GN=PHGDH PE=1 SV=4 | PHGDH |
| Isoform 2 of Protein flightless-1 homolog OS=Homo sapiens OX=9606 GN=FLII | FLII |
| Ribosome-binding protein 1 OS=Homo sapiens OX=9606 GN=RRBP1 PE=1 SV=5 | RRBP1 |
| 60 kDa heat shock protein, mitochondrial OS=Homo sapiens OX=9606 GN=HSPD1 PE=1 SV=2 | HSPD1 |
| Protein disulfide-isomerase OS=Homo sapiens OX=9606 GN=P4HB PE=1 SV=3 | P4HB |
| EH domain-containing protein 2 OS=Homo sapiens OX=9606 GN=EHD2 PE=1 SV=2 | EHD2 |
| Plastin-3 OS=Homo sapiens OX=9606 GN=PLS3 PE=1 SV=4 | PLS3 |
| Coatomer subunit alpha OS=Homo sapiens OX=9606 GN=COPA PE=1 SV=2 | COPA |
| Transcription intermediary factor 1-beta OS=Homo sapiens OX=9606 GN=TRIM28 PE=1 SV=5 | TRIM28 |
| Importin subunit beta-1 OS=Homo sapiens OX=9606 GN=KPNB1 PE=1 SV=2 | KPNB1 |
| Isoform 2 of Tropomyosin alpha-3 chain OS=Homo sapiens OX=9606 GN=TPM3 | TPM3 |
| Transaldolase OS=Homo sapiens OX=9606 GN=TALDO1 PE=1 SV=2 | TALDO1 |
| Matrin-3 OS=Homo sapiens OX=9606 GN=MATR3 PE=1 SV=2 | MATR3 |
| C-1-tetrahydrofolate synthase, cytoplasmic OS=Homo sapiens OX=9606 GN=MTHFD1 PE=1 SV=4 | MTHFD1 |
| Heat shock protein beta-1 OS=Homo sapiens OX=9606 GN=HSPB1 PE=1 SV=2 | HSPB1 |
| Microtubule-associated protein 1B OS=Homo sapiens OX=9606 GN=MAP1B PE=1 SV=2 | MAP1B |
| 60S ribosomal protein L6 OS=Homo sapiens OX=9606 GN=RPL6 PE=1 SV=3 | RPL6 |
| Isoform 3 of Zinc finger protein 185 OS=Homo sapiens OX=9606 GN=ZNF185 | ZNF185 |
| F-actin-capping protein subunit alpha-2 OS=Homo sapiens OX=9606 GN=CAPZA2 PE=1 SV=3 | CAPZA2 |
| Transgelin-2 OS=Homo sapiens OX=9606 GN=TAGLN2 PE=1 SV=3 | TAGLN2 |
| Heterogeneous nuclear ribonucleoprotein M OS=Homo sapiens OX=9606 GN=HNRNPM PE=1 SV=3 | HNRNPM |
| Heterogeneous nuclear ribonucleoproteins C1/C2 OS=Homo sapiens OX=9606 GN=HNRNPC PE=1 SV=4 | HNRNPC |
| Catenin delta-1 OS=Homo sapiens OX=9606 GN=CTNND1 PE=1 SV=1 | CTNND1 |
| Splicing factor, proline- and glutamine-rich OS=Homo sapiens OX=9606 GN=SFPQ PE=1 SV=2 | SFPQ |
| 60S ribosomal protein L8 OS=Homo sapiens OX=9606 GN=RPL8 PE=1 SV=2 | RPL8 |
| PDZ and LIM domain protein 1 OS=Homo sapiens OX=9606 GN=PDLIM1 PE=1 SV=4 | PDLIM1 |
| Coatomer subunit gamma-1 OS=Homo sapiens OX=9606 GN=COPG1 PE=1 SV=1 | COPG1 |
| Heat shock protein HSP 90-alpha OS=Homo sapiens OX=9606 GN=HSP90AA1 PE=1 SV=5 | HSP90AA1 |
| ATP-dependent RNA helicase DDX3X OS=Homo sapiens OX=9606 GN=DDX3X PE=1 SV=3 | DDX3X |
| Rab GDP dissociation inhibitor beta OS=Homo sapiens OX=9606 GN=GDI2 PE=1 SV=2 | GDI2 |
| CAD protein OS=Homo sapiens OX=9606 GN=CAD PE=1 SV=3 | CAD |
| Tubulin beta-4B chain OS=Homo sapiens OX=9606 GN=TUBB4B PE=1 SV=1 | TUBB4B |

| | |
|---|---|
| Protein AHNAK2 OS=Homo sapiens OX=9606 GN=AHNAK2 PE=1 SV=2 | AHNAK2 |
| Isoform 6 of Unconventional myosin-VI OS=Homo sapiens OX=9606 GN=MYO6 | MYO6 |
| Cytoplasmic FMR1-interacting protein 1 OS=Homo sapiens OX=9606 GN=CYFIP1 PE=1 SV=1 | CYFIP1 |
| Heterogeneous nuclear ribonucleoprotein A1 OS=Homo sapiens OX=9606 GN=HNRNPA1 PE=1 SV=5 | HNRNPA1 |
| Cysteine--tRNA ligase, cytoplasmic OS=Homo sapiens OX=9606 GN=CARS1 PE=1 SV=3 | CARS1 |
| Inositol 1,4,5-trisphosphate receptor type 3 OS=Homo sapiens OX=9606 GN=ITPR3 PE=1 SV=2 | ITPR3 |
| Sodium/potassium-transporting ATPase subunit alpha-1 OS=Homo sapiens OX=9606 GN=ATP1A1 PE=1 SV=1 | ATP1A1 |
| Nuclear mitotic apparatus protein 1 OS=Homo sapiens OX=9606 GN=NUMA1 PE=1 SV=2 | NUMA1 |
| EH domain-containing protein 4 OS=Homo sapiens OX=9606 GN=EHD4 PE=1 SV=1 | EHD4 |
| Eukaryotic translation initiation factor 3 subunit C-like protein OS=Homo sapiens OX=9606 GN=EIF3CL PE=1 SV=1 | EIF3CL |
| Valine--tRNA ligase OS=Homo sapiens OX=9606 GN=VARS1 PE=1 SV=4 | VARS1 |
| Isoform 2 of Glycine--tRNA ligase OS=Homo sapiens OX=9606 GN=GARS1 | GARS1 |
| T-complex protein 1 subunit gamma OS=Homo sapiens OX=9606 GN=CCT3 PE=1 SV=4 | CCT3 |
| Programmed cell death 6-interacting protein OS=Homo sapiens OX=9606 GN=PDCD6IP PE=1 SV=1 | PDCD6IP |
| ATP-citrate synthase OS=Homo sapiens OX=9606 GN=ACLY PE=1 SV=3 | ACLY |
| Integrin beta-1 OS=Homo sapiens OX=9606 GN=ITGB1 PE=1 SV=2 | ITGB1 |
| Coronin-1C OS=Homo sapiens OX=9606 GN=CORO1C PE=1 SV=1 | CORO1C |
| Eukaryotic initiation factor 4A-I OS=Homo sapiens OX=9606 GN=EIF4A1 PE=1 SV=1 | EIF4A1 |
| Phosphoglycerate mutase 1 OS=Homo sapiens OX=9606 GN=PGAM1 PE=1 SV=2 | PGAM1 |
| Galectin-1 OS=Homo sapiens OX=9606 GN=LGALS1 PE=1 SV=2 | LGALS1 |
| LIM and SH3 domain protein 1 OS=Homo sapiens OX=9606 GN=LASP1 PE=1 SV=2 | LASP1 |
| Fermitin family homolog 3 OS=Homo sapiens OX=9606 GN=FERMT3 PE=1 SV=1 | FERMT3 |
| Profilin-1 OS=Homo sapiens OX=9606 GN=PFN1 PE=1 SV=2 | PFN1 |
| EF-hand domain-containing protein D2 OS=Homo sapiens OX=9606 GN=EFHD2 PE=1 SV=1 | EFHD2 |
| T-complex protein 1 subunit zeta OS=Homo sapiens OX=9606 GN=CCT6A PE=1 SV=3 | CCT6A |
| Serine/threonine-protein kinase N1 OS=Homo sapiens OX=9606 GN=PKN1 PE=1 SV=2 | PKN1 |
| Ubiquitin carboxyl-terminal hydrolase 5 OS=Homo sapiens OX=9606 GN=USP5 PE=1 SV=2 | USP5 |
| 14-3-3 protein zeta/delta OS=Homo sapiens OX=9606 GN=YWHAZ PE=1 SV=1 | YWHAZ |
| 40S ribosomal protein S8 OS=Homo sapiens OX=9606 GN=RPS8 PE=1 SV=2 | RPS8 |
| SH3 and multiple ankyrin repeat domains protein 3 OS=Homo sapiens OX=9606 GN=SHANK3 PE=1 SV=3 | SHANK3 |
| EH domain-containing protein 1 OS=Homo sapiens OX=9606 GN=EHD1 PE=1 SV=2 | EHD1 |
| Arf-GAP with Rho-GAP domain, ANK repeat and PH domain-containing protein 1 OS=Homo sapiens OX=9606 GN=ARAP1 PE=1 SV=3 | ARAP1 |
| 60S ribosomal protein L5 OS=Homo sapiens OX=9606 GN=RPL5 PE=1 SV=3 | RPL5 |
| Zyxin OS=Homo sapiens OX=9606 GN=ZYX PE=1 SV=1 | ZYX |
| X-ray repair cross-complementing protein 5 OS=Homo sapiens OX=9606 GN=XRCC5 PE=1 SV=3 | XRCC5 |
| Dihydropyrimidinase-related protein 2 OS=Homo sapiens OX=9606 GN=DPYSL2 PE=1 SV=1 | DPYSL2 |
| Probable ATP-dependent RNA helicase DDX5 OS=Homo sapiens OX=9606 GN=DDX5 PE=1 SV=1 | DDX5 |
| V-type proton ATPase catalytic subunit A OS=Homo sapiens OX=9606 GN=ATP6V1A PE=1 SV=2 | ATP6V1A |
| T-complex protein 1 subunit alpha OS=Homo sapiens OX=9606 GN=TCP1 PE=1 SV=1 | TCP1 |
| Heterogeneous nuclear ribonucleoprotein L OS=Homo sapiens OX=9606 GN=HNRNPL PE=1 SV=2 | HNRNPL |
| Constitutive coactivator of PPAR-gamma-like protein 1 OS=Homo sapiens OX=9606 GN=FAM120A PE=1 SV=2 | FAM120A |
| Elongation factor 1-gamma OS=Homo sapiens OX=9606 GN=EEF1G PE=1 SV=3 | EEF1G |

| | |
|---|---|
| T-complex protein 1 subunit epsilon OS=Homo sapiens OX=9606 GN=CCT5 PE=1 SV=1 | CCT5 |
| Galactokinase OS=Homo sapiens OX=9606 GN=GALK1 PE=1 SV=1 | GALK1 |
| Copine-1 OS=Homo sapiens OX=9606 GN=CPNE1 PE=1 SV=1 | CPNE1 |
| Histidine--tRNA ligase, cytoplasmic OS=Homo sapiens OX=9606 GN=HARS1 PE=1 SV=2 | HARS1 |
| Actin, cytoplasmic 1 OS=Homo sapiens OX=9606 GN=ACTB PE=1 SV=1 | ACTB |
| Tryptophan--tRNA ligase, cytoplasmic OS=Homo sapiens OX=9606 GN=WARS1 PE=1 SV=2 | WARS1 |
| Non-POU domain-containing octamer-binding protein OS=Homo sapiens OX=9606 GN=NONO PE=1 SV=4 | NONO |
| L-lactate dehydrogenase B chain OS=Homo sapiens OX=9606 GN=LDHB PE=1 SV=2 | LDHB |
| Isoform 5 of Septin-9 OS=Homo sapiens OX=9606 GN=SEPTIN9 | SEPTIN9 |
| RNA-binding motif protein, X chromosome OS=Homo sapiens OX=9606 GN=RBMX PE=1 SV=3 | RBMX |
| Drebrin-like protein OS=Homo sapiens OX=9606 GN=DBNL PE=1 SV=1 | DBNL |
| 60S ribosomal protein L24 OS=Homo sapiens OX=9606 GN=RPL24 PE=1 SV=1 | RPL24 |
| Sequestosome-1 OS=Homo sapiens OX=9606 GN=SQSTM1 PE=1 SV=1 | SQSTM1 |
| Fascin OS=Homo sapiens OX=9606 GN=FSCN1 PE=1 SV=3 | FSCN1 |
| 60S ribosomal protein L13 OS=Homo sapiens OX=9606 GN=RPL13 PE=1 SV=4 | RPL13 |
| GTPase-activating protein and VPS9 domain-containing protein 1 OS=Homo sapiens OX=9606 GN=GAPVD1 PE=1 SV=2 | GAPVD1 |
| 14-3-3 protein theta OS=Homo sapiens OX=9606 GN=YWHAQ PE=1 SV=1 | YWHAQ |
| Leucine-rich repeat-containing protein 47 OS=Homo sapiens OX=9606 GN=LRRC47 PE=1 SV=1 | LRRC47 |
| Nucleoplasmin-3 OS=Homo sapiens OX=9606 GN=NPM3 PE=1 SV=3 | NPM3 |
| Aminopeptidase N OS=Homo sapiens OX=9606 GN=ANPEP PE=1 SV=4 | ANPEP |
| Ribonuclease inhibitor OS=Homo sapiens OX=9606 GN=RNH1 PE=1 SV=2 | RNH1 |
| Catenin alpha-1 OS=Homo sapiens OX=9606 GN=CTNNA1 PE=1 SV=1 | CTNNA1 |
| Collagen alpha-1(XVIII) chain OS=Homo sapiens OX=9606 GN=COL18A1 PE=1 SV=5 | COL18A1 |
| Microtubule-actin cross-linking factor 1, isoforms 1/2/3/5 OS=Homo sapiens OX=9606 GN=MACF1 PE=1 SV=4 | MACF1 |
| Isoform 4 of A-kinase anchor protein 2 OS=Homo sapiens OX=9606 GN=AKAP2 | AKAP2 |
| Leucine--tRNA ligase, cytoplasmic OS=Homo sapiens OX=9606 GN=LARS1 PE=1 SV=2 | LARS1 |
| Histone H3.1 OS=Homo sapiens OX=9606 GN=H3C1 PE=1 SV=2 | H3C1 |
| PDZ and LIM domain protein 7 OS=Homo sapiens OX=9606 GN=PDLIM7 PE=1 SV=1 | PDLIM7 |
| 40S ribosomal protein S3a OS=Homo sapiens OX=9606 GN=RPS3A PE=1 SV=2 | RPS3A |
| Protein disulfide-isomerase A4 OS=Homo sapiens OX=9606 GN=PDIA4 PE=1 SV=2 | PDIA4 |
| Heterogeneous nuclear ribonucleoprotein U-like protein 2 OS=Homo sapiens OX=9606 GN=HNRNPUL2 PE=1 SV=1 | HNRNPUL2 |
| Extended synaptotagmin-1 OS=Homo sapiens OX=9606 GN=ESYT1 PE=1 SV=1 | ESYT1 |
| Isoform Beta-4B of Integrin beta-4 OS=Homo sapiens OX=9606 GN=ITGB4 | ITGB4 |
| Cysteine-rich protein 2 OS=Homo sapiens OX=9606 GN=CRIP2 PE=1 SV=1 | CRIP2 |
| Insulin-like growth factor 2 mRNA-binding protein 3 OS=Homo sapiens OX=9606 GN=IGF2BP3 PE=1 SV=2 | IGF2BP3 |
| Triosephosphate isomerase OS=Homo sapiens OX=9606 GN=TPI1 PE=1 SV=4 | TPI1 |
| Drebrin OS=Homo sapiens OX=9606 GN=DBN1 PE=1 SV=4 | DBN1 |
| Isoform LCRMP-4 of Dihydropyrimidinase-related protein 3 OS=Homo sapiens OX=9606 GN=DPYSL3 | DPYSL3 |
| S-adenosylmethionine synthase isoform type-2 OS=Homo sapiens OX=9606 GN=MAT2A PE=1 SV=1 | MAT2A |
| Calponin-2 OS=Homo sapiens OX=9606 GN=CNN2 PE=1 SV=4 | CNN2 |
| Transportin-1 OS=Homo sapiens OX=9606 GN=TNPO1 PE=1 SV=2 | TNPO1 |
| Protein phosphatase 1F OS=Homo sapiens OX=9606 GN=PPM1F PE=1 SV=3 | PPM1F |
| ATP-dependent 6-phosphofructokinase, liver type OS=Homo sapiens OX=9606 GN=PFKL PE=1 SV=6 | PFKL |
| Ras-related C3 botulinum toxin substrate 1 OS=Homo sapiens OX=9606 GN=RAC1 PE=1 SV=1 | RAC1 |
| Protein phosphatase 1 regulatory subunit 12A OS=Homo sapiens OX=9606 GN=PPP1R12A PE=1 SV=1 | PPP1R12A |
| Mitogen-activated protein kinase 1 OS=Homo sapiens OX=9606 GN=MAPK1 PE=1 SV=3 | MAPK1 |

| | |
|---|---|
| Liprin-beta-1 OS=Homo sapiens OX=9606 GN=PPFIBP1 PE=1 SV=2 | PPFIBP1 |
| Mitogen-activated protein kinase 3 OS=Homo sapiens OX=9606 GN=MAPK3 PE=1 SV=4 | MAPK3 |
| Palladin OS=Homo sapiens OX=9606 GN=PALLD PE=1 SV=3 | PALLD |
| Glutamate dehydrogenase 1, mitochondrial OS=Homo sapiens OX=9606 GN=GLUD1 PE=1 SV=2 | GLUD1 |
| F-actin-capping protein subunit beta OS=Homo sapiens OX=9606 GN=CAPZB PE=1 SV=4 | CAPZB |
| Utrophin OS=Homo sapiens OX=9606 GN=UTRN PE=1 SV=2 | UTRN |
| Alanine--tRNA ligase, cytoplasmic OS=Homo sapiens OX=9606 GN=AARS1 PE=1 SV=2 | AARS1 |
| Eukaryotic translation initiation factor 3 subunit M OS=Homo sapiens OX=9606 GN=EIF3M PE=1 SV=1 | EIF3M |
| 14-3-3 protein gamma OS=Homo sapiens OX=9606 GN=YWHAG PE=1 SV=2 | YWHAG |
| Isoform 5 of RNA-binding protein 14 OS=Homo sapiens OX=9606 GN=RBM14 | RBM14 |
| Nidogen-1 OS=Homo sapiens OX=9606 GN=NID1 PE=1 SV=3 | NID1 |
| Pre-mRNA-processing-splicing factor 8 OS=Homo sapiens OX=9606 GN=PRPF8 PE=1 SV=2 | PRPF8 |
| Eukaryotic translation initiation factor 4 gamma 1 OS=Homo sapiens OX=9606 GN=EIF4G1 PE=1 SV=4 | EIF4G1 |
| Serine--tRNA ligase, cytoplasmic OS=Homo sapiens OX=9606 GN=SARS1 PE=1 SV=3 | SARS1 |
| Puromycin-sensitive aminopeptidase OS=Homo sapiens OX=9606 GN=NPEPPS PE=1 SV=2 | NPEPPS |
| Adenosylhomocysteinase OS=Homo sapiens OX=9606 GN=AHCY PE=1 SV=4 | AHCY |
| E3 SUMO-protein ligase RanBP2 OS=Homo sapiens OX=9606 GN=RANBP2 PE=1 SV=2 | RANBP2 |
| Polyadenylate-binding protein 1 OS=Homo sapiens OX=9606 GN=PABPC1 PE=1 SV=2 | PABPC1 |
| Serpin H1 OS=Homo sapiens OX=9606 GN=SERPINH1 PE=1 SV=2 | SERPINH1 |
| Isoform 3 of Heterogeneous nuclear ribonucleoprotein D0 OS=Homo sapiens OX=9606 GN=HNRNPD | HNRNPD |
| Proliferation-associated protein 2G4 OS=Homo sapiens OX=9606 GN=PA2G4 PE=1 SV=3 | PA2G4 |
| Protein unc-45 homolog A OS=Homo sapiens OX=9606 GN=UNC45A PE=1 SV=1 | UNC45A |
| T-complex protein 1 subunit eta OS=Homo sapiens OX=9606 GN=CCT7 PE=1 SV=2 | CCT7 |
| Exportin-2 OS=Homo sapiens OX=9606 GN=CSE1L PE=1 SV=3 | CSE1L |
| KH domain-containing, RNA-binding, signal transduction-associated protein 1 OS=Homo sapiens OX=9606 GN=KHDRBS1 PE=1 SV=1 | KHDRBS1 |
| Far upstream element-binding protein 3 OS=Homo sapiens OX=9606 GN=FUBP3 PE=1 SV=2 | FUBP3 |
| 5'-3' exoribonuclease 2 OS=Homo sapiens OX=9606 GN=XRN2 PE=1 SV=1 | XRN2 |
| Isoform 2 of Hypoxia up-regulated protein 1 OS=Homo sapiens OX=9606 GN=HYOU1 | HYOU1 |
| Splicing factor 3A subunit 3 OS=Homo sapiens OX=9606 GN=SF3A3 PE=1 SV=1 | SF3A3 |
| RuvB-like 2 OS=Homo sapiens OX=9606 GN=RUVBL2 PE=1 SV=3 | RUVBL2 |
| Ezrin OS=Homo sapiens OX=9606 GN=EZR PE=1 SV=4 | EZR |
| Far upstream element-binding protein 1 OS=Homo sapiens OX=9606 GN=FUBP1 PE=1 SV=3 | FUBP1 |
| RNA-binding protein FUS OS=Homo sapiens OX=9606 GN=FUS PE=1 SV=1 | FUS |
| Asparagine--tRNA ligase, cytoplasmic OS=Homo sapiens OX=9606 GN=NARS1 PE=1 SV=1 | NARS1 |
| Isoform 2 of Gelsolin OS=Homo sapiens OX=9606 GN=GSN | GSN |
| Glutamine--tRNA ligase OS=Homo sapiens OX=9606 GN=QARS1 PE=1 SV=1 | QARS1 |
| 1-phosphatidylinositol 4,5-bisphosphate phosphodiesterase beta-3 OS=Homo sapiens OX=9606 GN=PLCB3 PE=1 SV=2 | PLCB3 |
| 40S ribosomal protein S11 OS=Homo sapiens OX=9606 GN=RPS11 PE=1 SV=3 | RPS11 |
| Phenylalanine--tRNA ligase alpha subunit OS=Homo sapiens OX=9606 GN=FARSA PE=1 SV=3 | FARSA |
| Annexin A7 OS=Homo sapiens OX=9606 GN=ANXA7 PE=1 SV=3 | ANXA7 |
| Dedicator of cytokinesis protein 6 OS=Homo sapiens OX=9606 GN=DOCK6 PE=1 SV=3 | DOCK6 |
| Spermidine synthase OS=Homo sapiens OX=9606 GN=SRM PE=1 SV=1 | SRM |
| Fragile X mental retardation syndrome-related protein 1 OS=Homo sapiens OX=9606 GN=FXR1 PE=1 SV=3 | FXR1 |
| Tight junction protein ZO-2 OS=Homo sapiens OX=9606 GN=TJP2 PE=1 SV=2 | TJP2 |

| | |
|---|---|
| Ran GTPase-activating protein 1 OS=Homo sapiens OX=9606 GN=RANGAP1 PE=1 SV=1 | RANGAP1 |
| Acylamino-acid-releasing enzyme OS=Homo sapiens OX=9606 GN=APEH PE=1 SV=4 | APEH |
| Caveolae-associated protein 1 OS=Homo sapiens OX=9606 GN=CAVIN1 PE=1 SV=1 | CAVIN1 |
| Procollagen galactosyltransferase 1 OS=Homo sapiens OX=9606 GN=COLGALT1 PE=1 SV=1 | COLGALT1 |
| Eukaryotic translation elongation factor 1 epsilon-1 OS=Homo sapiens OX=9606 GN=EEF1E1 PE=1 SV=1 | EEF1E1 |
| 60S ribosomal protein L3 OS=Homo sapiens OX=9606 GN=RPL3 PE=1 SV=2 | RPL3 |
| Rho GDP-dissociation inhibitor 1 OS=Homo sapiens OX=9606 GN=ARHGDIA PE=1 SV=3 | ARHGDIA |
| Cysteine and glycine-rich protein 1 OS=Homo sapiens OX=9606 GN=CSRP1 PE=1 SV=3 | CSRP1 |
| E3 ubiquitin-protein ligase RNF213 OS=Homo sapiens OX=9606 GN=RNF213 PE=1 SV=3 | RNF213 |
| H/ACA ribonucleoprotein complex subunit DKC1 OS=Homo sapiens OX=9606 GN=DKC1 PE=1 SV=3 | DKC1 |
| Centrosomal protein of 170 kDa OS=Homo sapiens OX=9606 GN=CEP170 PE=1 SV=1 | CEP170 |
| Endophilin-A2 OS=Homo sapiens OX=9606 GN=SH3GL1 PE=1 SV=1 | SH3GL1 |
| Polyadenylate-binding protein 2 OS=Homo sapiens OX=9606 GN=PABPN1 PE=1 SV=3 | PABPN1 |
| Threonine--tRNA ligase 1, cytoplasmic OS=Homo sapiens OX=9606 GN=TARS1 PE=1 SV=3 | TARS1 |
| Elongation factor Tu, mitochondrial OS=Homo sapiens OX=9606 GN=TUFM PE=1 SV=2 | TUFM |
| GTP-binding nuclear protein Ran OS=Homo sapiens OX=9606 GN=RAN PE=1 SV=3 | RAN |
| 14-3-3 protein eta OS=Homo sapiens OX=9606 GN=YWHAH PE=1 SV=4 | YWHAH |
| TAR DNA-binding protein 43 OS=Homo sapiens OX=9606 GN=TARDBP PE=1 SV=1 | TARDBP |
| FACT complex subunit SPT16 OS=Homo sapiens OX=9606 GN=SUPT16H PE=1 SV=1 | SUPT16H |
| Isoform 2 of HLA class I histocompatibility antigen, A alpha chain OS=Homo sapiens OX=9606 GN=HLA-A | HLA-A |
| Stress-induced-phosphoprotein 1 OS=Homo sapiens OX=9606 GN=STIP1 PE=1 SV=1 | STIP1 |
| Beta-2-syntrophin OS=Homo sapiens OX=9606 GN=SNTB2 PE=1 SV=1 | SNTB2 |
| Unconventional myosin-Va OS=Homo sapiens OX=9606 GN=MYO5A PE=1 SV=2 | MYO5A |
| Importin-7 OS=Homo sapiens OX=9606 GN=IPO7 PE=1 SV=1 | IPO7 |
| Zinc finger CCCH-type antiviral protein 1 OS=Homo sapiens OX=9606 GN=ZC3HAV1 PE=1 SV=3 | ZC3HAV1 |
| Actin-related protein 2/3 complex subunit 1B OS=Homo sapiens OX=9606 GN=ARPC1B PE=1 SV=3 | ARPC1B |
| Kinectin OS=Homo sapiens OX=9606 GN=KTN1 PE=1 SV=1 | KTN1 |
| Abl interactor 1 OS=Homo sapiens OX=9606 GN=ABI1 PE=1 SV=4 | ABI1 |
| Atlastin-3 OS=Homo sapiens OX=9606 GN=ATL3 PE=1 SV=1 | ATL3 |
| Adenine phosphoribosyltransferase OS=Homo sapiens OX=9606 GN=APRT PE=1 SV=2 | APRT |
| Ataxin-2 OS=Homo sapiens OX=9606 GN=ATXN2 PE=1 SV=2 | ATXN2 |
| Splicing factor 3B subunit 3 OS=Homo sapiens OX=9606 GN=SF3B3 PE=1 SV=4 | SF3B3 |
| Poly(rC)-binding protein 1 OS=Homo sapiens OX=9606 GN=PCBP1 PE=1 SV=2 | PCBP1 |
| U5 small nuclear ribonucleoprotein 200 kDa helicase OS=Homo sapiens OX=9606 GN=SNRNP200 PE=1 SV=2 | SNRNP200 |
| Leucine-rich PPR motif-containing protein, mitochondrial OS=Homo sapiens OX=9606 GN=LRPPRC PE=1 SV=3 | LRPPRC |
| Vacuolar protein sorting-associated protein 35 OS=Homo sapiens OX=9606 GN=VPS35 PE=1 SV=2 | VPS35 |
| Lysosomal Pro-X carboxypeptidase OS=Homo sapiens OX=9606 GN=PRCP PE=1 SV=1 | PRCP |
| Cullin-associated NEDD8-dissociated protein 1 OS=Homo sapiens OX=9606 GN=CAND1 PE=1 SV=2 | CAND1 |
| Guanine nucleotide-binding protein G(I)/G(S)/G(T) subunit beta-1 OS=Homo sapiens OX=9606 GN=GNB1 PE=1 SV=3 | GNB1 |
| Protein PML OS=Homo sapiens OX=9606 GN=PML PE=1 SV=3 | PML |
| Multimerin-2 OS=Homo sapiens OX=9606 GN=MMRN2 PE=1 SV=2 | MMRN2 |
| Eukaryotic translation initiation factor 4 gamma 2 OS=Homo sapiens OX=9606 GN=EIF4G2 PE=1 SV=1 | EIF4G2 |

| | |
|---|---|
| Isoform 6 of Thioredoxin reductase 1, cytoplasmic OS=Homo sapiens OX=9606 GN=TXNRD1 | TXNRD1 |
| Dynamin-2 OS=Homo sapiens OX=9606 GN=DNM2 PE=1 SV=2 | DNM2 |
| Heterogeneous nuclear ribonucleoprotein A3 OS=Homo sapiens OX=9606 GN=HNRNPA3 PE=1 SV=2 | HNRNPA3 |
| DNA damage-binding protein 1 OS=Homo sapiens OX=9606 GN=DDB1 PE=1 SV=1 | DDB1 |
| Tropomyosin alpha-4 chain OS=Homo sapiens OX=9606 GN=TPM4 PE=1 SV=3 | TPM4 |
| Importin-5 OS=Homo sapiens OX=9606 GN=IPO5 PE=1 SV=4 | IPO5 |
| 182 kDa tankyrase-1-binding protein OS=Homo sapiens OX=9606 GN=TNKS1BP1 PE=1 SV=4 | TNKS1BP1 |
| Toll-interacting protein OS=Homo sapiens OX=9606 GN=TOLLIP PE=1 SV=1 | TOLLIP |
| Xaa-Pro aminopeptidase 1 OS=Homo sapiens OX=9606 GN=XPNPEP1 PE=1 SV=3 | XPNPEP1 |
| Integrin-linked protein kinase OS=Homo sapiens OX=9606 GN=ILK PE=1 SV=2 | ILK |
| Caldesmon OS=Homo sapiens OX=9606 GN=CALD1 PE=1 SV=3 | CALD1 |
| Rho-related GTP-binding protein RhoC OS=Homo sapiens OX=9606 GN=RHOC PE=1 SV=1 | RHOC |
| Serine/arginine-rich splicing factor 6 OS=Homo sapiens OX=9606 GN=SRSF6 PE=1 SV=2 | SRSF6 |
| ATP-binding cassette sub-family E member 1 OS=Homo sapiens OX=9606 GN=ABCE1 PE=1 SV=1 | ABCE1 |
| Leucine zipper protein 1 OS=Homo sapiens OX=9606 GN=LUZP1 PE=1 SV=2 | LUZP1 |
| Ras-related protein Rap-1b-like protein OS=Homo sapiens OX=9606 PE=2 SV=1 | RP1BL |
| Twinfilin-1 OS=Homo sapiens OX=9606 GN=TWF1 PE=1 SV=3 | TWF1 |
| 26S proteasome non-ATPase regulatory subunit 12 OS=Homo sapiens OX=9606 GN=PSMD12 PE=1 SV=3 | PSMD12 |
| Pre-mRNA-processing factor 19 OS=Homo sapiens OX=9606 GN=PRPF19 PE=1 SV=1 | PRPF19 |
| Interleukin enhancer-binding factor 3 OS=Homo sapiens OX=9606 GN=ILF3 PE=1 SV=3 | ILF3 |
| Guanine nucleotide-binding protein G(I)/G(S)/G(T) subunit beta-2 OS=Homo sapiens OX=9606 GN=GNB2 PE=1 SV=3 | GNB2 |
| Y-box-binding protein 1 OS=Homo sapiens OX=9606 GN=YBX1 PE=1 SV=3 | YBX1 |
| RNA-splicing ligase RtcB homolog OS=Homo sapiens OX=9606 GN=RTCB PE=1 SV=1 | RTCB |
| Exosome complex component RRP40 OS=Homo sapiens OX=9606 GN=EXOSC3 PE=1 SV=3 | EXOSC3 |
| Calpain small subunit 1 OS=Homo sapiens OX=9606 GN=CAPNS1 PE=1 SV=1 | CAPNS1 |
| 26S proteasome regulatory subunit 7 OS=Homo sapiens OX=9606 GN=PSMC2 PE=1 SV=3 | PSMC2 |
| RNA-binding protein 14 OS=Homo sapiens OX=9606 GN=RBM14 PE=1 SV=2 | RBM14 |
| Adenylyl cyclase-associated protein 1 OS=Homo sapiens OX=9606 GN=CAP1 PE=1 SV=5 | CAP1 |
| 60S ribosomal protein L7a OS=Homo sapiens OX=9606 GN=RPL7A PE=1 SV=2 | RPL7A |
| Carnitine O-palmitoyltransferase 1, liver isoform OS=Homo sapiens OX=9606 GN=CPT1A PE=1 SV=2 | CPT1A |
| Stathmin OS=Homo sapiens OX=9606 GN=STMN1 PE=1 SV=3 | STMN1 |
| Inosine-5'-monophosphate dehydrogenase 2 OS=Homo sapiens OX=9606 GN=IMPDH2 PE=1 SV=2 | IMPDH2 |
| 14-3-3 protein epsilon OS=Homo sapiens OX=9606 GN=YWHAE PE=1 SV=1 | YWHAE |
| Signal recognition particle subunit SRP68 OS=Homo sapiens OX=9606 GN=SRP68 PE=1 SV=2 | SRP68 |
| Isoform 2 of Macrophage-capping protein OS=Homo sapiens OX=9606 GN=CAPG | CAPG |
| Hsp90 co-chaperone Cdc37 OS=Homo sapiens OX=9606 GN=CDC37 PE=1 SV=1 | CDC37 |
| Core histone macro-H2A.1 OS=Homo sapiens OX=9606 GN=MACROH2A1 PE=1 SV=4 | MACROH2A1 |
| Dynein axonemal assembly factor 5 OS=Homo sapiens OX=9606 GN=DNAAF5 PE=1 SV=4 | DNAAF5 |
| Dihydrolipoyllysine-residue acetyltransferase component of pyruvate dehydrogenase complex, mitochondrial OS=Homo sapiens OX=9606 GN=DLAT PE=1 SV=3 | DLAT |
| 40S ribosomal protein S3 OS=Homo sapiens OX=9606 GN=RPS3 PE=1 SV=2 | RPS3 |
| Histone H2B type 1-O OS=Homo sapiens OX=9606 GN=H2BC17 PE=1 SV=3 | H2BC17 |

| | |
|---|---|
| Isoform 3 of LIM domain only protein 7 OS=Homo sapiens OX=9606 GN=LMO7 | LMO7 |
| Coiled-coil domain-containing protein 22 OS=Homo sapiens OX=9606 GN=CCDC22 PE=1 SV=1 | CCDC22 |
| GRB10-interacting GYF protein 2 OS=Homo sapiens OX=9606 GN=GIGYF2 PE=1 SV=1 | GIGYF2 |
| Glutamine synthetase OS=Homo sapiens OX=9606 GN=GLUL PE=1 SV=4 | GLUL |
| Keratin, type I cytoskeletal 9 OS=Homo sapiens OX=9606 GN=KRT9 PE=1 SV=3 | KRT9 |
| Arginine--tRNA ligase, cytoplasmic OS=Homo sapiens OX=9606 GN=RARS1 PE=1 SV=2 | RARS1 |
| Hexokinase-1 OS=Homo sapiens OX=9606 GN=HK1 PE=1 SV=3 | HK1 |
| Adenylosuccinate synthetase isozyme 2 OS=Homo sapiens OX=9606 GN=ADSS2 PE=1 SV=3 | ADSS2 |
| Serine/threonine-protein phosphatase 2A 65 kDa regulatory subunit A alpha isoform OS=Homo sapiens OX=9606 GN=PPP2R1A PE=1 SV=4 | PPP2R1A |
| PDZ and LIM domain protein 5 OS=Homo sapiens OX=9606 GN=PDLIM5 PE=1 SV=5 | PDLIM5 |
| Calpain-1 catalytic subunit OS=Homo sapiens OX=9606 GN=CAPN1 PE=1 SV=1 | CAPN1 |
| Isoform 3 of Septin-2 OS=Homo sapiens OX=9606 GN=SEPTIN2 | SEPTIN2 |
| Glucosidase 2 subunit beta OS=Homo sapiens OX=9606 GN=PRKCSH PE=1 SV=2 | PRKCSH |
| N-acetylglucosamine-6-sulfatase OS=Homo sapiens OX=9606 GN=GNS PE=1 SV=3 | GNS |
| Myosin-10 OS=Homo sapiens OX=9606 GN=MYH10 PE=1 SV=3 | MYH10 |
| Heterogeneous nuclear ribonucleoprotein Q OS=Homo sapiens OX=9606 GN=SYNCRIP PE=1 SV=2 | SYNCRIP |
| Endoglin OS=Homo sapiens OX=9606 GN=ENG PE=1 SV=2 | ENG |
| Heterogeneous nuclear ribonucleoprotein U-like protein 1 OS=Homo sapiens OX=9606 GN=HNRNPUL1 PE=1 SV=2 | HNRNPUL1 |
| 3-mercaptopyruvate sulfurtransferase OS=Homo sapiens OX=9606 GN=MPST PE=1 SV=3 | MPST |
| Tripeptidyl-peptidase 1 OS=Homo sapiens OX=9606 GN=TPP1 PE=1 SV=2 | TPP1 |
| Regulator of nonsense transcripts 1 OS=Homo sapiens OX=9606 GN=UPF1 PE=1 SV=2 | UPF1 |
| Keratin, type II cytoskeletal 2 epidermal OS=Homo sapiens OX=9606 GN=KRT2 PE=1 SV=2 | KRT2 |
| Tyrosine--tRNA ligase, cytoplasmic OS=Homo sapiens OX=9606 GN=YARS1 PE=1 SV=4 | YARS1 |
| Trifunctional purine biosynthetic protein adenosine-3 OS=Homo sapiens OX=9606 GN=GART PE=1 SV=1 | GART |
| Isoform Non-brain of Clathrin light chain A OS=Homo sapiens OX=9606 GN=CLTA | CLTA |
| Multifunctional procollagen lysine hydroxylase and glycosyltransferase LH3 OS=Homo sapiens OX=9606 GN=PLOD3 PE=1 SV=1 | PLOD3 |
| Plasminogen activator inhibitor 1 RNA-binding protein OS=Homo sapiens OX=9606 GN=SERBP1 PE=1 SV=2 | SERBP1 |
| 40S ribosomal protein S6 OS=Homo sapiens OX=9606 GN=RPS6 PE=1 SV=1 | RPS6 |
| E3 ubiquitin-protein ligase HUWE1 OS=Homo sapiens OX=9606 GN=HUWE1 PE=1 SV=3 | HUWE1 |
| Serine/arginine repetitive matrix protein 2 OS=Homo sapiens OX=9606 GN=SRRM2 PE=1 SV=2 | SRRM2 |
| Nitric oxide synthase, endothelial OS=Homo sapiens OX=9606 GN=NOS3 PE=1 SV=4 | NOS3 |
| Calpastatin OS=Homo sapiens OX=9606 GN=CAST PE=1 SV=4 | CAST |
| Protein kinase C alpha type OS=Homo sapiens OX=9606 GN=PRKCA PE=1 SV=4 | PRKCA |
| ATP-dependent RNA helicase DHX15 OS=Homo sapiens OX=9606 GN=DHX15 PE=1 SV=2 | DHX15 |
| TRIO and F-actin-binding protein OS=Homo sapiens OX=9606 GN=TRIOBP PE=1 SV=3 | TRIOBP |
| Nuclear protein localization protein 4 homolog OS=Homo sapiens OX=9606 GN=NPLOC4 PE=1 SV=3 | NPLOC4 |
| Isoform 4 of Plectin OS=Homo sapiens OX=9606 GN=PLEC | PLEC |
| Heat shock 70 kDa protein 1A OS=Homo sapiens OX=9606 GN=HSPA1A PE=1 SV=1 | HSPA1A |
| Poly [ADP-ribose] polymerase 1 OS=Homo sapiens OX=9606 GN=PARP1 PE=1 SV=4 | PARP1 |
| Glutaredoxin-3 OS=Homo sapiens OX=9606 GN=GLRX3 PE=1 SV=2 | GLRX3 |
| Isoform 2 of UDP-glucose:glycoprotein glucosyltransferase 1 OS=Homo sapiens OX=9606 GN=UGGT1 | UGGT1 |
| Dynamin-binding protein OS=Homo sapiens OX=9606 GN=DNMBP PE=1 SV=1 | DNMBP |

| | |
|---|---|
| Enhancer of mRNA-decapping protein 4 OS=Homo sapiens OX=9606 GN=EDC4 PE=1 SV=1 | EDC4 |
| Alkyldihydroxyacetonephosphate synthase, peroxisomal OS=Homo sapiens OX=9606 GN=AGPS PE=1 SV=1 | AGPS |
| 1,4-alpha-glucan-branching enzyme OS=Homo sapiens OX=9606 GN=GBE1 PE=1 SV=3 | GBE1 |
| 40S ribosomal protein S18 OS=Homo sapiens OX=9606 GN=RPS18 PE=1 SV=3 | RPS18 |
| cAMP-dependent protein kinase type II-alpha regulatory subunit OS=Homo sapiens OX=9606 GN=PRKAR2A PE=1 SV=2 | PRKAR2A |
| Vacuolar protein sorting-associated protein 52 homolog OS=Homo sapiens OX=9606 GN=VPS52 PE=1 SV=1 | VPS52 |
| Heat shock 70 kDa protein 4 OS=Homo sapiens OX=9606 GN=HSPA4 PE=1 SV=4 | HSPA4 |
| RNA-binding protein 39 OS=Homo sapiens OX=9606 GN=RBM39 PE=1 SV=2 | RBM39 |
| Actin-related protein 3 OS=Homo sapiens OX=9606 GN=ACTR3 PE=1 SV=3 | ACTR3 |
| Coatomer subunit beta OS=Homo sapiens OX=9606 GN=COPB1 PE=1 SV=3 | COPB1 |
| Protein transport protein Sec31A OS=Homo sapiens OX=9606 GN=SEC31A PE=1 SV=3 | SEC31A |
| Heterochromatin protein 1-binding protein 3 OS=Homo sapiens OX=9606 GN=HP1BP3 PE=1 SV=1 | HP1BP3 |
| Nucleolar protein 58 OS=Homo sapiens OX=9606 GN=NOP58 PE=1 SV=1 | NOP58 |
| Serine/arginine-rich splicing factor 7 OS=Homo sapiens OX=9606 GN=SRSF7 PE=1 SV=1 | SRSF7 |
| Filamin-binding LIM protein 1 OS=Homo sapiens OX=9606 GN=FBLIM1 PE=1 SV=2 | FBLIM1 |
| Nuclear pore complex protein Nup153 OS=Homo sapiens OX=9606 GN=NUP153 PE=1 SV=2 | NUP153 |
| Inverted formin-2 OS=Homo sapiens OX=9606 GN=INF2 PE=1 SV=2 | INF2 |
| Aminopeptidase B OS=Homo sapiens OX=9606 GN=RNPEP PE=1 SV=2 | RNPEP |
| Aldehyde dehydrogenase, mitochondrial OS=Homo sapiens OX=9606 GN=ALDH2 PE=1 SV=2 | ALDH2 |
| DNA-directed RNA polymerase II subunit RPB1 OS=Homo sapiens OX=9606 GN=POLR2A PE=1 SV=2 | POLR2A |

**S10: Identified NC9-treated 3F-TG2-associated proteins from HUVEC using anti-Flag M2 agarose resin for the CoIP (NC9-treated TG2 is considered as an opened TG2; n=3)**

| Gene Name | Name of proteins |
|---|---|
| ABCD1 | ATP-binding cassette sub-family D member 1 |
| ABCD3 | ATP-binding cassette sub-family D member 3 |
| ABLIM1 | Actin-binding LIM protein 1 |
| ACAP2 | Arf-GAP with coiled-coil, ANK repeat and PH domain-containing protein 2 |
| ACIN1 | Apoptotic chromatin condensation inducer in the nucleus |
| ACTN1 | Alpha-actinin-1 |
| ACTR1A | Alpha-centractin |
| ADRM1 | Proteasomal ubiquitin receptor ADRM1 |
| AHNAK | Neuroblast differentiation-associated protein AHNAK |
| ALDOA | Fructose-bisphosphate aldolase A |
| ANKFY1 | Rabankyrin-5 |
| ANO10 | Anoctamin-10 |
| ANPEP | Aminopeptidase N |
| ANXA5 | Annexin A5 |
| ANXA6 | Annexin A6 |
| AP2A1 | AP-2 complex subunit alpha-1 |
| AP2B1 | AP-2 complex subunit beta |
| APOA1 | Apolipoprotein A-I |
| ARCN1 | Coatomer subunit delta |
| ATAD3A | Isoform 2 of ATPase family AAA domain-containing protein 3A |
| ATP13A1 | Endoplasmic reticulum transmembrane helix translocase |
| ATP1A1 | Sodium/potassium-transporting ATPase subunit alpha-1 |
| ATP2A2 | Sarcoplasmic/endoplasmic reticulum calcium ATPase 2 |
| ATP5F1C | ATP synthase subunit gamma, mitochondrial |
| ATP6V0A1 | V-type proton ATPase 116 kDa subunit a 1 |
| ATP6V0A2 | V-type proton ATPase 116 kDa subunit a 2 |
| ATP6V0D1 | V-type proton ATPase subunit d 1 |
| ATP6V1B2 | V-type proton ATPase subunit B, brain isoform |
| BCLAF1 | Bcl-2-associated transcription factor 1 |
| C3 | Complement C3 |
| CALR | Calreticulin |
| CAP1 | Adenylyl cyclase-associated protein 1 |
| CAPZA1 | F-actin-capping protein subunit alpha-1 |
| CAPZB | F-actin-capping protein subunit beta |
| CAV1 | Caveolin-1 |
| CAVIN1 | Caveolae-associated protein 1 |
| CCAR1 | Cell division cycle and apoptosis regulator protein 1 |
| CCT2 | T-complex protein 1 subunit beta |
| CCT3 | T-complex protein 1 subunit gamma |
| CCT5 | T-complex protein 1 subunit epsilon |
| CCT7 | T-complex protein 1 subunit eta |
| CCT8 | T-complex protein 1 subunit theta |
| CD44 | CD44 antigen |
| CD55 | Complement decay-accelerating factor |
| CD59 | CD59 glycoprotein |
| CDC42 | Cell division control protein 42 homolog |
| CHCHD3 | MICOS complex subunit MIC19 |
| CKAP4 | Cytoskeleton-associated protein 4 |
| COPB2 | Coatomer subunit beta' |
| COPG1 | Coatomer subunit gamma-1 |
| CPT1A | Carnitine O-palmitoyltransferase 1, liver isoform |
| CTNND1 | Catenin delta-1 |
| CSE1L | Exportin-2 |
| DCTN1 | Isoform 3 of Dynactin subunit 1 |
| DDOST | Dolichyl-diphosphooligosaccharide--protein glycosyltransferase 48 kDa subunit |
| DDX23 | Probable ATP-dependent RNA helicase DDX23 |
| DDX39B | Spliceosome RNA helicase DDX39B |
| DHX15 | ATP-dependent RNA helicase DHX15 |
| DKC1 | H/ACA ribonucleoprotein complex subunit DKC1 |
| DLD | Dihydrolipoyl dehydrogenase, mitochondrial |
| DLST | Dihydrolipoyllysine-residue succinyltransferase component of 2-oxoglutarate dehydrogenase complex, mitochondrial |
| DPP7 | Dipeptidyl peptidase 2 |
| DPY19L1 | Probable C-mannosyltransferase DPY19L1 |
| DYNC1H1 | Cytoplasmic dynein 1 heavy chain 1 |
| EEF1D | Isoform 2 of Elongation factor 1-delta |
| EEF1G | Elongation factor 1-gamma |
| EFTUD2 | 116 kDa U5 small nuclear ribonucleoprotein component |
| EHD4 | EH domain-containing protein 4 |
| EIF3A | Eukaryotic translation initiation factor 3 subunit A |

| | |
|---|---|
| EIF3B | Isoform 2 of Eukaryotic translation initiation factor 3 subunit B |
| EIF3CL | Eukaryotic translation initiation factor 3 subunit C-like protein |
| EIF3E | Eukaryotic translation initiation factor 3 subunit E |
| EIF3I | Eukaryotic translation initiation factor 3 subunit I |
| EIF3L | Eukaryotic translation initiation factor 3 subunit L |
| EIF4A3 | Eukaryotic initiation factor 4A-III |
| EIF4B | Eukaryotic translation initiation factor 4B |
| EMC1 | ER membrane protein complex subunit 1 |
| EMC2 | ER membrane protein complex subunit 2 |
| ENDOD1 | Endonuclease domain-containing 1 protein |
| ENG | Endoglin |
| ENO1 | Alpha-enolase |
| ERLIN1 | Erlin-1 |
| ERLIN2 | Erlin-2 |
| ERMP1 | Endoplasmic reticulum metallopeptidase 1 |
| FBL | rRNA 2'-O-methyltransferase fibrillarin |
| FERMT3 | Fermitin family homolog 3 |
| FLII | Protein flightless-1 homolog |
| FLOT1 | Flotillin-1 |
| FLOT2 | Flotillin-2 |
| FMNL2 | Formin-like protein 2 |
| FSCN1 | Fascin |
| FUT8 | Alpha-(1,6)-fucosyltransferase |
| FXR1 | Fragile X mental retardation syndrome-related protein 1 |
| GANAB | Neutral alpha-glucosidase AB |
| GDI2 | Rab GDP dissociation inhibitor beta |
| GLG1 | Isoform 3 of Golgi apparatus protein 1 |
| GLIPR2 | Golgi-associated plant pathogenesis-related protein 1 |
| GNAI2 | Guanine nucleotide-binding protein G(i) subunit alpha-2 |
| GNB1 | Guanine nucleotide-binding protein G(I)/G(S)/G(T) subunit beta-1 |
| GNB2 | Guanine nucleotide-binding protein G(I)/G(S)/G(T) subunit beta-2 |
| GNG12 | Guanine nucleotide-binding protein G(I)/G(S)/G(O) subunit gamma-12 |
| GORASP2 | Isoform 3 of Golgi reassembly-stacking protein 2 |
| H1-0 | Histone H1.0 |
| H2AC6 | Histone H2A type 1-C |
| H2AX | Histone H2A.X |
| H2BC4 | Histone H2B type 1-C/E/F/G/I |
| HCLS1 | Hematopoietic lineage cell-specific protein |
| HDLBP | Vigilin |
| HMGA2 | High mobility group protein HMGI-C |
| HP1BP3 | Heterochromatin protein 1-binding protein 3 |
| HSP90AA1 | Heat shock protein HSP 90-alpha |
| HSPA9 | Stress-70 protein, mitochondrial |
| IFI16 | Gamma-interferon-inducible protein 16 |
| IGKV2D-24 | Probable non-functional immunoglobulin kappa variable 2D-24 |
| IGKV2D-29 | Immunoglobulin kappa variable 2D-29 |
| ILF2 | Interleukin enhancer-binding factor 2 |
| IMMT | MICOS complex subunit MIC60 |
| IQGAP1 | Ras GTPase-activating-like protein IQGAP1 |
| ITGA6 | Integrin alpha-6 |
| ITGB1 | Integrin beta-1 |
| ITGB4 | Isoform Beta-4D of Integrin beta-4 |
| ITPR3 | Inositol 1,4,5-trisphosphate receptor type 3 |
| JAK1 | Tyrosine-protein kinase JAK1 |
| KCTD5 | BTB/POZ domain-containing protein KCTD5 |
| KIF11 | Kinesin-like protein KIF11 |
| KIF5B | Kinesin-1 heavy chain |
| KPNB1 | Importin subunit beta-1 |
| KRT14 | Keratin, type I cytoskeletal 14 |
| KRT18 | Keratin, type I cytoskeletal 18 |
| KRT2 | Keratin, type II cytoskeletal 18 |
| KRT7 | Keratin, type I cytoskeletal 7 |
| KTN1 | Kinectin |
| LBR | Delta(14)-sterol reductase LBR |
| LCP1 | Plastin-2 |
| LDHB | L-lactate dehydrogenase B chain |
| LMAN1 | Protein ERGIC-53 |
| LMNB2 | Lamin-B2 |
| LRPPRC | Leucine-rich PPR motif-containing protein, mitochondrial |
| LRRC8A | Volume-regulated anion channel subunit LRRC8A |
| LRRC8C | Volume-regulated anion channel subunit LRRC8C |
| MAGT1 | Magnesium transporter protein 1 |
| MAP1B | Microtubule-associated protein 1B |
| MBOAT7 | Lysophospholipid acyltransferase 7 |

| | |
|---|---|
| MCAM | Cell surface glycoprotein MUC18 |
| MGST3 | Microsomal glutathione S-transferase 3 |
| MLEC | Malectin |
| MSN | Moesin |
| MTDH | Protein LYRIC |
| MYADM | Myeloid-associated differentiation marker |
| MYH10 | Myosin-10 |
| MYH14 | Myosin-14 |
| MYL6 | Myosin light polypeptide 6 |
| MYOF | Myoferlin |
| NACA | Nascent polypeptide-associated complex subunit alpha, muscle-specific form |
| NES | Nestin |
| NNT | NAD(P) transhydrogenase, mitochondrial |
| NOP2 | Probable 28S rRNA (cytosine(4447)-C(5))-methyltransferase |
| NOP56 | Nucleolar protein 56 |
| NOP58 | Nucleolar protein 58 |
| NT5E | 5'-nucleotidase |
| OCIAD2 | OCIA domain-containing protein 2 |
| OSBPL3 | Oxysterol-binding protein-related protein 3 |
| PA2G4 | Proliferation-associated protein 2G4 |
| PCBP1 | Poly(rC)-binding protein 1 |
| PCBP2 | Poly(rC)-binding protein 2 |
| PDCD6IP | Programmed cell death 6-interacting protein |
| PDIA3 | Protein disulfide-isomerase A3 |
| PFKP | ATP-dependent 6-phosphofructokinase, platelet type |
| PGAM5 | Serine/threonine-protein phosphatase PGAM5, mitochondrial |
| PHB2 | Prohibitin-2 |
| PKM | Pyruvate kinase PKM |
| PLAT | Tissue-type plasminogen activator |
| PLOD2 | Isoform 2 of Procollagen-lysine,2-oxoglutarate 5-dioxygenase 2 |
| PPM1B | Protein phosphatase 1B |
| PRMT5 | Protein arginine N-methyltransferase 5 |
| PRPF31 | U4/U6 small nuclear ribonucleoprotein Prp31 |
| PRPF6 | Pre-mRNA-processing factor 6 |
| PRPF8 | Pre-mRNA-processing-splicing factor 8 |
| PRPS1 | Ribose-phosphate pyrophosphokinase 1 |
| PRPSAP1 | Phosphoribosyl pyrophosphate synthase-associated protein 1 |
| PTDSS1 | Phosphatidylserine synthase 1 |
| RAB10 | Ras-related protein Rab-10 |
| RAB11B | Ras-related protein Rab-11B |
| RAB5C | Ras-related protein Rab-5C |
| RAB7A | Ras-related protein Rab-7a |
| RAC2 | Ras-related C3 botulinum toxin substrate 2 |
| RACK1 | Receptor of activated protein C kinase 1 |
| RAI14 | Ankycorbin |
| RALA | Ras-related protein Ral-A |
| RALY | RNA-binding protein Raly |
| RAN | GTP-binding nuclear protein Ran |
| RAP1A | Ras-related protein Rap-1A |
| RBBP4 | Histone-binding protein RBBP4 |
| RBM10 | RNA-binding protein 10 |
| RER1 | Protein RER1 |
| RHOC | Rho-related GTP-binding protein RhoC |
| RNH1 | Ribonuclease inhibitor |
| ROCK1 | Rho-associated protein kinase 1 |
| RPL10A | 60S ribosomal protein L10a |
| RPL11 | 60S ribosomal protein L11 |
| RPL12 | 60S ribosomal protein L12 |
| RPL13 | 60S ribosomal protein L13 |
| RPL13A | 60S ribosomal protein L13a |
| RPL14 | 60S ribosomal protein L14 |
| RPL17 | 60S ribosomal protein L17 |
| RPL18 | 60S ribosomal protein L18 |
| RPL23 | 60S ribosomal protein L23 |
| RPL29 | 60S ribosomal protein L29 |
| RPL3 | 60S ribosomal protein L3 |
| RPL5 | 60S ribosomal protein L5 |
| RPL8 | 60S ribosomal protein L8 |
| RPLP0 | 60S acidic ribosomal protein P0 |
| RPN1 | Dolichyl-diphosphooligosaccharide--protein glycosyltransferase subunit 1 |
| RPN2 | Dolichyl-diphosphooligosaccharide--protein glycosyltransferase subunit 2 |
| RPS19 | 40S ribosomal protein S19 |
| RPS25 | 40S ribosomal protein S25 |
| RPS26 | 40S ribosomal protein S26 |

| | |
|---|---|
| RPS3 | 40S ribosomal protein S3 |
| RPS3A | 40S ribosomal protein S3a |
| RPS4X | 40S ribosomal protein S4, X isoform |
| RPS7 | 40S ribosomal protein S7 |
| RPSA | 40S ribosomal protein SA |
| RRAS | Ras-related protein R-Ras |
| RRBP1 | Ribosome-binding protein 1 |
| RSL1D1 | Ribosomal L1 domain-containing protein 1 |
| SAMM50 | Sorting and assembly machinery component 50 homolog |
| SEC11A | Signal peptidase complex catalytic subunit SEC11A |
| SEC22B | Vesicle-trafficking protein SEC22b |
| SEC62 | Translocation protein SEC62 |
| SERPINE1 | Plasminogen activator inhibitor 1 |
| SF3B1 | Splicing factor 3B subunit 1 |
| SF3B2 | Splicing factor 3B subunit 2 |
| SFXN1 | Sideroflexin-1 |
| SFXN3 | Sideroflexin-3 |
| SLC12A4 | Solute carrier family 12 member 4 |
| SLC25A11 | Mitochondrial 2-oxoglutarate/malate carrier protein |
| SLC2A1 | Solute carrier family 2, facilitated glucose transporter member 1 |
| SLC35B2 | Adenosine 3'-phospho 5'-phosphosulfate transporter 1 |
| SLTM | SAFB-like transcription modulator |
| SND1 | Staphylococcal nuclease domain-containing protein 1 |
| SNRNP200 | U5 small nuclear ribonucleoprotein 200 kDa helicase |
| SNRPD1 | Small nuclear ribonucleoprotein Sm D1 |
| SRSF1 | Serine/arginine-rich splicing factor 1 |
| SRSF6 | Serine/arginine-rich splicing factor 6 |
| SRSF7 | Isoform 2 of Serine/arginine-rich splicing factor 7 |
| SSRP1 | FACT complex subunit SSRP1 |
| STING1 | Stimulator of interferon genes protein |
| STK38 | Serine/threonine-protein kinase 38 |
| STOM | Stomatin |
| STT3A | Dolichyl-diphosphooligosaccharide--protein glycosyltransferase subunit STT3A |
| STT3B | Dolichyl-diphosphooligosaccharide--protein glycosyltransferase subunit STT3B |
| SUN2 | Isoform 2 of SUN domain-containing protein 2 |
| SUPT16H | FACT complex subunit SPT16 |
| SVIL | Supervillin |
| TAB1 | TGF-beta-activated kinase 1 and MAP3K7-binding protein 1 |
| TAF4 | Transcription initiation factor TFIID subunit 4 |
| TAGLN2 | Transgelin-2 |
| TALDO1 | Transaldolase |
| TBL2 | Transducin beta-like protein 2 |
| TCIRG1 | V-type proton ATPase 116 kDa subunit a 3 |
| TCP1 | T-complex protein 1 subunit alpha |
| TGM2 | Protein-glutamine gamma-glutamyltransferase 2 |
| THRAP3 | Thyroid hormone receptor-associated protein 3 |
| TICAM2 | Isoform 2 of TIR domain-containing adapter molecule 2 |
| TLN1 | Talin-1 |
| TMED10 | Transmembrane emp24 domain-containing protein 10 |
| TMED9 | Transmembrane emp24 domain-containing protein 9 |
| TMEM43 | Transmembrane protein 43 |
| TMPO | Lamina-associated polypeptide 2, isoforms beta/gamma |
| TMTC3 | Protein O-mannosyl-transferase TMTC3 |
| TPI1 | Isoform 2 of Triosephosphate isomerase |
| TRA2B | Transformer-2 protein homolog beta |
| TRIM21 | E3 ubiquitin-protein ligase TRIM21 |
| TRPV2 | Transient receptor potential cation channel subfamily V member 2 |
| TUBA1A | Tubulin alpha-1A chain |
| TUBB4A | Tubulin beta-4A chain |
| TXNDC5 | Thioredoxin domain-containing protein 5 |
| UBA52 | Ubiquitin-60S ribosomal protein L40 |
| UQCRC1 | Cytochrome b-c1 complex subunit 1, mitochondrial |
| UQCRC2 | Cytochrome b-c1 complex subunit 2, mitochondrial |
| VAPA | Isoform 2 of Vesicle-associated membrane protein-associated protein A |
| VAT1 | Synaptic vesicle membrane protein VAT-1 homolog |
| VDAC1 | Voltage-dependent anion-selective channel protein 1 |
| VDAC2 | Isoform 1 of Voltage-dependent anion-selective channel protein 2 |
| VDAC3 | Voltage-dependent anion-selective channel protein 3 |
| VPS35 | Vacuolar protein sorting-associated protein 35 |
| WARS1 | Tryptophan--tRNA ligase, cytoplasmic |
| WDR1 | WD repeat-containing protein 1 |
| WDR77 | Methylosome protein 50 |
| XRCC5 | X-ray repair cross-complementing protein 5 |
| YBX1 | Y-box-binding protein 1 |

| | |
|---|---|
| YES1 | Tyrosine-protein kinase Yes |
| YWHAZ | 14-3-3 protein zeta/delta |
| ZDHHC13 | Palmitoyltransferase ZDHHC13 |
| ZMPSTE24 | CAAX prenyl protease 1 homolog |

| S11: Identified 3F-TG2-associated proteins in the presence of GTPγS from HUVEC using anti-Flag M2 agarose resin for the CoIP (in the presence of GTPγS TG2 is considered as a closed TG2; n=3) | |
|---|---|
| **Gene Name** | **Name of proteins** |
| ABLIM1 | Actin-binding LIM protein 1 |
| ACAP2 | Arf-GAP with coiled-coil, ANK repeat and PH domain-containing protein 2 |
| ACIN1 | Apoptotic chromatin condensation inducer in the nucleus |
| ACTN1 | Alpha-actinin-1 |
| ACTR1A | Alpha-centractin |
| ADRM1 | Proteasomal ubiquitin receptor ADRM1 |
| AHNAK | Neuroblast differentiation-associated protein AHNAK |
| ALB | Albumin |
| ALDOA | Fructose-bisphosphate aldolase A |
| ALYREF | THO complex subunit 4 |
| ANKFY1 | Rabankyrin-5 |
| ANPEP | Aminopeptidase N |
| ANXA5 | Annexin A5 |
| ANXA6 | Annexin A6 |
| BCLAF1 | Bcl-2-associated transcription factor 1 |
| CALR | Calreticulin |
| CAP1 | Adenylyl cyclase-associated protein 1 |
| CAPZA1 | F-actin-capping protein subunit alpha-1 |
| CAVIN1 | Caveolae-associated protein 1 |
| CCT2 | T-complex protein 1 subunit beta |
| CCT3 | T-complex protein 1 subunit gamma |
| CCT5 | T-complex protein 1 subunit epsilon |
| CCT6A | T-complex protein 1 subunit zeta |
| CCT7 | T-complex protein 1 subunit eta |
| CCT8 | T-complex protein 1 subunit theta |
| CD59 | CD59 glycoprotein |
| CDC42 | Cell division control protein 42 homolog |
| CKAP4 | Cytoskeleton-associated protein 4 |
| COPG1 | Coatomer subunit gamma-1 |
| CPT1A | Carnitine O-palmitoyltransferase 1, liver isoform |
| CTSB | Cathepsin B |
| DDX39B | Spliceosome RNA helicase DDX39B |
| DPP7 | Dipeptidyl peptidase 2 |
| DPYSL2 | Dihydropyrimidinase-related protein 2 |
| DYNC1H1 | Cytoplasmic dynein 1 heavy chain 1 |
| EEF1D | Isoform 2 of Elongation factor 1-delta |
| EEF1G | Elongation factor 1-gamma |
| EFTUD2 | 116 kDa U5 small nuclear ribonucleoprotein component |
| EHD2 | EH domain-containing protein 2 |
| EHD4 | EH domain-containing protein 4 |
| EIF3A | Eukaryotic translation initiation factor 3 subunit A |
| EIF3B | Isoform 2 of Eukaryotic translation initiation factor 3 subunit B |
| EIF3CL | Eukaryotic translation initiation factor 3 subunit CL |
| EIF3E | Eukaryotic translation initiation factor 3 subunit E |
| EIF3I | Eukaryotic translation initiation factor 3 subunit I |
| EIF3L | Eukaryotic translation initiation factor 3 subunit L |
| EIF4A3 | Eukaryotic initiation factor 4A-III |
| EIF4B | Eukaryotic translation initiation factor 4B |
| ENO1 | Alpha-enolase |
| ERLIN1 | Erlin-1 |
| ESYT1 | Extended synaptotagmin-1 |
| FERMT3 | Fermitin family homolog 3 |
| FLII | Protein flightless-1 homolog |
| FSCN1 | Fascin |
| GDI2 | Rab GDP dissociation inhibitor beta |
| GLG1 | Isoform 3 of Golgi apparatus protein 1 |
| GNB1 | Guanine nucleotide-binding protein G(I)/G(S)/G(T) subunit beta-1 |
| GNB2 | Guanine nucleotide-binding protein G(I)/G(S)/G(T) subunit beta-2 |
| H2AX | Histone H2AX |
| H2BC4 | Histone H2B type 1-C/E/F/G/I |
| HDLBP | High Density Lipoprotein Binding Protein |
| HP1BP3 | Heterochromatin protein 1-binding protein 3 |
| HSP90AA1 | Heat shock protein HSP 90-alpha |
| HSPA9 | Stress-70 protein, mitochondrial |
| IFI16 | Gamma-interferon-inducible protein 16 |
| IGKV2D-24 | Probable non-functional immunoglobulin kappa variable 2D-24 |
| IGKV2D-29 | Immunoglobulin kappa variable 2D-29 |
| IQGAP1 | Ras GTPase-activating-like protein IQGAP1 |
| ITGB1 | Integrin beta-1 |
| JAK1 | Tyrosine-protein kinase JAK1 |

| | |
|---|---|
| KCTD5 | BTB/POZ domain-containing protein KCTD5 |
| KIF11 | Kinesin-like protein KIF11 |
| KPNB1 | Importin subunit beta-1 |
| KRT18 | Keratin, type I cytoskeletal 18 |
| KRT2 | Keratin, type II cytoskeletal 2 |
| KRT7 | Keratin, type II cytoskeletal 7 |
| KTN1 | Kinectin |
| LCP1 | Plastin-2 |
| LDHB | L-lactate dehydrogenase B chain |
| LMAN1 | Protein ERGIC-53 |
| LMNB2 | Lamin-B2 |
| LRPPRC | Leucine-rich PPR motif-containing protein, mitochondrial |
| MAP1B | Microtubule-associated protein 1B |
| MSN | Moesin |
| MTDH | Protein LYRIC |
| MYH14 | Myosin-14 |
| MYL6 | Myosin Light Chain 6 |
| MYOF | Myoferlin |
| NT5E | 5'-nucleotidase |
| PA2G4 | Proliferation-associated protein 2G4 |
| PCBP1 | Poly(rC)-binding protein 1 |
| PCBP2 | Poly(rC)-binding protein 2 |
| PDIA3 | Protein disulfide-isomerase A3 |
| PGD | 6-phosphogluconate dehydrogenase, decarboxylating |
| PKM | Pyruvate kinase PKM |
| PLOD2 | Isoform 2 of Procollagen-lysine,2-oxoglutarate 5-dioxygenase 2 |
| PPM1B | Protein phosphatase 1B |
| PRMT5 | Protein arginine N-methyltransferase 5 |
| PRPF31 | U4/U6 small nuclear ribonucleoprotein Prp31 |
| PRPF8 | Pre-mRNA-processing-splicing factor 8 |
| RAB7A | Ras-related protein Rab-7a |
| RALY | RNA-binding protein Raly |
| RAN | GTP-binding nuclear protein Ran |
| RHOC | Rho-related GTP-binding protein RhoC |
| RNH1 | Ribonuclease inhibitor |
| ROCK1 | Rho-associated protein kinase 1 |
| RPL10A | 60S ribosomal protein L10a |
| RPL11 | 60S ribosomal protein L11 |
| RPL13 | 60S ribosomal protein L13 |
| RPL14 | 60S ribosomal protein L14 |
| RPL17 | 60S ribosomal protein L17 |
| RPL23 | 60S ribosomal protein L23 |
| RPL29 | 60S ribosomal protein L29 |
| RPL3 | 60S ribosomal protein L3 |
| RPL5 | 60S ribosomal protein L5 |
| RPL8 | 60S ribosomal protein L8 |
| RPN1 | Dolichyl-diphosphooligosaccharide--protein glycosyltransferase subunit 1 |
| RPS16 | 40S ribosomal protein S16 |
| RPS19 | 40S ribosomal protein S19 |
| RPS2 | 40S ribosomal protein S2 |
| RPS26 | 40S ribosomal protein S26 |
| RPS3 | 40S ribosomal protein S3 |
| RPS3A | 40S ribosomal protein S3a |
| RPS4X | 40S ribosomal protein S4, X isoform |
| RPS7 | 40S ribosomal protein S7 |
| RPSA | 40S ribosomal protein SA |
| RRBP1 | Ribosome-binding protein 1 |
| SERBP1 | Plasminogen activator inhibitor 1 RNA-binding protein |
| SERPINE1 | Plasminogen activator inhibitor 1 |
| SERPINH1 | Serpin H1 |
| SND1 | Staphylococcal nuclease domain-containing protein 1 |
| SNRNP200 | U5 small nuclear ribonucleoprotein 200 kDa helicase |
| SRSF1 | Serine/arginine-rich splicing factor 1 |
| SRSF3 | Serine/arginine-rich splicing factor 3 |
| SRSF6 | Serine/arginine-rich splicing factor 6 |
| SRSF7 | Isoform 2 of Serine/arginine-rich splicing factor 7 |
| STK38 | Serine/threonine-protein kinase 38 |
| SUN2 | Isoform 2 of SUN domain-containing protein 2 |
| SUPT16H | FACT complex subunit SPT16 |
| TAB1 | TGF-beta-activated kinase 1 and MAP3K7-binding protein 1 |
| TAF4 | Transcription initiation factor TFIID subunit 4 |
| TAGLN2 | Transgelin-2 |
| TALDO1 | Transaldolase 1 |
| TCP1 | T-complex protein 1 subunit alpha |

| | |
|---|---|
| TGM2 | Protein-glutamine gamma-glutamyltransferase 2 |
| THRAP3 | Thyroid hormone receptor-associated protein 3 |
| TLN1 | Talin-1 |
| TPI1 | Isoform 2 of Triosephosphate isomerase |
| TPP1 | Tripeptidyl-peptidase 1 |
| TRA2B | Transformer-2 protein homolog beta |
| TRIM21 | E3 ubiquitin-protein ligase TRIM21 |
| TUBA1A | Tubulin alpha-1A chain |
| TUBB4A | Tubulin beta-4A chain |
| TXNDC5 | Thioredoxin domain-containing protein 5 |
| UBA52 | Ubiquitin A-52 Residue Ribosomal Protein Fusion Product 1 |
| VAT1 | Ubiquitin-like modifier-activating enzyme 1 |
| WARS1 | Tryptophan--tRNA ligase, cytoplasmic |
| WDR1 | WD repeat-containing protein 1 |
| WDR77 | Methylosome protein 50 |
| WDR93 | WD repeat-containing protein 93 |
| YWHAZ | 14-3-3 protein zeta/delta |

**S12: Grouping of the identified 3F-TG2-associated proteins in the absence (untreated) or presence of inhibitors (NC9 or GTPγS) from HUVEC using anti-Flag M2 agarose resin for the CoIP**

| Only in untreated | Only with NC9 | Only with GTPγS | In both of untreated and with NC9 | In both of untreated and with GTPγS | Both with GTPγS and NC9 | Detected in the case of each experimental setting (untreated, GTPγS, NC9) |
|---|---|---|---|---|---|---|
| ACTR3 | CHCHD3 | CCT6A | ABCD1 | ALB | CCT7 | ABLIM1 |
| AKAP2 | CSE1L | DPYSL2 | ABCD3 | ALYREF | DDX39B | ACAP2 |
| AP2A2 | DKC1 | PGD | ANO10 | CTSB | EIF3CL | ACIN1 |
| APMAP | DPY19L1 | WDR93 | AP2A1 | EHD2 | EIF3E | ACTN1 |
| ATP6AP1 | ENG | | AP2B1 | ESYT1 | H2AX | ACTR1A |
| ATP6V1E1 | FBL | | APOA1 | RPS16 | LCP1 | ADRM1 |
| BST1 | FUT8 | | ARCN1 | RPS2 | TAGLN2 | AHNAK |
| CALD1 | GORASP2 | | ATAD3A | SERBP1 | WDR1 | ALDOA |
| CALU | HMGA2 | | ATP13A1 | SERPINH1 | | ANKFY1 |
| CANX | ITGB4 | | ATP1A1 | SRSF3 | | ANPEP |
| CCT4 | KRT14 | | ATP2A2 | TPP1 | | ANXA5 |
| CDC25C | NACA | | ATP5F1C | | | ANXA6 |
| CDC42BPB | PDCD6IP | | ATP6V0A1 | | | BCLAF1 |
| CDC5L | PGAM5 | | ATP6V0A2 | | | CALR |
| CHTOP | PRPF6 | | ATP6V0D1 | | | CAP1 |
| CMAS | RAI14 | | ATP6V1B2 | | | CAPZA1 |
| COL5A2 | RAP1A | | C3 | | | CAVIN1 |
| COPS3 | RBBP4 | | CAPZB | | | CCT2 |
| CSRP1 | RRAS | | CAV1 | | | CCT3 |
| DHCR7 | SF3B2 | | CCAR1 | | | CCT5 |
| DNAJC11 | SLC35B2 | | CD44 | | | CCT8 |
| DNHD1 | SLTM | | CD55 | | | CD59 |
| EFHD2 | SNRPD1 | | COPB2 | | | CDC42 |
| EIF2S1 | VDAC1 | | CTNND1 | | | CKAP4 |
| EIF3M | VDAC2 | | DCTN1 | | | COPG1 |
| GOLGA3 | VPS35 | | DDOST | | | CPT1A |
| HSPA1L | ZDHHC13 | | DDX23 | | | DPP7 |
| IARS1 | | | DHX15 | | | DYNC1H1 |
| IGF2BP1 | | | DLD | | | EEF1D |
| ITGA2 | | | DLST | | | EEF1G |
| KCTD12 | | | EMC1 | | | EFTUD2 |
| KIF5C | | | EMC2 | | | EHD4 |
| KRT16 | | | ENDOD1 | | | EIF3A |
| KRT6A | | | ERLIN2 | | | EIF3B |
| LIMA1 | | | ERMP1 | | | EIF3I |
| LMNB1 | | | FLOT1 | | | EIF3L |
| MAP4 | | | FLOT2 | | | EIF4A3 |
| MCM3 | | | FMNL2 | | | EIF4B |
| MTCH2 | | | FXR1 | | | ENO1 |
| NAP1L1 | | | GANAB | | | ERLIN1 |
| NAPA | | | GLIPR2 | | | FERMT3 |
| PARP1 | | | GNAI2 | | | FLII |
| PHGDH | | | GNG12 | | | FSCN1 |
| PHLDB1 | | | H1-0 | | | GDI2 |
| PHOX2B | | | H2AC6 | | | GLG1 |

| | | | | | | |
|---|---|---|---|---|---|---|
| PICALM | | | HCLS1 | | | GNB1 |
| PIGR | | | ILF2 | | | GNB2 |
| PIK3C2A | | | IMMT | | | H2BC4 |
| PLOD3 | | | ITGA6 | | | HDLBP |
| PPP1CA | | | ITPR3 | | | HP1BP3 |
| PPP1CB | | | KIF5B | | | HSP90AA1 |
| PPP1R12A | | | LBR | | | HSPA9 |
| PPP1R18 | | | LRRC8A | | | IFI16 |
| QARS1 | | | LRRC8C | | | IGKV2D-24 |
| RAB14 | | | MAGT1 | | | IGKV2D-29 |
| RB1CC1 | | | MBOAT7 | | | IQGAP1 |
| RPL10 | | | MCAM | | | ITGB1 |
| RPL21 | | | MGST3 | | | JAK1 |
| RPL23A | | | MLEC | | | KCTD5 |
| RPL26 | | | MYADM | | | KIF11 |
| RPL27A | | | MYH10 | | | KPNB1 |
| RPL31 | | | NES | | | KRT18 |
| RPL34 | | | NNT | | | KRT2 |
| RPL35 | | | NOP2 | | | KRT7 |
| RPL36A | | | NOP56 | | | KTN1 |
| RPL9 | | | NOP58 | | | LDHB |
| RPS14 | | | OCIAD2 | | | LMAN1 |
| S100A11 | | | OSBPL3 | | | LMNB2 |
| SART1 | | | PFKP | | | LRPPRC |
| SF3B3 | | | PHB2 | | | MAP1B |
| SLC25A24 | | | PLAT | | | MSN |
| SMCHD1 | | | PRPS1 | | | MTDH |
| SRP14 | | | PRPSAP1 | | | MYH14 |
| SRRT | | | PTDSS1 | | | MYL6 |
| SURF4 | | | RAB10 | | | MYOF |
| SYNE1 | | | RAB11B | | | NT5E |
| THBS1 | | | RAB5C | | | PA2G4 |
| TMOD3 | | | RAC2 | | | PCBP1 |
| TPM4 | | | RACK1 | | | PCBP2 |
| TRIM28 | | | RALA | | | PDIA3 |
| TUBA1C | | | RBM10 | | | PKM |
| UGGT1 | | | RER1 | | | PLOD2 |
| VARS1 | | | RPL12 | | | PPM1B |
| YBX3 | | | RPL13A | | | PRMT5 |
| YWHAQ | | | RPL18 | | | PRPF31 |
| ZC3HAV1 | | | RPLP0 | | | PRPF8 |
| | | | RPN2 | | | RAB7A |
| | | | RPS25 | | | RALY |
| | | | RSL1D1 | | | RAN |
| | | | SAMM50 | | | RHOC |
| | | | SEC11A | | | RNH1 |
| | | | SEC22B | | | ROCK1 |
| | | | SEC62 | | | RPL10A |
| | | | SF3B1 | | | RPL11 |
| | | | SFXN1 | | | RPL13 |

| | | | | | | |
|---|---|---|---|---|---|---|
| | | | SFXN3 | | | RPL14 |
| | | | SLC12A4 | | | RPL17 |
| | | | SLC25A11 | | | RPL23 |
| | | | SLC2A1 | | | RPL29 |
| | | | SSRP1 | | | RPL3 |
| | | | STING1 | | | RPL5 |
| | | | STOM | | | RPL8 |
| | | | STT3A | | | RPN1 |
| | | | STT3B | | | RPS19 |
| | | | SVIL | | | RPS26 |
| | | | TBL2 | | | RPS3 |
| | | | TCIRG1 | | | RPS3A |
| | | | TICAM2 | | | RPS4X |
| | | | TMED10 | | | RPS7 |
| | | | TMED9 | | | RPSA |
| | | | TMEM43 | | | RRBP1 |
| | | | TMPO | | | SERPINE1 |
| | | | TMTC3 | | | SND1 |
| | | | TRPV2 | | | SNRNP200 |
| | | | UQCRC1 | | | SRSF1 |
| | | | UQCRC2 | | | SRSF6 |
| | | | VAPA | | | SRSF7 |
| | | | VDAC3 | | | STK38 |
| | | | XRCC5 | | | SUN2 |
| | | | YBX1 | | | SUPT16H |
| | | | YES1 | | | TAB1 |
| | | | ZMPSTE24 | | | TAF4 |
| | | | | | | TALDO1 |
| | | | | | | TCP1 |
| | | | | | | TGM2 |
| | | | | | | THRAP3 |
| | | | | | | TLN1 |
| | | | | | | TPI1 |
| | | | | | | TRA2B |
| | | | | | | TRIM21 |
| | | | | | | TUBA1A |
| | | | | | | TUBB4A |
| | | | | | | TXNDC5 |
| | | | | | | UBA52 |
| | | | | | | VAT1 |
| | | | | | | WARS1 |
| | | | | | | WDR77 |
| | | | | | | YWHAZ |

**S13: The most enriched GO Molecular Functions of the identified 3F-TG2-associated proteins in the absence (untreated) or presence of inhibitors (NC9 or GTPγS) from HUVEC using anti-Flag M2 agarose resin for the CoIP**

| | | | |
|---|---|---|---|
| **UTR** | | | |
| #term ID | term description | false discovery rate | matching proteins in your network |
| GO:0003723 | RNA binding | 7.76e-09 | YBX3,ZC3HAV1,CANX,TRIM28,EIF2S1,RPL35,SRP14,IGF2BP1,SF3B3,SART1,RPL27A,RPL21,MAP4,PARP1,SYNE1,CSRP1,CHTOP,CDC5L,IARS1,KCTD12,CCT4,RPL34,RPS14,RPL31,RPL23A,RPL36A,RPL10,RPL26,NAP1L1,SRRT,RPL9 |
| GO:0005198 | Structural molecule activity | 4.69e-08 | RPL35,THBS1,LMNB1,ACTR3,KRT16,RPL27A,RPL21,MAP4,CSRP1,COL5A2,KRT6A,RPL34,RPS14,RPL31,RPL23A,RPL36A,RPL10,TUBA1C,RPL26,TPM4,RPL9 |
| GO:0003735 | Structural constituent of ribosome | 5.79e-07 | RPL35,RPL27A,RPL21,RPL34,RPS14,RPL31,RPL23A,RPL36A,RPL10,RPL26,RPL9 |
| GO:0045296 | Cadherin binding | 0.00036 | GOLGA3,ZC3HAV1,S100A11,TMOD3,PPP1CA,CALD1,EFHD2,PICALM,RPL34,LIMA1,RPL23A |
| GO:0097159 | Organic cyclic compound binding | 0.00036 | PLOD3,PHOX2B,YBX3,ZC3HAV1,CANX,TRIM28,DNHD1,EIF2S1,RPL35,LMNB1,ACTR3,PIK3C2A,SRP14,IGF2BP1,SF3B3,QARS1,SART1,SMCHD1,RPL27A,RPL21,DHCR7,MAP4,CDC42BPB,PARP1,SYNE1,CSRP1,CHTOP,CDC5L,RAB14,IARS1,HSPA1L,VARS1,KCTD12,CCT4,RPL34,RPS14,RPL31,RPL23A,KIF5C,RPL36A,RPL10,EIF3M,TUBA1C,RPL26,NAP1L1,SRRT,MCM3,PHGDH,RPL9 |
| GO:1901363 | Heterocyclic compound binding | 0.00036 | PLOD3,PHOX2B,YBX3,ZC3HAV1,CANX,TRIM28,DNHD1,EIF2S1,RPL35,LMNB1,ACTR3,PIK3C2A,SRP14,IGF2BP1,SF3B3,QARS1,SART1,SMCHD1,RPL27A,RPL21,DHCR7,MAP4,CDC42BPB,PARP1,SYNE1,CSRP1,CHTOP,CDC5L,RAB14,IARS1,HSPA1L,VARS1,KCTD12,CCT4,RPL34,RPS14,RPL31,RPL23A,KIF5C,RPL36A,RPL10,EIF3M,TUBA1C,RPL26,NAP1L1,SRRT,MCM3,PHGDH,RPL9 |
| GO:0050839 | Cell adhesion molecule binding | 0.00078 | GOLGA3,ZC3HAV1,THBS1,S100A11,ITGA2,TMOD3,PPP1CA,CALD1,EFHD2,PICALM,RPL34,LIMA1,RPL23A |
| GO:0003676 | Nucleic acid binding | 0.0030 | PHOX2B,YBX3,ZC3HAV1,CANX,TRIM28,EIF2S1,RPL35,LMNB1,SRP14,IGF2BP1,SF3B3,SART1,SMCHD1,RPL27A,RPL21,MAP4,PARP1,SYNE1,CSRP1,CHTOP,CDC5L,IARS1,KCTD12,CCT4,RPL34,RPS14,RPL31,RPL23A,RPL36A,RPL10,EIF3M,RPL26,NAP1L1,SRRT,MCM3,RPL9 |
| GO:0005488 | Binding | 0.0068 | RB1CC1,GOLGA3,PLOD3,PHOX2B,YBX3,ZC3HAV1,CANX,TRIM28,ATP6V1E1,DNHD1,EIF2S1,UGGT1,RPL35,THBS1,LMNB1,ACTR3,NAPA,PIK3C2A,SRP14,S100A11,IGF2BP1,ITGA2,SF3B3,QARS1,TMOD3,SART1,CDC25C,PPP1CA,SMCHD1,AP2A2,RPL27A,RPL21,DHCR7,MAP4,CALD1,CDC42BPB,PARP1,SYNE1,CSRP1,CHTOP,ATP6AP1,CDC5L,RAB14,COL5A2,IARS1,HSPA1L,VARS1,EFHD2,KCTD12,YWHAQ,PICALM,CCT4,RPL34,LIMA1,PPP1CB,RPS14,RPL31,RPL23A,PPP1R12A,KIF5C,RPL36A,RPL10,EIF3M,CALU,TUBA1C,SLC25A24,RPL26,NAP1L1,PPP1R18,SRRT,MCM3,PHGDH,TPM4,RPL9 |
| GO:0098641 | Cadherin binding involved in cell-cell adhesion | 0.0492 | S100A11,TMOD3,PPP1CA |
| NC9 | | | |
| #term ID | term description | false discovery rate | matching proteins in your network |
| GO:0044877 | Protein-containing complex binding | 0.0180 | KRT14,ITGB4,FBL,RRAS,PRPF6,SNRPD1,RAP1A,ENG,HMGA2,PGAM5 |
| **UTR+NC9** | | | |
| #term ID | term description | false discovery rate | matching proteins in your network |
| GO:0005215 | Transporter activity | 6.76e-06 | UQCRC1,ABCD1,SFXN3,SLC25A11,APOA1,LRRC8A,ATP6V0A1,NNT,TCIRG1,ATP6V1B2,ATP6V0D1,ANO10,TMED10,OSBPL3,SFXN1,ATP6V0A2,TRPV2,ATP5F1C,ATP13A1,MAGT1,ABCD3,LRRC8C,ITPR3,SLC12A4,SLC2A1,VDAC3,ATP2A2,ATP1A1 |
| GO:0015075 | Ion transmembrane transporter activity | 6.76e-06 | UQCRC1,SFXN3,SLC25A11,LRRC8A,ATP6V0A1,NNT,TCIRG1,ATP6V1B2,ATP6V0D1,ANO10,SFXN1,ATP6V0A2,TRPV2,ATP5F1C,ATP13A1,MAGT1,LRRC8C,ITPR3,SLC12A4,SLC2A1,VDAC3,ATP2A2,ATP1A1 |
| GO:0019829 | ATPase-coupled cation transmembrane transporter activity | 6.76e-06 | ATP6V0A1,TCIRG1,ATP6V1B2,ATP6V0D1,ATP6V0A2,ATP13A1,ATP2A2,ATP1A1 |
| GO:0022857 | Transmembrane transporter activity | 6.76e-06 | UQCRC1,ABCD1,SFXN3,SLC25A11,LRRC8A,ATP6V0A1,NNT,TCIRG1,ATP6V1B2,ATP6V0D1,ANO10,TMED10,SFXN1,ATP6V0A2,TRPV2,ATP5F1C,ATP13A1,MAGT1,ABCD3,LRRC8C,ITPR3,SLC12A4,SLC2A1,VDAC3,ATP2A2,ATP1A1 |
| GO:0042626 | ATPase-coupled transmembrane transporter activity | 6.76e-06 | ABCD1,ATP6V0A1,TCIRG1,ATP6V1B2,ATP6V0D1,ATP6V0A2,ATP13A1,ABCD3,ATP2A2,ATP1A1 |
| GO:0015399 | Primary active transmembrane transporter activity | 1.30e-05 | UQCRC1,ABCD1,ATP6V0A1,TCIRG1,ATP6V1B2,ATP6V0D1,ATP6V0A2,ATP13A1,ABCD3,ATP2A2,ATP1A1 |

| | | | |
|---|---|---|---|
| GO:0015318 | Inorganic molecular entity transmembrane transporter activity | 2.47e-05 | UQCRC1,SLC25A11,LRRC8A,ATP6V0A1,NNT,TCIRG1,ATP6V1B2,ATP6V0D1,ANO10,ATP6V0A2,TRPV2,ATP5F1C,ATP13A1,MAGT1,LRRC8C,ITPR3,SLC12A4,VDAC3,ATP2A2,ATP1A1 |
| GO:0008324 | Cation transmembrane transporter activity | 8.21e-05 | UQCRC1,SFXN3,ATP6V0A1,NNT,TCIRG1,ATP6V1B2,ATP6V0D1,ANO10,SFXN1,ATP6V0A2,TRPV2,ATP5F1C,ATP13A1,MAGT1,ITPR3,SLC12A4,ATP2A2,ATP1A1 |
| GO:0045296 | Cadherin binding | 0.00013 | RAB10,TMPO,FMNL2,KIF5B,RAB11B,VAPA,PFKP,NOP56,CTNND1,ITGA6,CAPZB,RACK1,RSL1D1 |
| GO:0003723 | RNA binding | 0.00017 | SLC25A11,ARCN1,NOP58,CCAR1,SSRP1,TBL2,DDX23,RBM10,SF3B1,DHX15,LBR,GANAB,H1-0,ATP5F1C,FXR1,MYH10,RPL12,ILF2,YBX1,NOP56,NOP2,RPL13A,XRCC5,IMMT,RACK1,RPS25,RPL18,RPLP0,RSL1D1 |
| GO:0140657 | ATP-dependent activity | 0.00017 | ABCD1,ATP6V0A1,TCIRG1,ATP6V1B2,ATP6V0D1,KIF5B,DDX23,ATP6V0A2,DHX15,ATP13A1,MYH10,ABCD3,ATAD3A,XRCC5,ATP2A2,ATP1A1 |
| GO:0046961 | Proton-transporting ATPase activity, rotational mechanism | 0.00035 | ATP6V0A1,TCIRG1,ATP6V1B2,ATP6V0D1,ATP6V0A2 |
| GO:0022890 | Inorganic cation transmembrane transporter activity | 0.00037 | UQCRC1,ATP6V0A1,NNT,TCIRG1,ATP6V1B2,ATP6V0D1,ANO10,ATP6V0A2,TRPV2,ATP5F1C,ATP13A1,MAGT1,ITPR3,SLC12A4,ATP2A2,ATP1A1 |
| GO:0022853 | Active ion transmembrane transporter activity | 0.00043 | UQCRC1,SLC25A11,ATP6V0A1,TCIRG1,ATP6V1B2,ATP6V0D1,ATP6V0A2,ATP13A1,SLC12A4,ATP2A2,ATP1A1 |
| GO:0015078 | Proton transmembrane transporter activity | 0.00049 | UQCRC1,ATP6V0A1,NNT,TCIRG1,ATP6V1B2,ATP6V0D1,ATP6V0A2,ATP5F1C |
| GO:0022804 | Active transmembrane transporter activity | 0.00073 | UQCRC1,ABCD1,SLC25A11,ATP6V0A1,TCIRG1,ATP6V1B2,ATP6V0D1,ATP6V0A2,ATP13A1,ABCD3,SLC12A4,ATP2A2,ATP1A1 |
| GO:0017111 | Nucleoside-triphosphatase activity | 0.0036 | RALA,ABCD1,RAC2,RAB10,KIF5B,DDX23,GNAI2,RAB11B,DHX15,ATP13A1,ABCD3,ATAD3A,ATP2A2,ATP1A1,RAB5C |
| GO:0051117 | ATPase binding | 0.0056 | RALA,NOP58,ATP6V0A1,TCIRG1,ATP6V0A2,CAV1 |
| GO:0019899 | Enzyme binding | 0.0057 | RALA,UQCRC1,ABCD1,MLEC,APOA1,RAC2,NOP58,ATP6V0A1,TCIRG1,STOM,FMNL2,TBL2,HCLS1,ATP6V0A2,CAV1,AP2A1,DCTN1,YBX1,FLOT1,NOP56,XRCC5,CTNND1,SLC12A4,SLC2A1,RACK1,ERLIN2,ATP2A2,YES1,STING1 |
| GO:0044877 | Protein-containing complex binding | 0.0074 | UQCRC1,APOA1,RPN2,NOP58,SSRP1,FMNL2,ATP6V0D1,KIF5B,GNAI2,HCLS1,CAV1,H1-0,SVIL,MYH10,NES,PFKP,XRCC5,ITGA6,CD44,CAPZB,RACK1 |
| **UTR+NC9+GTPγS** | | | |
| #term ID | term description | false discovery rate | matching proteins in your network |
| GO:0003723 | RNA binding | 2.75e-34 | SUPT16H,SRSF6,RALY,TRIM21,SRSF1,LRPPRC,RPL8,ACIN1,RPS3,CCT5,KPNB1,RPL29,CCT3,RPN1,HSPA9,PDIA3,PA2G4,PCBP1,RPL13,SNRNP200,TCP1,PKM,CALR,PRPF31,SRSF7,HSP90AA1,MTDH,RPL3,RPS3A,THRAP3,SND1,RPS26,DYNC1H1,CAVIN1,PCBP2,MSN,EIF3B,IFI16,EIF3A,RPL5,EIF3I,RPS4X,RPL10A,RRBP1,AHNAK,CKAP4,GDI2,FSCN1,KRT18,HDLBP,ACTN1,KTN1,YWHAZ,RPL14,EIF4B,RPSA,EFTUD2,TRA2B,RPL23,BCLAF1,RAN,PRPF8,RPL17,RPS19,RPS7,ENO1,ALDOA,RPL11,EIF4A3,EIF3L |
| GO:0097159 | Organic cyclic compound binding | 9.13e-19 | SUPT16H,EHD4,STK38,WDR77,SRSF6,RALY,TAF4,TRIM21,NT5E,SRSF1,LRPPRC,KIF11,RPL8,ACIN1,TUBB4A,RAB7A,RPS3,CCT5,PLOD2,RHOC,CCT8,KPNB1,RPL29,CCT3,RPN1,HSPA9,CCT2,PDIA3,PA2G4,PCBP1,RPL13,SNRNP200,TCP1,PRMT5,PKM,CALR,H2BC4,PRPF31,SRSF7,EEF1G,HSP90AA1,MTDH,RPL3,RPS3A,ANXA6,THRAP3,SND1,WARS1,RPS26,DYNC1H1,CAVIN1,PCBP2,MSN,EIF3B,TGM2,IFI16,EIF3A,ACTR1A,RPL5,EIF3I,RPS4X,RPL10A,RRBP1,AHNAK,CKAP4,GDI2,FSCN1,KRT18,HDLBP,ACTN1,KTN1,YWHAZ,RPL14,ROCK1,EIF4B,RPSA,EFTUD2,HP1BP3,EEF1D,ERLIN1,TRA2B,RPL23,BCLAF1,TUBA1A,RAN,PRPF8,RPL17,RPS19,MYH14,RPS7,ENO1,ALDOA,RPL11,EIF4A3,CDC42,EIF3L,JAK1 |
| GO:1901363 | Heterocyclic compound binding | 1.28e-18 | SUPT16H,EHD4,STK38,WDR77,SRSF6,RALY,TAF4,TRIM21,NT5E,SRSF1,LRPPRC,KIF11,RPL8,ACIN1,TUBB4A,RAB7A,RPS3,CCT5,PLOD2,RHOC,CCT8,KPNB1,RPL29,CCT3,RPN1,HSPA9,CCT2,PDIA3,PA2G4,PCBP1,RPL13,SNRNP200,TCP1,PRMT5,PKM,CALR,H2BC4,PRPF31,SRSF7,EEF1G,HSP90AA1,MTDH,RPL3,RPS3A,ANXA6,THRAP3,SND1,WARS1,RPS26,DYNC1H1,CAVIN1,PCBP2,MSN,EIF3B,TGM2,IFI16,EIF3A,ACTR1A,RPL5,EIF3I,RPS4X,RPL10A,RRBP1,AHNAK,CKAP4,GDI2,FSCN1,KRT18,HDLBP,ACTN1,KTN1,YWHAZ,RPL14,ROCK1,EIF4B,RPSA,EFTUD2,HP1BP3,EEF1D,TRA2B,RPL23,BCLAF1,TUBA1A,RAN,PRPF8,RPL17,RPS19,MYH14,RPS7,ENO1,ALDOA,RPL11,EIF4A3,CDC42,EIF3L,JAK1 |

| | | | |
|---|---|---|---|
| GO:0003676 | Nucleic acid binding | 6.17e-18 | SUPT16H,EHD4,WDR77,SRSF6,RALY,TAF4,TRIM21,SRSF1,LRPPRC,RPL8,ACIN1,RPS3,CCT5,KPNB1,RPL29,CCT3,RPN1,HSPA9,PDIA3,PA2G4,PCBP1,RPL13,SNRNP200,TCP1,PRMT5,PKM,CALR,H2BC4,PRPF31,SRSF7,EEF1G,HSP90AA1,MTDH,RPL3,RPS3A,THRAP3,SND1,RPS26,DYNC1H1,CAVIN1,PCBP2,MSN,EIF3B,IFI16,EIF3A,RPL5,EIF3I,RPS4X,RPL10A,RRBP1,AHNAK,CKAP4,GDI2,FSCN1,KRT18,HDLBP,ACTN1,KTN1,YWHAZ,RPL14,EIF4B,RPSA,EFTUD2,HP1BP3,EEF1D,TRA2B,RPL23,BCLAF1,RAN,PRPF8,RPL17,RPS19,RPS7,ENO1,ALDOA,RPL11,EIF4A3,EIF3L |
| GO:0005198 | Structural molecule activity | 7.36e-15 | RPL8,TUBB4A,RPS3,RPL29,MAP1B,RPL13,KRT2,TLN1,H2BC4,COPG1,LMNB2,KRT7,RPL3,RPS3A,RPS26,MSN,EIF3A,RPL5,RPS4X,RPL10A,AHNAK,KRT18,ACTN1,RPL14,UBA52,RPSA,HP1BP3,RPL23,TUBA1A,MYL6,RPL17,RPS19,RPS7,RPL11 |
| GO:0005488 | Binding | 1.02e-14 | GLG1,TAB1,SUPT16H,EHD4,SERPINE1,TPI1,STK38,WDR77,SRSF6,RALY,LMAN1,TAF4,TRIM21,NT5E,SRSF1,LRPPRC,KIF11,RPL8,ACIN1,CAPZA1,TUBB4A,RAB7A,CPT1A,IQGAP1,ABLIM1,RPS3,FERMT3,CCT5,PPM1B,PLOD2,RHOC,CCT8,KPNB1,RPL29,CCT3,RPN1,ANXA5,MAP1B,HSPA9,CCT2,ANPEP,PDIA3,KCTD5,PA2G4,GNB2,PCBP1,RPL13,KRT2,TLN1,SNRNP200,TCP1,PRMT5,PKM,CALR,TALDO1,H2BC4,PRPF31,ACAP2,FLII,SRSF7,LMNB2,EEF1G,HSP90AA1,MTDH,RPL3,RPS3A,ANXA6,THRAP3,SND1,WARS1,VAT1,RPS26,DYNC1H1,CAVIN1,MYOF,PCBP2,MSN,EIF3B,TGM2,IFI16,EIF3A,ACTR1A,RPL5,CAP1,EIF3I,RPS4X,RPL10A,RRBP1,AHNAK,CKAP4,GNB1,GDI2,FSCN1,KRT18,HDLBP,ACTN1,KTN1,YWHAZ,ITGB1,RPL14,ROCK1,SUN2,UBA52,EIF4B,RPSA,EFTUD2,HP1BP3,EEF1D,ERLIN1,TRA2B,RPL23,BCLAF1,TUBA1A,RAN,MYL6,ANKFY1,PRPF8,RPL17,RPS19,ADRM1,MYH14,RPS7,ENO1,ALDOA,RPL11,EIF4A3,CDC42,CD59,EIF3L,JAK1 |
| GO:0045296 | Cadherin binding | 2.22e-14 | EHD4,STK38,CAPZA1,IQGAP1,CCT8,RPL29,PCBP1,TLN1,PKM,EEF1G,SND1,RPS26,AHNAK,FSCN1,KRT18,HDLBP,KTN1,YWHAZ,ITGB1,RPL14,EEF1D,RAN,ENO1,ALDOA |
| GO:0050839 | Cell adhesion molecule binding | 2.76e-14 | EHD4,STK38,CAPZA1,IQGAP1,FERMT3,CCT8,RPL29,PCBP1,TLN1,PKM,CALR,EEF1G,SND1,RPS26,MSN,AHNAK,FSCN1,KRT18,HDLBP,ACTN1,KTN1,YWHAZ,ITGB1,RPL14,RPSA,EEF1D,RAN,ENO1,ALDOA |
| GO:0003735 | Structural constituent of ribosome | 3.84e-13 | RPL8,RPS3,RPL29,RPL13,RPL3,RPS3A,RPS26,RPL5,RPS4X,RPL10A,RPL14,UBA52,RPSA,RPL23,RPL17,RPS19,RPS7,RPL11 |
| GO:0005515 | Protein binding | 3.10e-12 | GLG1,TAB1,EHD4,SERPINE1,TPI1,STK38,LMAN1,TAF4,TRIM21,SRSF1,LRPPRC,KIF11,ACIN1,CAPZA1,RAB7A,CPT1A,IQGAP1,ABLIM1,RPS3,FERMT3,CCT5,RHOC,CCT8,KPNB1,RPL29,CCT3,MAP1B,HSPA9,CCT2,PDIA3,KCTD5,PA2G4,GNB2,PCBP1,KRT2,TLN1,SNRNP200,TCP1,PRMT5,PKM,CALR,H2BC4,PRPF31,FLII,SRSF7,LMNB2,EEF1G,HSP90AA1,MTDH,ANXA6,THRAP3,SND1,WARS1,RPS26,DYNC1H1,CAVIN1,PCBP2,MSN,EIF3B,IFI16,RPL5,CAP1,AHNAK,GNB1,GDI2,FSCN1,KRT18,HDLBP,ACTN1,KTN1,YWHAZ,ITGB1,RPL14,ROCK1,SUN2,UBA52,RPSA,EEF1D,ERLIN1,TRA2B,RPL23,TUBA1A,RAN,ANKFY1,PRPF8,RPS19,ADRM1,MYH14,RPS7,ENO1,ALDOA,RPL11,CDC42,CD59,JAK1 |
| GO:0003729 | mRNA binding | 1.21e-09 | SRSF6,SRSF1,LRPPRC,RPS3,CCT5,PCBP1,PKM,CALR,HSP90AA1,RPS3A,RPS26,PCBP2,EIF3A,RPL5,HDLBP,TRA2B,BCLAF1,RPS7,EIF4A3 |
| GO:0019899 | Enzyme binding | 4.74e-08 | TAB1,SERPINE1,TPI1,STK38,TRIM21,SRSF1,LRPPRC,KIF11,ACIN1,RAB7A,IQGAP1,RPS3,RHOC,KPNB1,HSPA9,CCT2,PA2G4,GNB2,TCP1,CALR,HSP90AA1,WARS1,PCBP2,MSN,RPL5,CAP1,GNB1,GDI2,YWHAZ,ITGB1,ROCK1,UBA52,ERLIN1,RPL23,ANKFY1,RPS19,ADRM1,RPS7,ENO1,RPL11,CDC42,JAK1 |
| GO:0031625 | Ubiquitin protein ligase binding | 1.99e-07 | TPI1,LRPPRC,HSPA9,CCT2,PA2G4,TCP1,CALR,HSP90AA1,PCBP2,RPL5,YWHAZ,UBA52,ERLIN1,RPL23,RPL11,JAK1 |
| GO:0044877 | Protein-containing complex binding | 8.74e-07 | TAB1,SUPT16H,CAPZA1,RAB7A,IQGAP1,ABLIM1,RPS3,FERMT3,MAP1B,KCTD5,GNB2,TLN1,PRMT5,PKM,CALR,PRPF31,FLII,HSP90AA1,ANXA6,SND1,GNB1,FSCN1,ACTN1,ITGB1,EIF4B,RPSA,HP1BP3,ADRM1,MYH14,EIF4A3 |
| GO:0044183 | Protein folding chaperone | 2.65e-06 | CCT5,CCT8,CCT3,HSPA9,CCT2,TCP1,CALR,HSP90AA1 |
| GO:0019843 | rRNA binding | 1.32e-05 | RPL8,RPS3,RPL3,CAVIN1,RPL5,RPS4X,RPL23,RPL11 |
| GO:0051082 | Unfolded protein binding | 4.97e-05 | LMAN1,CCT5,CCT8,CCT3,HSPA9,CCT2,TCP1,CALR,HSP90AA1 |
| GO:0140662 | ATP-dependent protein folding chaperone | 4.97e-05 | CCT5,CCT8,CCT3,HSPA9,CCT2,TCP1 |
| GO:0008092 | Cytoskeletal protein binding | 0.00012 | LRPPRC,KIF11,CAPZA1,IQGAP1,ABLIM1,RPS3,CCT5,MAP1B,KRT2,TLN1,FLII,HSP90AA1,ANXA6,MSN,CAP1,FSCN1,ACTN1,KTN1,ITGB1,ROCK1,SUN2,MYH14,ALDOA |
| GO:1990948 | Ubiquitin ligase inhibitor activity | 0.00034 | RPL5,RPL23,RPS7,RPL11 |

**S14: Oligonucleotides used in the study**

| **Manufacturer** | **Name** | **Sequence (5’-3’)** |
|---|---|---|
| Sigma-Aldrich (primers for BAP insertion) | N-BAPINSfor | [Phos]ct cag aaa atc gaa tgg cac gaa ggt atg aaa gaa acc gct g |
| | N-BAPINSrev | [Phos]c ctc gaa gat gtc gtt cag acc aga aga atg atg atg |
| Sigma-Aldrich (primers for 3xFlag insertion) | 3xflagINS_for | [Phos]GGTGATTATAAAGATCATGACATCGATTACAAGGACGATGAC |
| | 3xflagINS rev | [Phos]GTCATGGTCTTTGTAGTCCATGGTGGAGCCTGCTTT |
| Sigma-Aldrich (primers for silencing) | TRCN0000272816 (shTG2)For | CCGGCCACCCACCATATTGTTTGATCTCGAGATCAAACAATATGGTGGGTGGTTTTTG |
| | TRCN0000272816 (shTG2)Rev | AATTCAAAAACCACCCACCATATTGTTTGATCTCGAGATCAAACAATATGGTGGGTGG |
| Sigma-Aldrich (primers for scramble) | scramble For | AATTCCTAAGGTTAAGTCGCCCTCGCTCGAGCGAGGGCGACTTAACCTTAGGTTTTTTTAT |
| | scramble Rev | AAAAAAACCTAAGGTTAAGTCGCCCTCGCTCGAGCGAGGGCGACTTAACCTTAGG |

**S15: Coding sequence of pET-30 Ek/LIC BAP rhTG2 vector**

ATGCACCATCATCATCATCATtcttctGGTCTGaacgacatcttcgaggctcagAAAatcgaatggcacga
aggtatgaaagaaaccgctgctgctaaattcgaacgccagcacatggacagcccagatctgggtaccgatgacgacgacaagatga
gaattcagacc**ATG**gccgaggagctggtcttagagaggtgtgatctggagctggagaccaatggccgagaccaccacacggccg
acctgtgccgggagaagctggtggtgcgacggggccagcccttctggctgaccctgcactttgagggccgcaactacgaggccagt
gtagacagtctcaccttcagtgtcgtgaccggcccagcccctagccaggaggccgggaccaaggcccgttttccactaagagatgct
gtggaggagggtgactggacagccaccgtggtggaccagcaagactgcaccctctcgctgcagctcaccaccccggccaacgccc
ccatcggcctgtatcgcctcagcctggaggcctccactggctaccagggatccagctttgtgctgggccacttcattttgctcttcaacgc
ctggtgcccagcggatgctgtgtacctggactcggaagaggagcggcaggagtatgtcctcacccagcagggctttatctaccaggg
ctcggccaagttcatcaagaacataccttggaattttgggcagtttgaagatgggatcctagacatctgcctgatccttctagatgtcaac
cccaagttcctgaagaacgccggccgtgactgctcccgccgcagcagccccgtctacgtgggccgggtggttagtggcatggtcaac
tgcaacgatgaccagggtgtgctgctgggacgctgggacaacaactacggggacggcgtcagccccatgtcctggatcggcagcgt
ggacatcctgcggcgctggaagaaccacggctgccagcgcgtcaagtatggccagtgctgggtcttcgccgccgtggcctgcacag
tgctgaggtgcctaggcatccctacccgcgtcgtgaccaactacaactcggcccatgaccagaacagcaaccttctcatcgagtacttc
cgcaatgagtttggggagatccagggtgacaagagcgagatgatctggaacttccactgctgggtggagtcgtggatgaccaggccg
gacctgcagccggggtacgagggctggcaggccctggacccaacgccccaggagaagagcgaagggacgtactgctgtggccca
gttccagttcgtgccatcaaggagggcgacctgagcaccaagtacgatgcgccctttgtctttgcggaggtcaatgccgacgtggtag
actggatccagcaggacgatgggtctgtgcacaaatccatcaaccgttccctgatcgttgggctgaagatcagcactaagagcgtggg
ccgagacgagcgggaggatatcacccacacctacaaatacccagaggggtcctcagaggagagggaggccttcacaagggcgaa
ccacctgaacaaactggccgagaaggaggagacagggatggccatgcggatccgtgtgggccagagcatgaacatgggcagtga
ctttgacgtctttgcccacatcaccaacaacaccgctgaggagtacgtctgccgcctcctgctctgtgcccgcaccgtcagctacaatg
ggatcttgggggcccgagtgtggcaccaagtacctgctcaacctcaacctggagcctttctctgagaagagcgttcctctttgcatcctcta
tgagaaataccgtgactgccttacggagtccaacctcatcaaggtgcgggccctcctcgtggagccagttatcaacagctacctgctgg
ctgagagggacctctacctggagaatccagaaatcaagatccggatccttggggagcccaagcagaaacgcaagctggtggctgag
gtgtccctgcagaacccgctccctgtggccctggaaggctgcaccttcactgtggagggggccggcctgactgaggagcagaagac
ggtggagatcccagaccccgtggaggcaggggaggaagttaaggtgagaatggacctgctgccgctccacatgggcctccacaag
ctggtggtgaacttcgagagcgacaagctgaaggctgtgaagggcttccggaatgtcatcattggccccgccTAA

**Amino acid sequence of pET-30 Ek/LIC BAP rhTG2 derived site-specifically biotinylated recombinant TG2 used in coi**

MHHHHHHSSGLNDIFEAQKIEWHEGMKETAAAKFERQHMDSPDLGTDDDDKMRIQT**M**AEELVLERCDLELE
TNGRDHHTADLCREKLVVRRGQPFWLTLHFEGRNYEASVDSLTFSVVTGPAPSQEAGTKARFPLRDAVEEGD
WTATVVDQQDCTLSLQLTTPANAPIGLYRLSLEASTGYQGSSFVLGHFILLFNAWCPADAVYLDSEEERQEYV
LTQQGFIYQGSAKFIKNIPWNFGQFEDGILDICLILLDVNPKFLKNAGRDCSRRSSPVYVGRVVSGMVNCNDD
QGVLLGRWDNNYGDGVSPMSWIGSVDILRRWKNHGCQRVKYGQCWVFAAVACTVLRCLGIPTRVVTNYNS
AHDQNSNLLIEYFRNEFGEIQGDKSEMIWNFHCWVESWMTRPDLQPGYEGWQALDPTPQEKSEGTYCCGPV
PVRAIKEGDLSTKYDAPFVFAEVNADVVDWIQQDDGSVHKSINRSLIVGLKISTKSVGRDEREDITHTYKYPE
GSSEEREAFTRANHLNKLAEKEETGMAMRIRVGQSMNMGSDFDVFAHITNNTAEEYVCRLLLCARTVSYNGI
LGPECGTKYLLNLNLEPFSEKSVPLCILYEKYRDCLTESNLIKVRALLVEPVINSYLLAERDLYLENPEIKIRILGE
PKQKRKLVAEVSLQNPLPVALEGCTFTVEGAGLTEEQKTVEIPDPVEAGEEVKVRMDLLPLHMGLHKLVVNF
ESDKLKAVKGFRNVIIGPA

**S16: Coding sequence of transgene 3xFlag human TG2 vector**

ATGGACTACAAAGACCATGACGGTGATTATAAAGATCATGACATCGATTACAAG
GACGATGACGACaaactcggatccatgggaaccaattcagtcgactggatccggtaccgaattcagacc**ATG**gccgag
gagctggtcttagagaggtgtgatctggagctggagaccaatggccgagaccaccacacggccgacctgtgccgggagaagctggt
ggtgcgacggggccagcccttctggctgaccctgcactttgagggccgcaactacgaggccagtgtagacagtctcaccttcagtgtc
gtgaccggcccagcccctagccaggaggccgggaccaaggcccgttttccactaagagatgctgtggaggagggtgactggacag
ccaccgtggtggaccagcaagactgcaccctctcgctgcagctcaccaccccggccaacgcccccatcggcctgtatcgcctcagcc
tggaggcctccactggctaccagggatccagctttgtgctgggccacttcattttgctcttcaacgcctggtgcccagcggatgctgtgt
acctggactcggaagaggagcggcaggagtatgtcctcacccagcagggctttatctaccagggctcggccaagttcatcaagaaca
taccttggaattttgggcagtttgaagatgggatcctagacatctgcctgatccttctagatgtcaaccccaagttcctgaagaacgccgg
ccgtgactgctcccgccgcagcagccccgtctacgtgggccgggtggttagtggcatggtcaactgcaacgatgaccagggtgtgct
gctgggacgctgggacaacaactacggggacggcgtcagccccatgtcctggatcggcagcgtggacatcctgcggcgctggaag
aaccacggctgccagcgcgtcaagtatggccagtgctgggtcttcgccgccgtggcctgcacagtgctgaggtgcctaggcatccct
acccgcgtcgtgaccaactacaactcggcccatgaccagaacagcaaccttctcatcgagtacttccgcaatgagtttggggagatcc
agggtgacaagagcgagatgatctggaacttccactgctgggtggagtcgtggatgaccaggccggacctgcagccggggtacgag
ggctggcaggccctggacccaacgccccaggagaagagcgaagggacgtactgctgtggcccagttccagttcgtgccatcaagg
agggcgacctgagcaccaagtacgatgcgccctttgtctttgcggaggtcaatgccgacgtggtagactggatccagcaggacgatg
ggtctgtgcacaaatccatcaaccgttccctgatcgttgggctgaagatcagcactaagagcgtgggccgagacgagcgggaggata
tcacccacacctacaaatacccagaggggtcctcagaggagagggaggccttcacaagggcgaaccacctgaacaaactggccga
gaaggaggagacagggatggccatgcggatccgtgtgggccagagcatgaacatgggcagtgactttgacgtctttgcccacatcac
caacaacaccgctgaggagtacgtctgccgcctcctgctctgtgcccgcaccgtcagctacaatgggatcttggggcccgagtgtggc
accaagtacctgctcaacctcaacctggagcctttctctgagaagagcgttcctctttgcatcctctatgagaaataccgtgactgccttac
ggagtccaacctcatcaaggtgcgggccctcctcgtggagccagttatcaacagctacctgctggctgagagggacctctacctggag
aatccagaaatcaagatccggatccttggggagcccaagcagaaacgcaagctggtggctgaggtgtccctgcagaacccgctccct
gtggccctggaaggctgcaccttcactgtggagggggccggcctgactgaggagcagaagacggtggagatcccagaccccgtgg
aggcaggggaggaagttaaggtgagaatggacctgctgccgctccacatgggcctccacaagctggtggtgaacttcgagagcgac
aagctgaaggctgtgaagggcttccggaatgtcatcattggccccgccTAA

**Amino acid sequence of transgene 3xFlag human TG2**

M**DYKDHDGDYKDHDIDYKDDDD**KLGSMGTNSVDWIRYRIQT**M**AEELVLERCDLELETNGRDHHTAD
LCREKLVVRRGQPFWLTLHFEGRNYEASVDSLTFSVVTGPAPSQEAGTKARFPLRDAVEEGDWTATVV
DQQDCTLSLQLTTPANAPIGLYRLSLEASTGYQGSSFVLGHFILLFNAWCPADAVYLDSEEERQEYVLTQ
QGFIYQGSAKFIKNIPWNFGQFEDGILDICLILLDVNPKFLKNAGRDCSRRSSPVYVGRVVSGMVNCNDD
QGVLLGRWDNNYGDGVSPMSWIGSVDILRRWKNHGCQRVKYGQCWVFAAVACTVLRCLGIPTRVVTN
YNSAHDQNSNLLIEYFRNEFGEIQGDKSEMIWNFHCWVESWMTRPDLQPGYEGWQALDPTPQEKSEGT
YCCGPVPVRAIKEGDLSTKYDAPFVFAEVNADVVDWIQQDDGSVHKSINRSLIVGLKISTKSVGRDERED
ITHTYKYPEGSSEEREAFTRANHLNKLAEKEETGMAMRIRVGQSMNMGSDFDVFAHITNNTAEEYVCRL
LLCARTVSYNGILGPECGTKYLLNLNLEPFSEKSVPLCILYEKYRDCLTESNLIKVRALLVEPVINSYLLAE
RDLYLENPEIKIRILGEPKQKRKLVAEVSLQNPLPVALEGCTFTVEGAGLTEEQKTVEIPDPVEAGEEVKV
RMDLLPLHMGLHKLVVNFESDKLKAVKGFRNVIIGPA
